# Chinese fir genome and the evolution of gymnosperms

**DOI:** 10.1101/2022.10.25.513437

**Authors:** Si-Zu Lin, Yu Chen, Chao Wu, Wei-Hong Sun, Zhen Li, Hengchi Chen, Jieyu Wang, Changmian Ji, Shu-Bin Li, Zhiwen Wang, Wen-Chieh Tsai, Xiang-Qing Ma, Si-Ren Lan, Fei-Ping Zhang, Ya-Cong Xie, Lei Yao, Yan Zhang, Meng-Meng Lü, Jia-Jun Zhang, Di-Yang Zhang, Yi-Quan Ye, Xia Yu, Shan-Shan Xu, Zhi-Hui Ma, Guo-Chang Ding, Guang-Qiu Cao, Zong-Ming He, Peng-Fei Wu, Kai-Min Lin, Ai-Qin Liu, Yan-Qing Lin, Shao-Ning Ruan, Bao Liu, Shi-Jiang Cao, Li-Li Zhou, Ming Li, Peng Shuai, Xiao-Long Hou, Yi-Han Wu, Nuo Li, Sheng Xiong, Yang Hao, Zhuang Zhou, Xue-Die Liu, Dan-Dan Zuo, Jia Li, Pei Wang, Jian Zhang, Ding-Kun Liu, Gui-Zhen Chen, Jie Huang, Ming-Zhong Huang, Yuanyuan Li, Qinyao Zheng, Xiang Zhao, Wen-Ying Zhong, Feng-Ling Wang, Xin-Chao Cheng, Yin Yu, Zhi-Wei Liu, Hongkun Zheng, Ray Ming, Yves Van de Peer, Zhong-Jian Liu

## Abstract

Seed plants comprise angiosperms and gymnosperms. The latter includes gnetophytes, cycads, Ginkgo, and conifers. Conifers are distributed worldwide, with 630 species distributed across eight families and 70 genera. Their distinctiveness has triggered much debate on their origin, evolution, and phylogenetic placement among seed plants. To better understand the evolution of gymnosperms and their relation to other seed plants, we report here a high-quality genome sequence for a tree species, Chinese fir (*Cunninghamia lanceolata*), which has excellent timber quality and high aluminum adaptability and is a member of Cupressaceae with high levels of heterozygosity. We assembled an 11.24 Gb genome with a contig N50 value of 2.15 Mb and anchored the 10.89 Gb sequence to 11 chromosomes. Phylogenomic analyses showed that cycads sister to Ginkgo, which place to sister in all gymnosperm lineages, and Gnetales within conifers sister to Pinaceae. Whole-genome duplication (WGD) analysis showed that the ancestor of seed plants has differentiated into angiosperms and gymnosperms after having experienced a WGD event. The ancestor of extant gymnosperm has experienced a gymnosperm-specific WGD event and the extant angiosperms do not share a common WGD before their most recent common ancestor diverged into existing angiosperms lineages. Analysis of the MADS-box gene family of *C. lanceolata* revealed the developmental mechanism of the reproductive organs in *C. lanceolata*, which supported the (A)B(C) model of the development of gymnosperms reproductive organs. In addition, astringent seeds and shedding of whole branches (with withered leaves) might be a strategy of *C. lanceolata* that evolved during long-term adaptation to an aluminum-rich environment. The findings also reveal the molecular regulation mechanism of shade tolerance in *C. lanceolata* seedlings. Our results improve the resolution of ancestral genomic features within seed plants and the knowledge of genome evolution and diversification of gymnosperms.

## Introduction

Seed plants are represented by two distinct lineages: gymnosperms and angiosperms (flowering plants). Gymnosperms are a group of terrestrial plants comprising the extant taxa cycads, Ginkgo, conifers, and gnetophytes^1–3^. Although the species diversity of living gymnosperms is low, with just over 1,000 species compared to the 369,000 angiosperm species, the distribution of gymnosperms on Earth is approximately similar to that of angiosperms. The fossil record indicates considerable seed plant diversity in the late Devonian period, approximately 360 million years ago (Mya)^4^. Gymnosperms appeared approximately 300 Mya, long before the origin of flowering plants^5,6^. Thus, today’s gymnosperms are only a relic of their former diversity, which presents a major challenge in reconstructing the evolutionary relationship between the extant lineages^7^.

The relationships within gymnosperm lineages have been the subject of debate for decades and many hypotheses have been described^8–10^. Conifers are paraphyletic, with gnetophytes as sister to cupressophytes (non-Pinaceae families Araucariaceae, Cupressaceae, Podocarpaceae, Sciadopityaceae, and Taxaceae; ‘Gnecup’ hypothesis) or Pinaceae (‘Gnepine’ hypothesis)^9–11^. Furthermore, gnetophytes are placed as sister to all conifers (‘Gnetifer’ hypothesis) or all other gymnosperms^12,13^.

One way to help infer the correct phylogenetic relationships between extant gymnosperm lineages is to obtain genomes of more gymnosperms. However, gymnosperms have dramatically larger genomes than those of angiosperms, in part because of the large number of repetitive sequences, transposable elements, and gene duplication in their genomes^14–18^. The large size and overall complexity of gymnosperm genomes make it difficult to obtain high-quality genomes. To date, the availability of whole-genome sequences for gymnosperms has been limited to conifers (*Picea abies*, *Picea glauca*, *Pinus taeda*, *Pinus tabuliformis*, *Larix kaempferi*)^16,19–25^, *Ginkgo biloba*^26,27^, *Taxus wallichiana*^28^, *Cycas panzhihuaensis*^29^, and the gnetophytes *Gnetum montanum* and *Welwitschia mirabilis*^12,30^. The genome data for Cupressaceae, an important member of gymnosperms, is still not available.

Conifers dominated forests both before and after the major mass extinction events at the end of the Permian and Cretaceous periods^19^, with approximately 630 species distributed across eight families and 70 genera. Cupressaceae is the only lineage of gymnosperms that are widely distributed in both the northern and southern hemispheres, with 30 genera and 150 species. Many Cupressaceae plants, such as arborvitae, cypress, and juniper, constitute the dominant tree in the forest, as well as are important tree species for afforestation, timber production, and landscaping. The genus *Cunninghamia*, which belongs to Cupressaceae, has only two species, Chinese fir (*C. lanceolata*) and *C. konishii*, which are native to south-central and southeastern China, Laos, and Vietnam^31^. *Cunninghamia lanceolata* is endemic to south-central and southeastern China and northern Vietnam^32^. The main soil types in its distribution area are acid aluminized soils, such as mountain yellow brown soil, red-yellow soil and lateritic red soil (soil pH 4.5–6.4, active aluminum 0.8–2.4 mg/100 g)^33^. The characteristic phenotype of *C. lanceolata* is that the leaves do not fall after withering. There are many astringent seeds in the mature seeds. As early as 8,000 years ago in the Neolithic era, the ancient Yue people in China began to use *C. lanceolata* resources to make tools for production and living^34^. *Cunninghamia lanceolata* has been cultivated in China for more than 2,000 years because of its ecological and economic value^35^. *Cunninghamia lanceolata* plantations account for approximately 21.40% of the area of the artificial arbor forest, and its wood accounts for approximately 30.00% of the plantation wood accumulation. Its area and accumulation are ranked first among the major afforestation tree species in China.

Here, we report a high-quality genome sequence of *C. lanceolata*. The data reveal the phylogenetic relationships within gymnosperms and the origin and distribution of conifers. Comparisons within gymnosperms and angiosperms highlight the unique nature of the Cupressaceae genome, providing new insights into patterns of genome divergence across seed plants.

## Results and discussion

### Genome sequence and assembly

*Cunninghamia lanceolata* is a diploid plant and contains 11 chromosomes (2n = 2× = 22)^36^. Survey analysis indicated that the *C. lanceolata* genome showed a high level of heterozygosity, corresponding to 0.69% of the 10.42 Gb genome size according to *K*-mer analysis (**Supplementary Table 1** and **Supplementary Fig. 1**). For *de novo* whole-genome sequencing of *C. lanceolata*, we obtained 1,113.16 Gb of clean reads with an average length of 12.44 kb using PacBio technology (**Supplementary Table 2**). The final assembled genome was 11.24 Gb with a contig N50 value of 2.16 Mb (**Supplementary Table 3**). Benchmarking Universal Single-Copy Orthologs (BUSCO) assessment showed that the completeness of the gene set of the assembled genome was 50.00% (**Supplementary Table 4**), and the Illumina read alignment rate was 92.83% (**Supplementary Table 5**). The assembled genome integrity of Chinese fir is larger than that of other conifer species, but the assembled genome integrity of conifers is relatively low (**Supplementary Table 4**). BUSCO with a default setting tends to miss genes with long introns because predicted introns in conifers are longer than the maximum intron length in the gene predictor. This results in the low integrity of the assembled genomes of gymnosperms with large genomes. We further performed high-throughput/resolution chromosome conformation capture (Hi-C) technology to obtain a chromosome-level assembly. A total of 10.89 Gb reads (96.90% of assembled genome) were located on 11 chromosomes (**Supplementary Table 6**). The length of the 11 chromosomes ranged from 0.63–1.56 Gb with a scaffold N50 value of 0.93 Gb (**Supplementary Tables 5, 7**). The heat map of the interaction between the chromosomes indicated that the Hi-C assembly of *C. lanceolata* is of high-quality (**Supplementary Fig. 2**).

### Gene prediction and annotation

The *C. lanceolata* genome was predicted to contain 37,225 protein-coding genes (**Supplementary Table 8**). Of these, 34,559 (92.84%) protein-coding genes were functionally annotated (**Supplementary Table 9**). The completeness of the annotated genome was 85.10% based on BUSCO assessment. The value was higher than that of *P. abies* (25.40%), *P. taeda* (23.23%), *Ginkgo* (60.04%), *T. wallichiana* (65.20%), and *G. montanum* (82.84%) (**Supplementary Table 3**). We identified 50 microRNAs (miRNAs), 3,955 transfer RNAs (tRNAs), and 2,930 ribosomal RNAs (rRNA) in the *C. lanceolata* genome (**Supplementary Table 10**). In addition, the important feature of the predicted gene structure of *C. lanceolata* was the presence of numerous long introns. The average intron length of 6,482 bp (**Supplementary Table 11**) is longer than the average intron length in genomes of gymnosperms that have been sequenced^12,16,19–27^. Generally, species with long intron also have large genomes^37^. More long introns may be one reason for the large genome size of *C. lanceolata*.

The larger intron size of Ginkgo and conifers may be because of the long-term stable amplification of long terminal repeat retrotransposons (LTR-RTs)^19,38^. We estimated that 92.31% (10.38 Gb) of the *C. lanceolata* genome consisted of repetitive sequences (**Supplementary Table 12**). Compared with that in other gymnosperms, the proportion of repetitive sequences in the *C. lanceolata* genome was the largest (**Supplementary Table 12**). The repetitive sequences of *C. lanceolata* are relatively abundant, of which LTRs account for 69.92% (Ty3/Gypsy 42.38%; Ty1/ Copia 23.88%), followed by DIRS (8.26%) and LINE (3.26%) (**Supplementary Table 13**). We analyzed the LTR insertion time of gymnosperms and the LTR insertion frequency of *P. abies*, *P. taeda*, *G, montanum*, and Ginkgo. A trend of an initial rise followed by a decline was evident, reaching a peak between 18–14 Mya. However, the insertion frequency of *C. lanceolata* LTR gradually rose beginning abruptly at 3–7 Mya, except for a peak at 18–14 Mya, while that of other gymnosperms maintained a downward trend (**Fig. 1a**). One reason for the larger genomes of gymnosperms is the lack of effective LTR elimination mechanisms^12,19^. We speculate that the abundant LTR retrotransposons, steady accumulation of LTRs, and lack of efficient LTR elimination mechanisms in the *C. lanceolata* genome may explain the large genome size of *C. lanceolata*.

**Figure 1.**
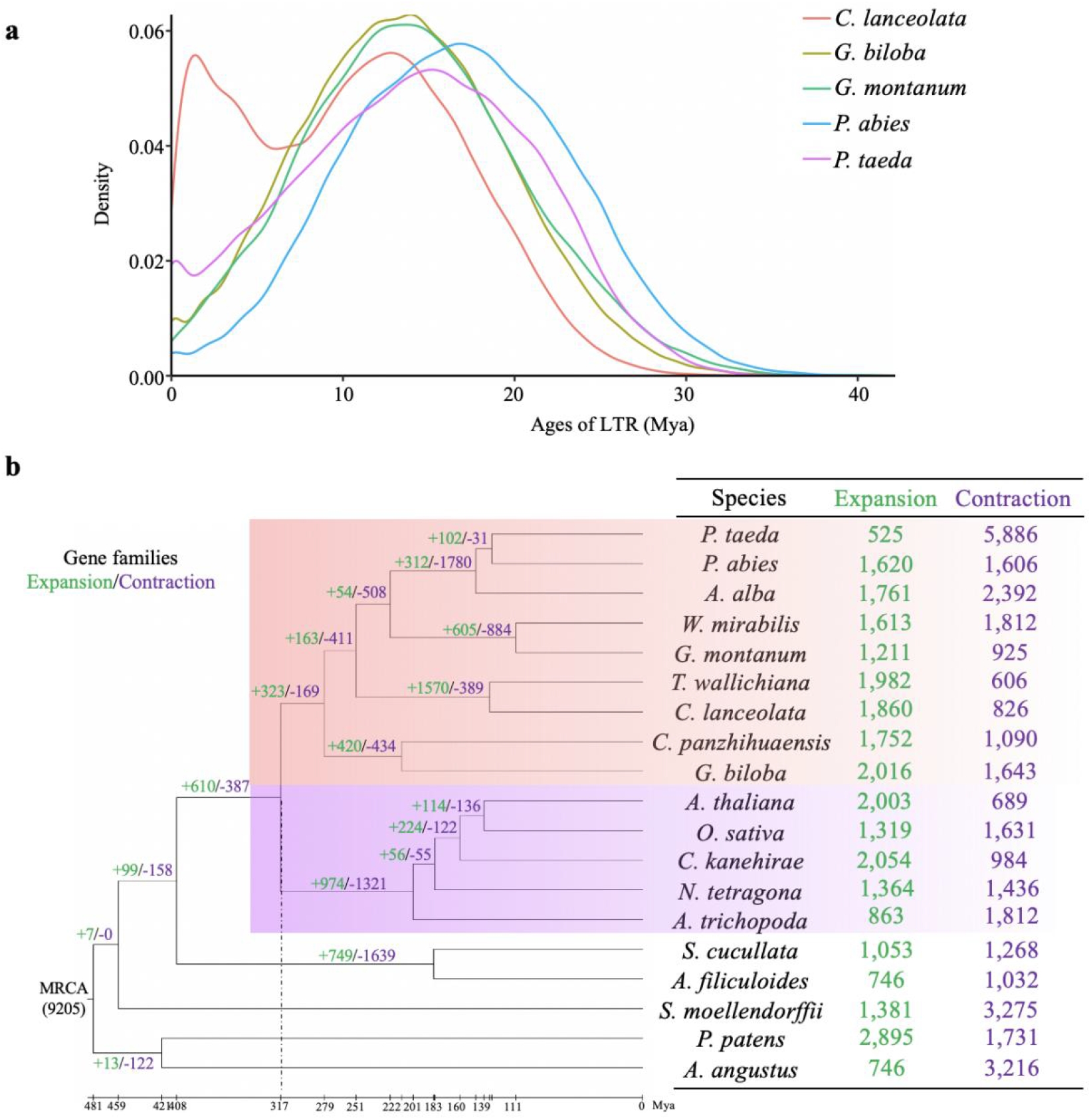
Genome evolutionary history of *C. lanceolata*. **a.** LTR insertion time and frequency of *C. lanceolata*, *G. biloba*, *G. montanum*, *P. abies*, and *P. taeda*. LTR insertion frequency of *C. lanceolata* displayed two obvious peaks. One occurred at 3–7 Mya and the other at 14–18 Mya. Only one peak was present in the other gymnosperms. The results indicate the stable accumulation of LTR in the genome of *C. lanceolata*. **b**. Phylogenetic tree, gene family expansion and contraction, and divergence time in genomes of 19 species. Phylogenetic tree was constructed based on single-copy gene from the 19 species (see **Methods**). The most recent common ancestor (MRCA) that have expanded or contracted during recent differentiation have 9,205 gene families. The green numbers are the numbers of expanded gene families, and the purple numbers are the numbers of contracted gene families. The area covered by rose red represents gymnosperms, and the area covered by purple represents angiosperms. The dotted line marks the differentiation time of angiosperms and gymnosperms.

### Evolution of gene families

We found that 797 gene families are unique to the *C. lanceolata* genome by comparing the orthologous genes of different plants with 19 known plant genomes (See **Methods**, **Supplementary Table 14,** and **Supplementary Fig. 3**). Expansion and contraction analysis of orthologous gene families revealed that 1,860 gene families in *C. lanceolata* expanded. Of these, 120 gene families significantly expanded (*P* < 0.01) and 826 gene families contracted, of which four gene families were significant (**Figs. 1b, 1c**). Enrichment analyses revealed that the significantly expanded gene families were enriched in the Kyoto Encyclopedia of Genes and Genomes (KEGG) pathways of ‘monoterpenoid biosynthesis (map00902)’ ‘plant hormone signal transduction (map04075)’ ‘steroid hormone biosynthesis (map00140)’ and ‘betalain biosynthesis (map00965)’ (**Supplementary Table 15**), and were enriched in the gene ontology (GO) terms of ‘lignin biosynthetic/metabolic process (GO:0009809/GO:0009808)’ and ‘terpene synthase activity (GO:0010333)’ (**Supplementary Table 16**). Furthermore, we found two significantly expanded gene families: Beta-glucosidase (OG0000016) and Laccase (OG0000039), which lead the GO enrichment to lignin metabolic process (GO:0009808) (**Supplementary Table 17**). Lignin biosynthesis is formed by hydrolysis of its precursor glucoside by β-glucosidase and then catalyzed by Laccase and peroxidase systems. Interestingly, we found that 58% (28/48) and 51% (20/39) of genes significantly expanded in Beta-glucosidase and Laccase were tandem repeat genes, respectively (**Supplementary Tables 18 and 19**). Thus, such terms may be important for secondary growth in gymnosperms, and most of them undergo tandem replication.

### Phylogeny of gymnosperms

The branching order among extant gymnosperms major lineages has remained controversial, especially for Ginkgo and Gnetales^39^. On the basis of the single-copy families derived from 19 gymnosperms species with known genomes (see **Methods**), we constructed phylogenetic trees from the concatenated sequence alignments and astral sequence alignments of both nucleotide and amino acid sequences. The concatenated tree based on amino acid strongly support that extant gymnosperms and angiosperms are reciprocally monophyletic groups, *G. biloba* plus *Cycas panzhihuaensis* as a sister group of the remaining extant gymnosperms, Gnetales (*W. mirabilis* and *G. montanum*) within conifers sister to Pinaceae (*A. alba*, *P. abies*, and *P. taeda*), and *T. wallichiana* – *C. lanceolata* clade as a sister group of Gnetales – Pinaceae (**Supplementary Fig. 4a**). However, other gene trees support Gnetales as a sister group of the remaining extant gymnosperms, and *G. biloba* – *C. panzhihuaensis* clade is sister to conifers (*C. lanceolata*, *T. wallichiana*, and Pinaceae) (**Supplementary Fig. 4b**). Trees based on amino acids and nucleotides are different, a phenomenon associated with character state information when using concatenated data, i.e. model specification and parallelism/inversion in protein sequences being different from coding sequences which may be affected by substitution saturation.

Furthermore, similar to angiosperms, the branch lengths of *W. mirabilis* and *G. montanum* in each gene tree are significantly longer than that of other gymnosperms. (**Supplementary Fig. 4**). We speculate that *W. mirabilis* and *G. montanum* clade are located in base of gymnosperms because of the long-branch attraction (LBA) effect. Strategies to avoid LBA artifacts include excluding long-branch taxa and fast-evolving third codon positions, as well as using inference methods less sensitive to LBA^40^. We further selected the first two bases of the codon for nucleotides to construct concatenated and astral trees, and used PhyloBayes with the ‘cat model’ (which can effectively avoid LBA artifacts) to construct the Bayesian phylogenetic tree. The results of the Bayesian and concatenated trees support that *G. biloba* – *C. panzhihuaensis* is sister to the remainder of the extant gymnosperms, whereas only the astral tree resolved Gnetales in that position (**Supplementary Fig. 4**).

Our results tend to place Ginkgo sister to cycads, Gnetales within conifers sister to Pinaceae, and Cupressaceae in different clades of Pinaceae. Recent phylogenomic studies were based on nuclear genomes obtained from the expanded sampling of gymnosperm taxa or various phylogenetic approaches. The results provide strong support for the main topology inferred here^18,39,41^. In addition, the main conflict in the constructed gene trees is whether the sister group of the extant gymnosperms is *G. biloba* – *C. panzhihuaensis* or Gnetales. This conflict suggests the possibility of reticulation in the early evolution of gymnosperms, which needs further research.

### WGDs

WGD is a major driving force in plant evolution^42,43^. We conducted a collinearity analysis on chromosome-level genomes of six gymnosperms species, including *G. biloba*, *C. panzhihuaensis*, *C. lanceolata*, *T. wallichiana*, *W. mirabilis*, and *G. montanum*. Based on the collinearity result, orthologous gene pairs in the collinear block were extracted to calculate *K*s to infer WGDs. The clear *K*s peaks of *G. biloba* and *C. panzhihuaensis* of approximately 0.89, and approximately 1.47 and 1.41 for *C. lanceolata* and *T. wallichiana*, respectively, indicate that they experienced a WGD event (**Fig. 2a**). The collinearity relationship of *G. biloba*, *C. panzhihuaensis*, *C. lanceolata*, and *T. wallichiana* was almost 1:1. This implies that these species have the same WGD history, with no branch-specific WGD event occurring after differentiation (**Fig. 2a**). Thus, the common ancestor of these four species experienced an ancient WGD event. Combined with *K*s tree analysis (see **Methods**), we verified that this ancient WGD event was shared by all gymnosperms, and occurred approximately 295 Mya after the differentiation of angiosperms and gymnosperms (**Figs. 2b, 2c**). Due to the gymnosperm-wide ancient WGD event occurred too long ago, and the gene evolutionary rates of *W. mirabilis* and *G. montanum* genomes were faster than those of other gymnosperms. Thus, the signal of paralog *K*s peak representing this ancient polyploidy event is too weak to be detected. Furthermore, a signature *K*s peak for paralogous genes in the genome of *W. mirabilis* was observed, but was absent in the *G. montanum* genome (**Fig. 2a**). This peak indicated that the WGD event specific to *W. mirabilis* occurred at approximately 101 Mya (**Figs. 2b, 2c**). A Pinaceae-specific WGD event has been disputed in several studies^19,44^. We performed collinearity and *K*s analyses on the three Pinaceae species (*P. taeda*, *P. abies*, and *Abies alba*) that were not assembled to the chromosome-level (see **Methods**). The results showed that Pinaceae experienced an ancient WGD event shared by all gymnosperms, but no unique WGD event has occurred in Pinaceae (**Supplementary Fig. 5**). The currently available low-quality genome of Pinaceae species possibly prevented the detection of a Pinaceae-specific WGD event.

**Figure 2.**
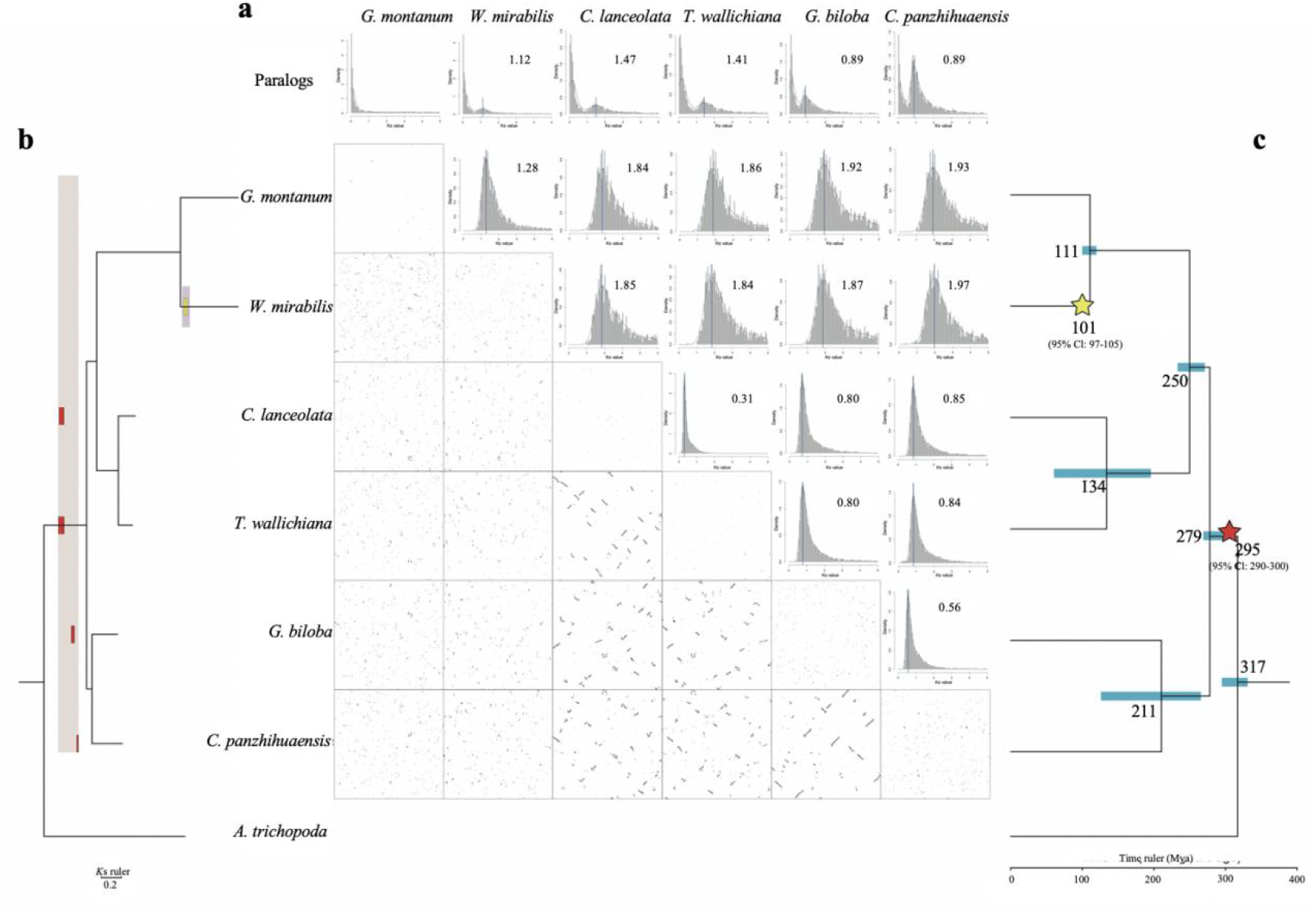
WGD analyses of gymnosperms. **a.***K*s distribution of *G. montanum*, *W. mirabilis*, *C. lanceolata*, *T. wallichiana*, *G. biloba*, and *C. panzhihuaensis* (upper right) and the collinear analysis between them (lower left). **b**. *K*s tree inferred based on the *K*s rate. The evolution rate of *G. montanum* and *W. mirabilis* is faster than other gymnosperms and the *K*s tree can more accurately determine the location of the gymnosperm species where WGD event occurs. The peak of *K*s differentiation between two species in the *K*s tree can be obtained by adding up the branch length between the two species. Based on the peak in the *K*s distribution map of each species (i.e., the corresponding WGD), starting from the end of the branch (right) and reversing the direction enables location of the WGD in the *K*s tree. WGD events of *C. lanceolata*, *T. wallichiana*, *G. biloba* and *C. panzhihuaensis* are obviously located in the ancestral branch of gymnosperms (red rectangle) and the WGD event of *W. mirabilis* is located in the branch of *W. mirabilis* (yellow rectangle). **c.**The occurrence time and locations of gymnosperm WGD events. The red five-pointed star represents the unique WGD of gymnosperms, and the yellow five-pointed star represents the unique WGD of *W. mirabilis*.

The phylogenetic placement of ancient polyploidization events could be achieved by both age-distribution-based methods with proper correction of substitution rate^45^ and reconciliation-based phylogenomic method with proper model of gene family evolution^46^. The aforementioned result of age-distribution analysis shows that there is a shared ancient polyploidization event at the base of gymnosperms. Here we use the model-based phylogenomic software WHALE to test the validity of this gymnosperm-shared WGD event and explore other potential WGD events. The species tree adopted consists of gymnosperm clade (six species), angiosperm clade (six species), fern clade (two species) and lycophyte clade (one species) as outgroup. The preset WGD models were put on the branch leading to seed plant clade (seed plants WGD, labelled as WGD1), gymnosperm clade (gymnosperms WGD, labelled as WGD2), the *C. lanceolata* and *Sequoidendron giganteum* (assumed to be Cupressaceae WGD, labelled as WGD3), the *C. lanceolata* (assumed to be *Cunninghamia* WGD, labelled as WGD4), and angiosperm clade (angiosperms WGD, labelled as WGD5), respectively. The relaxed branch-specific DL+WGD and critical branch-specific DL+WGD model were implemented for the inference of the significance and retention rates (*q*) of the assumed WGD events in the manner of Bayesian inference (BI). The significance of assumed WGD events was inferred by means of calculating the Bayes Factor (*K*) of the posterior distributions of retention rates using the Savage-Dickey density ratio^46^.

The results (**Supplementary Table 20**) show that the estimated retention rates of both Cupressaceae WGD (WGD3) and *Cunninghamia* WGD (WGD4) are not significantly larger than zero, indicating the absence of WGD events on the corresponding branches, which is consistent with the aforementioned results that *C. lanceolata* is lack of recent WGD events. With the DL+WGD model under relaxed branch-specific rates, the results show that the gymnosperms WGD, angiosperms WGD and seed plants WGD all have retention rates over 0.1. With the DL+WGD model under critical branch-specific rates, the results show that only the gymnosperms WGD and seed plants WGD have retention rates over 0.1 while the angiosperms WGD has a small retention rate of 0.00332 which is not significantly larger than zero, probably due to the model violation of the critical branch-specific rates DL+WGD modelling for distinct clades. The Bayesian inference of duplication and loss rates for each branch (**Fig. 3**) also indicate that the branch leading to seed plants and gymnosperms clade has highest duplication rate and lowest loss rate. Integrally, the results from the phylogenomic analysis confer significant and solid support for the validity of seed plants-specific WGD and gymnosperms-specific WGD.

**Figure 3.**
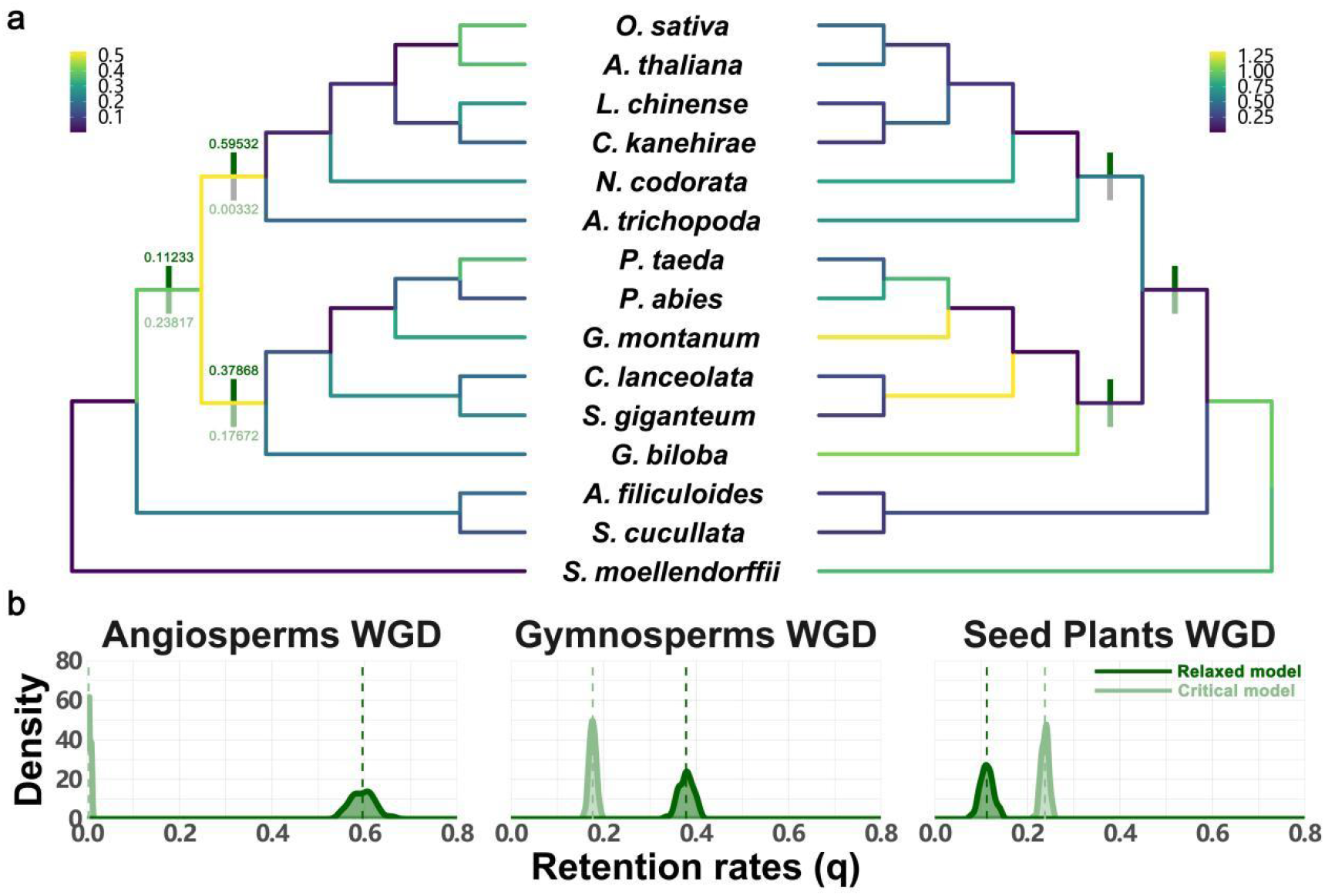
Branch-specific rates DL+WGD modeling analysis in Whale. **a.** Posterior mean of duplication (left) and loss (right) rates estimated under DL+WGD modelling colored on the clade tree. Dark green bars indicate the supported WGDs under relaxed branch-specific model with the posterior mean annotated. Light green bars indicate the supported WGDs under critical branch-specific model with the posterior mean annotated. Grey bar indicates WGD with a retention rate estimate not significantly different from zero. **b.** Kernel density estimates of the marginal posterior distributions for the retention rates of the angiosperms WGD, the gymnosperms WGD and the seed plants WGD under relaxed and critical branch-specific model.

In conclusion, the above results indicate that the ancestor of seed plants has differentiated into angiosperms and gymnosperms after having experienced a WGD event. The ancestor of extant gymnosperm, then, has experienced a gymnosperm-specific WGD event and diverged into the five extant gymnosperm lineages, which have not experienced further paleo-polyploidization events themselves, except for *W. mirabilis*. The extant angiosperms do not share a common WGD before their most recent common ancestor diverged into existing angiosperms lineages. However, almost every major lineage of angiosperms has experienced polyploid events themselves^47,48^, except for a few species, such as *Amborella trichopoda*^49^ and *Aristolochia fimbriata*^50^. Overall, these ancient polyploidy events provide an evolutionary basis for the formation of species diversity patterns of seed plants.

### Evolution of reproductive organs

Seed plants diverged into angiosperms and gymnosperms approximately 317 Mya (**Fig. 1**). The main difference between angiosperms and gymnosperms is the extreme morphological difference in their reproductive structures^51,52^. MIKCc-type MADS-box genes of angiosperms encode transcription factors that control floral organ morphogenesis and flowering time in flowering plants. The *A*(*AP1*/*FUL*), *B*, *C*/*D*(*AG*/*STK*), and *SEP* subfamilies are the major components in the well-known ‘ABCDE’ model in flowering plants that describes the roles of the four subfamilies in the development of petals, sepals, stamens, and ovaries, respectively^53,54^. This model is not suitable for gymnosperms because their reproductive organs do not have standard floral structures, such as the male and female cones of *C. lanceolata*. However, analyzing the MADS-box gene family of *C. lanceolata* can reveal the development of reproductive structures in gymnosperms, and clarify the origin of flowers in angiosperms. We identified 50 MADS-box genes in the *C. lanceolata* genome, including 32 type II genes (27 MIKCc-type genes and five MIKC*-type genes) and 18 type I genes (11 *Mα* genes and seven *Mγ* genes) (**Supplementary Table 21** and **Supplementary Fig. 6**). All the *Mβ* genes in *C. lanceolata* were lost (**Supplementary Fig. 6**). The loss of type I genes is less harmful than loss of type II genes^55^. Expression analysis showed that all type I genes were not or negligibly expressed in any organs (**Supplementary Fig. 7**).

The MIKCc-type genes of *C. lanceolata* were classified as *AGL6* (two members), *AGL32* (seven members), *AG* (six members), *GMADS* (two members), *TM8* (six members), *SOC1* (one member), and *SVP* (three members) (**Supplementary Table 21**). In the BC model of gymnosperm reproductive organ development, the B+C classes specify male organs, and the C class specify female organs^52^. Interestingly, two different expression patterns of *C. lanceolata AG*-like (C class) and *AGL32*-like (B class) genes have been revealed. The *AG*-like genes *Cl10262*, *Cl36126*, and *Cl26543* were all expressed in the reproductive organs, while *Cl13850*, *Cl35439*, and *Cl26063* were only expressed in the roots (**Fig. 4a**). The *AGL32*-like gene *Cl14691* is mainly expressed in male cones, female cones, and bract scales, while *Cl08169* is mainly expressed in female cones, male cones, seeds, homozygous female gametophytes, and seed scales (**Fig. 4a**). In addition, gymnosperms have two diverged *AGL6* genes with different functions, with one specific to gymnosperms and the other shared by both gymnosperms and angiosperms (**Supplementary Fig. 6**). Two *C. lanceolata AGL6*-like genes, *Cl29520* and *Cl35065*, exhibit two different expression patterns. *Cl29520* is only expressed in reproductive organs (male cones, female cones, seed scales, and bract scales), while *Cl35065* is expressed in reproductive and vegetative organs (**Fig. 4a**). The expression analysis showed that *AGL6*, *B,* and *C* genes may together determine the development of gymnosperm reproductive organs (**Fig. 4**).

**Figure 4.**
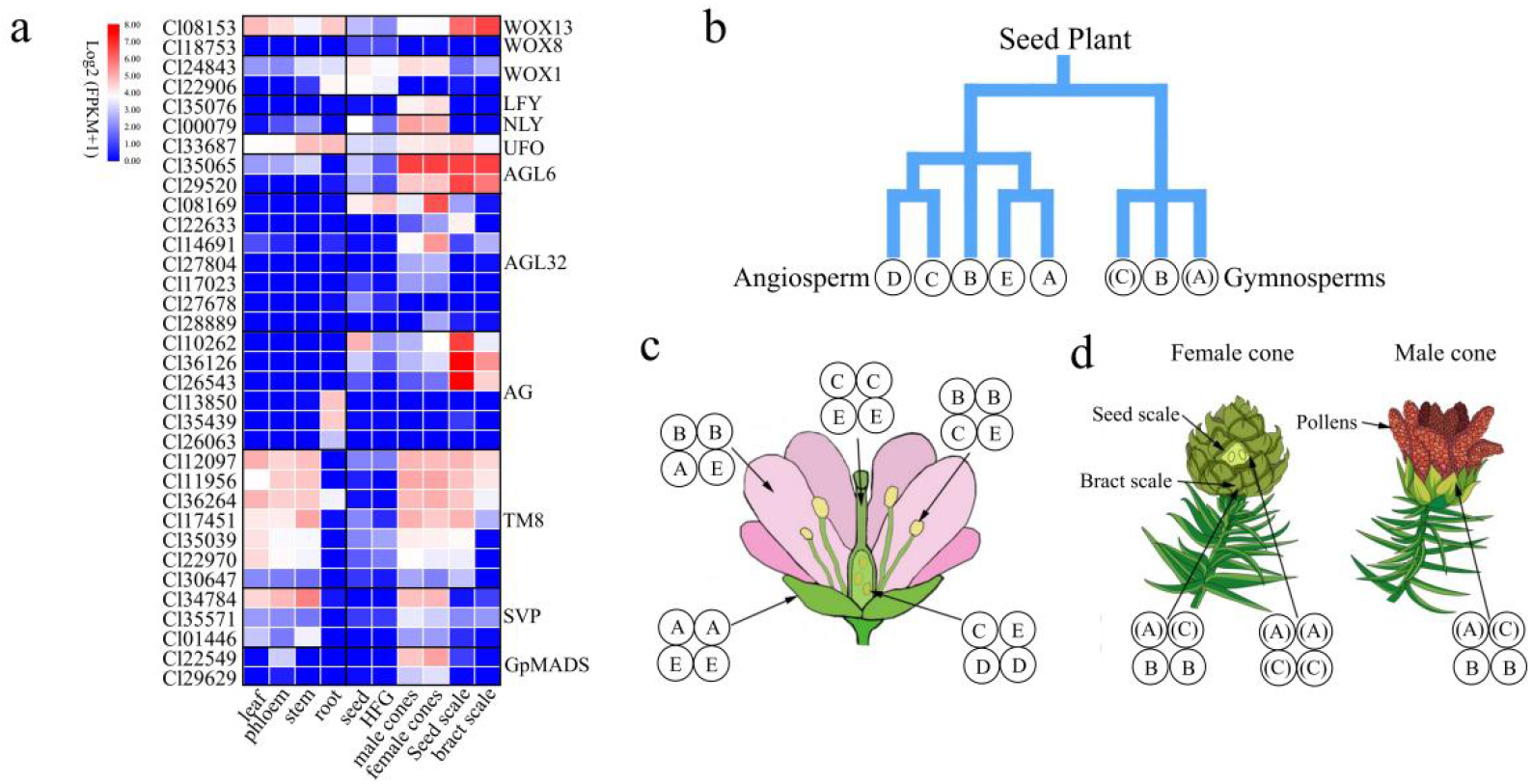
Analysis of reproductive organ genes involved in regulation in *C. lanceolata*. **a**. Expression profile of MIKCc-type of *C. lanceolata.* HFG, homozygous female gametophyte. **b**. Homologous genes regulating the development of reproductive organs in angiosperms and gymnosperms. The (A) from the gymnosperm reproductive organ development model (A)B(C) is homologous to the A and E classes gene in the angiosperm flower development ABCE model. (C) is homologous to the C and D classes gene (**Supplementary Fig. 6**). **c**. Angiosperm flower model. **d**. *C. lanceolata* female (left) and male (right) regulation model.

The *AGL6* subfamily in gymnosperms is divided into two clades. One is widely expressed in reproductive and vegetative organs. The other is only expressed in male and female reproductive tissues^56,57^. Phylogenetic tree analysis revealed that *A*, *SEP*, and *AGL6* genes from gymnosperms and angiosperms have a common ancestor, and that the *A* and *SEP* genes of gymnosperms have been lost (**Supplementary Fig. 6**). Thus, the *AGL6* subfamily of gymnosperms may be the only *E* function gene similar to the ABCDE model of angiosperm flower development^57^. The proposed gymnosperm (A)B(C) model suggests that the B-(A)-(C) genes are designated the male cone, while the (A)-(C) genes specify the female cone organ^58^ (**Fig. 4b**). The seed cones of *C. lanceolata* are usually indicative of the typical female reproductive unit. The seed cones are a determinate axis bearing multiple, spirally arranged seed scales that are axillary associated with bract scales (**Fig. 4c**). The bract scales are considered abnormal leaves without fertility, while seed scales are considered to be abnormal leaves with fertility (**Fig. 4c**). Based on expression analysis, the AGL6-like (*Cl29520*), AG-like (*Cl10262*, *Cl36126*, and *Cl26543*), and AGL32-like (*Cl14691*) genes of *C. lanceolata* collectively regulate male cones and bract scales, and the AGL6-like (*Cl29520*) and AG-like (*Cl10262*, *Cl36126*, and *Cl26543*) genes collectively regulate seed scales. We continue to perfect the gymnosperm (A)B(C) model: B-(A)-(C) specifies the bract scales and male cone, and (A)-(C) specifies the seed scales (**Fig. 4c**).

*LEAFY*/*FLORICAULA* (*LFY*), *NEEDLY* (*NLY*), *UNUSUAL FLORAL ORGANS* (*UFO*), and *WUSCHEL*-related homeobox (*WOX*) genes regulate the MADS-box genes determining floral meristem^59,60^. *LFY* and *NLY* of gymnosperms are important for the switch between the vegetative and reproductive phase. *LFY*-dependent activation of the homeotic B-like gene requires *UFO* activity^61^. The repression of *WOX* by *AG* is essential for terminating the floral meristem and *WOX* can induce *AG* expression in developing flowers^62^. The *LFY*-like gene *Cl35076* and *NLY*-like gene *Cl00079* are highly expressed in male and female cones (**Fig. 4a**). The *UFO*-like gene *Cl33687* is expressed in the reproductive and vegetative organs (**Fig. 4a, Supplementary Fig. 8**). A total of 36 *WOX* genes were identified in *C. lanceolata,* much higher than those found in Ginkgo and *P. abies* (**Supplementary Fig. 9**). The ancient clade *WOX13*-like gene *Cl08153* is expressed in all tissues (**Fig. 4a**), similar to the *P. abies PaWOX13* gene^63^, suggesting that the expression pattern of the ancient clade genes is similar among gymnosperms. The modern clade gene *ATWOX1* plays an important role in the patterning and morphogenesis of the early embryo^64,65^. The expression of the *WOX1*-like gene *Cl24843* was highest in male and female cones compared to that in other tissues (**Fig. 4a**). *AtWOX8/9* has been implicated in the patterning and morphogenesis of the early embryo^66–68^ and *PaWOX8/9* is preferentially expressed during embryo development^69^. The *WOX8*-like gene *Cl18753* was expressed in the seed and homozygous female gametophyte, but not in other tissues (**Fig. 4a**). In summary, the *LFY*, *NLY*, *UFO*, *WOX1*, and *WOX8* genes in *C. lanceolata* may regulate the MADS-box genes related to the development of *C. lanceolata* reproductive organs. The regulatory network warrants further investigation.

### Formation of astringent seeds

Nonflowering seed plants (conifers, cycads, Ginkgo, Ephedra) form a large homozygous female gametophytes to nourish the embryo within a seed. But some homozygous female gametes will miscarry and form astringent seed. An astringent seed is a type of abortive phenomenon paralleled in many lineages of seed plants. The proportion of astringent seeds in *C. lanceolata* seed orchards is as high as 30%^70^, which far exceeds other species and which has a substantial negative influence on the production of excellent seeds. Why *C. lanceolata* has retained the energy consuming trait of astringent seeds for so long in its evolutionary history is puzzling, since there seems to be no reward. Unlike other defective seeds, astringent seeds normally develop into a shape similar to that of normal seeds after abortion, including size, weight, and seed coat color (**Fig. 5a**). We examined semi-thin sections using light microscopy to observe the structure of astringent and germinating seeds at different developmental stages (**Supplementary Note 1**). As shown in **Fig. 5a** and **Fig. 5b**, in the early stage of the formation of the astringent seed (95–105 d), the embryo has been aborted and the homozygous female gametophyte has partially disintegrated, causing the entire astringent seed to shrink. Subsequently, a cavity is formed between the embryo and homozygous female gametophyte tissue. The cavity is gradually filled with secondary metabolites, causing the seed to re-expand, and form a plump state that looks like a germinated seed (105–185 d).

**Figure 5.**
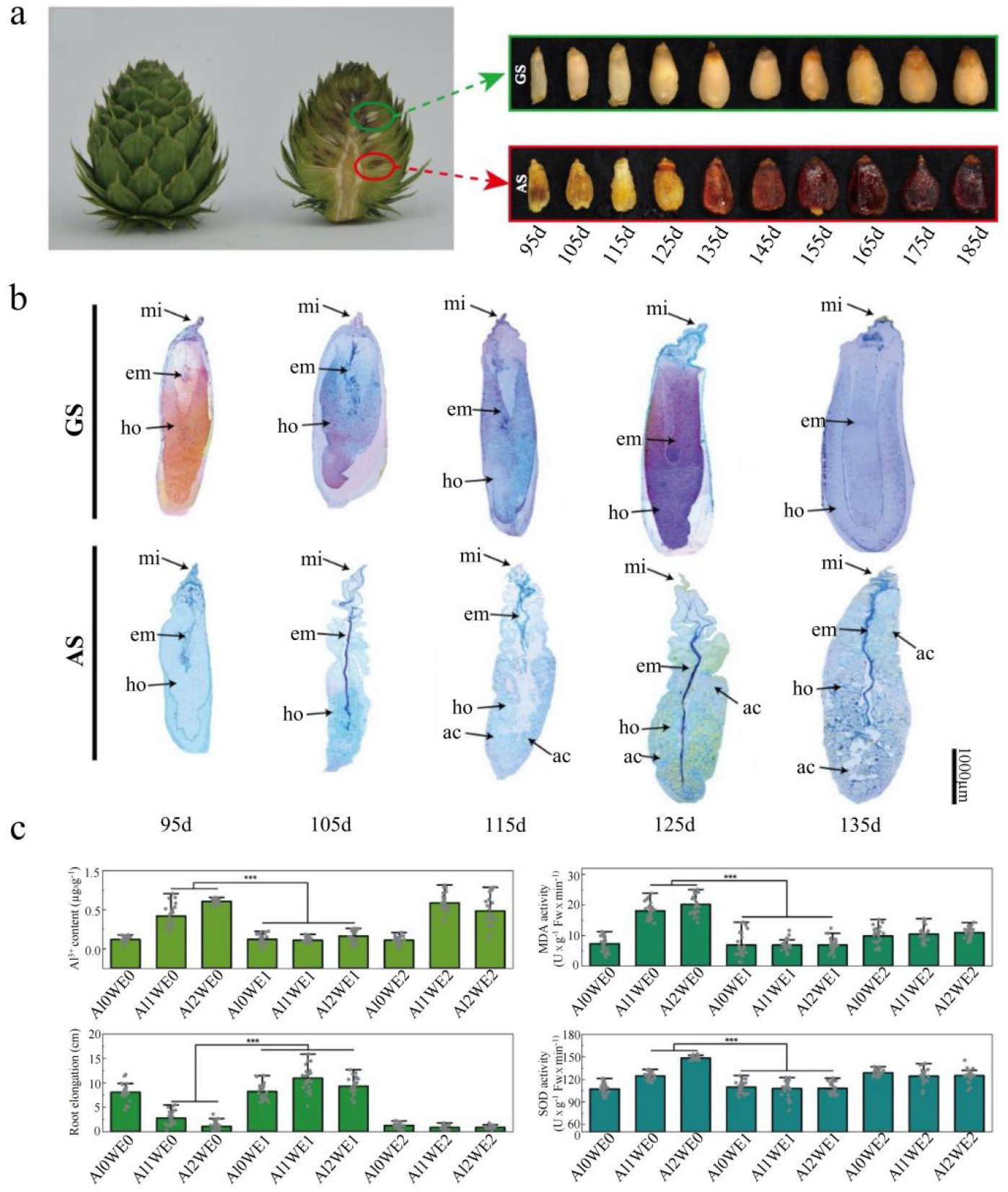
Astringent seeds (AS formation and mechanism. **a**. Phenotypic characteristics of the development of germinating seeds (GS) and AS. **b**. Microscopic observation of AS and GS in different development stages. mi: micropyle; em: embryo; ho: homozygous female gametophyte; ac: astringent compound. Microscopic scanning showed that GS has a complete embryo structure during development, while AS has aborted embryos, broken homozygous female gametophytes, and accumulated secondary metabolites. **c**. Statistics of root growth of *C. lanceolata* seedlings under different aluminum and astringent seed water extract concentration combinations after culturing for 30 days. Al0, Al1, and Al2 represent aluminum concentrations of 0.00, 0.50, and 1.00 mmol/L, respectively. WE0, WE1, and WE2 represent AS water concentrations of 0.00, 1.00, and 10.00 g/L, respectively. The histogram demonstrating that with the aggravation of Al stress, the Al3+ ions in root tip tissue of *C. lanceolata* seedlings increased significantly, and the relative root elongation was strongly inhibited. Moreover, the activities of malondialdehyde (MDA) and superoxide dismutase (SOD) in the root tip tissues significantly increased. This indicated a negative effect of Al stress and pronounced damage to root tissues. However, if a low concentration of water extract of AS was added under the Al stress, the performance of these indexes was equivalent to that without stress, indicating that the low concentration of AS water extract can effectively alleviate the adverse effects of Al stress on root growth. Further, high concentration of this extract was also detrimental to the root growth, regardless of the presence of Al stress.

The distribution of *C. lanceolata* coincides with the distribution of acid red soil (**Supplementary Fig. 10**). The acidification of the soil causes a large amount of active aluminum to be released, significantly increasing the aluminum content in the soil. This increases the toxicity of aluminum to *C. lanceolata*^71,72^. We observed that the content of aluminum ions in the root tip tissue of *C. lanceolata* seedlings increased significantly under aluminum stress, leading to damage to root tip tissue, and inhibiting root elongation (see **Methods** and **Fig. 5c**). These inhibitory effects are similar to those of other plants growing in acidic red soil^73–75^. Interestingly, low concentrations of astringent seed extraction can alleviate these inhibitory effects (**Supplementary Note 2** and **Fig. 5c**). We suggest that after the seed cones fall off, the astringent seeds dissolve the secondary metabolites under natural conditions, inhibiting the toxicity of aluminum and providing an environment for the surrounding seeds to germinate and grow. This may be a long-term protective response strategy for the astringent species of *C. lanceolata* to deal with environments high in aluminum.

Flavonoid compounds are the main component in astringent seeds, accounting for more than 30% of the content^76^. Metabolome sequencing of different stages of astringent seeds revealed the significantly increased content of flavonoid metabolites with the development of astringent seeds from 105 d after pollination (**Supplementary Note 3** and **Supplementary Table 22**). A total of 142 genes related to flavonoids and flavonoid biosynthesis have been identified in the *C. lanceolata* genome (**Supplementary Table 23**). Of these, the flavonoid synthesis-related genes flavanone-3-hydroxylase (*F3H*), flavonoid 3’-hydroxylase (*F3*’*H*), flavonoid 3’,5’-hydroxylase (*F3*’*5*’*H*), and flavonol synthase (*FLS*) have been duplicated (**Supplementary Figs. 11, 12**). A total of 29 genes related to flavonoid biosynthesis were highly expressed in astringent seeds after embryo abortion, but not in germinating seeds (**Supplementary Fig. 13**). Embryo-defective genes (EMBs) are a class of genes related to plant seed development and defects^77^. We identified 771 EMBs in the *C. lanceolata* genome (**Supplementary Table 24**). Of these, 109 EMBs displayed different expression levels between astringent and germinated seeds (**Supplementary Fig. 14**). The expression of the *PPT1*-like gene (*Cl01236*) had the highest level of differential expression between astringent and germinating seeds in the early stages (**Supplementary Fig. 14**). *PPT1* regulates the accumulation of reactive oxygen species (ROS)^78^. Excessive accumulation of ROS is often accompanied by a strong redox response in tissues, leading to cell death^79^. These findings suggest that the high expression of *PPT1* gene in astringent seeds may be involved in the seed damage and abortion.

Some plants can store activated aluminum ions in the vacuoles of mesophyll cells in the form of aluminum ion-organic acid complexes. This storage reduces the toxicity of aluminum to roots^80^. In contrast to most tree species, *C. lanceolata* leaves can survive on the branches for at least three years after withering, and do not fall off with the branches until self-pruning occurs. We observed that with prolonged survival of leaves on withered branches, the aluminum content in the leaves increased significantly, while the aluminum content in leaves that fell off of withered branches was significantly reduced (**Supplementary Note 4** and **Supplementary Fig. 15**). The collective findings support the view that *C. lanceolata* may prevent the rapid return of aluminum ions to the environmental soil through prolonged leaf persistence.

### Shade tolerance of seedlings

Plants depend on light for photosynthesis, with different needs for light at different growth stages. *C. lanceolata* seedlings do not grow well under full shade and require proper shade for healthy growth^81^. The dense canopy under natural conditions also provides an environment for the growth of *C. lanceolata* seedlings. Shade indicates a limited low red (660 nm)/far-red (730 nm) photon flux ratio (R:FR) and photosynthetically active radiation (PAR) for plants^82^. R:FR provides a highly reliable indication of shade because it is relatively unaffected by other environmental conditions^83^. Plants sense the environment R:FR signal via the phytochrome family of photoreceptors^84^. The phytochrome apoproteins are encoded by a small family of genes involving three main clades: *PhyA*, *PhyB*, and *PhyC*^84^. Low R:FR of shade reduces the levels of active *PhyB*, increasing the activity of phytochrome-interacting factors (PIFs), which promote auxin-synthesis genes, in turn promoting stem growth^85^. Full shade avoidance responses require PIF3 and PIF7 under low R:FR or natural shade light^86–89^. We identified four PhyB, three PIFs (two PIF3 and one PIF7) (**Supplementary Table 25**), and 20 *AUX*/*IAA* genes in the *C. lanceolata* genome (**Supplementary Fig. 16**). Expression analysis showed that the expression of the *PhyB*-like gene *Cl22557* in the leaves of seedlings and adult trees (lower, middle, and upper layers) was extremely low (**Supplementary Fig. 17**). Two *PIF3*-like genes (*Cl23109* and *Cl02626*) and one *PIF7*-like gene (*Cl30879*) were expressed in seedling leaves, but their expression was extremely low in adult tree leaves (**Supplementary Fig. 17**). The expression of 11 *AUX*/*IAA* genes in seedling leaves was higher than that in adult tree leaves (**Supplementary Fig. 17**).

Sensory photoreceptors involved in shade light include cryptochromes (*CRY*), phototropins (*Phot*), and *UV RESISTANCE LOCUS 8* (*UVR8*)^85^. All can perceive changes in irradiance. *Arabidopsis CRY1* activity can enhance plant defenses and *CRY2* contributes to some shade responses^90^. Both *Arabidopsis Phot1* and *Phot2* cause chloroplasts to accumulate at the periclinal wall of palisade cells under low irradiance^91,92^. This increases the efficiency of light capture^93^, which helps plants under shade make full use of limited PAR. We observed the low expression of two *CRY2*-like genes (*Cl24639* and *Cl22470*) in the leaves of seedlings, but negligible expression in the leaves of adult trees. One *CRY1*-like gene (*Cl17630*) and one *Phot2*-like gene (*Cl14893*) were expressed in all tissues. Their expression in seedlings was higher than that in the leaves of adult trees. Two *Phot1*-like genes (*Cl25517* and *Cl04142*) were negligibly expressed in all tissues (**Supplementary Fig. 17**). In shade, ultraviolet (UV) B levels are significantly reduced, stimulating *UVR8* to activate transcriptional genes involved in the UV-protective response to induce flavonoid synthesis^94–96^. We observed duplication of eight *UVR8* genes in *C. lanceolata* (**Supplementary Table 25**). Of these, three were highly expressed in seedling leaves, expression of four was low in all tissues, and one was not expressed in any tissue (**Supplementary Fig. 17**). We also identified 38 genes related to flavonoid synthesis that were highly expressed in seedlings (**Supplementary Fig. 18**). The findings indicate that the *UVP8* gene can induce the synthesis of flavonoids in the leaves of *C. lanceolata* seedlings, and flavonoids can improve plant resistance. Shade avoidance also requires constitutive photomorphogenesis 1 (*COP1*). *CRY1* and *PhyB* will reduce the activity of *COP1*, while *UBR8* will increase the activity of *COP1*^97,98^. *COP1*, *CRY1*, and *Phot* regulate light-controlled stomatal development and stem elongation in *Arabidopsis*^99^. We identified the expression of one *COP1*-like gene (*Cl21889*) in the leaves of seedlings, with negligible expression of another gene (*Cl14590*) in all tissues (**Supplementary Fig. 17**). Based on the aforementioned shade avoidance-related gene expression profiles, we constructed a molecular network regulation diagram of the shade tolerance of *C. lanceolata* seedlings (**Supplementary Fig. 17**). The diagram reveals the molecular mechanism of the shade tolerance of *C. lanceolata* seedlings.

## Conclusion

This study describes the assembly, annotation, and comparative analysis of the genome of the first Cupressaceae species, *C. lanceolata*. To avoid LBA artifacts, we used different phylogenetic methods. The findings support that Ginkgo and *C. panzhihuaensis* are sister groups, *G. biloba* plus *C. panzhihuaensis* is a sister group of the remaining extant gymnosperms, Gnetales (*W. mirabilis* and *G. montanum*) within conifers is sister to Pinaceae (*A. alba*, *P. abies*, and *P. taeda*), and *T. wallichiana* – *C. lanceolata* clade is a sister group of Gnetales –Pinaceae. WGD analysis showed the ancestor of seed plants has differentiated into angiosperms and gymnosperms after having experienced a WGD event. The ancestor of extant gymnosperm has experienced a gymnosperm-specific WGD event and the extant angiosperms do not share a common WGD before their most recent common ancestor diverged into existing angiosperms lineages. A total of 50 MADS-box genes were identified in *C. lanceolata* genome. Their expression patterns in vegetative and reproductive organs confirmed that the ‘(A)B(C)’ model of gymnosperm reproductive organs and the ‘ABCDE’ flowering model of angiosperms are derived from the ‘(A)B(C)’ model of gymnosperm reproductive organs. Formation of astringent seeds and the shedding of the whole branches (with withered leaves) of *C. lanceolata* have occurred during adaptation to acid soil. The main secondary metabolites of astringent seeds are flavonoids, and genes related to flavonoid synthesis were amplified. Finally, a molecular regulatory mechanism for seedlings to avoid shade is proposed. Our results provide new insights into the ancestral genome characteristics in seed plants and the genome evolution and diversification of gymnosperms.

## Methods

### DNA preparation and sequencing

All the genome sequencing materials used in this study were collected from an adult *C. lanceolata* plant growing in the *C. lanceolata* third-generation seed orchard of Youxi National Forest Farm in Fujian Province, China. Sodium dodecyl sulfate-based lysis was used to extract total genomic DNA from young leaves for Illumina and PacBio sequencing. For Illumina sequencing, DNA was ultrasonicated to a fragment size of 270 bp and the library was prepared using an Ultra DNA Library Prep Kit (NEB) according to the manufacturer’s instructions. Sequencing of the library was performed using Hiseq4000. Sequencing reads were generated from the paired-end. The library was trimmed using fastq_quality_trimmer in the FASTX-Toolkit (ver. 0.0.11) with default parameters. For PacBio sequencing, DNA was interrupted using g-TUBE (Covaris), and SMRTbell template preparation involved DNA concentration, damage repair, end repair, ligation of hairpin adapters, and template purification, performed using AMPure PB Magnetic Beads (Pacific Biosciences). The PacBio Sequel platform was used to perform 20 kb single-molecule real-time DNA sequencing.

### Genome size assembly

Eight 270 bp pair-end libraries were used to construct a *K*-mer distribution map to estimate the genome size and heterozygosity of *C. lanceolata*. As shown in **Supplementary Fig. 1**, the average *K*-mer depth corresponding to the main peak was 39, and the genome size was inferred based on *K*-mer number/*K*-mer depth. In addition, the *K*-mer depth of the small peak on the left side of the main peak was 20. We estimated the *C. lanceolata* genome size as 10.42 Gb by using GenomeScope ^100^. The heterozygosity was estimated using combined the SNP calling results. We compared the 217 Gb Illumine data with the assembled *C. lanceolata* genome using Bwa^101^ software to call SNPs for the determination of the heterozygosis SNPs number on the genome. Then it was divided by the genome size to obtain an estimate rate of heterozygosity. Finally, a total of 77,334,119 SNPS were obtained, among which the number of heterozygous SNP was 77,050,655 and the number of homozygous SNPS was 283,464, so the heterozygosity of Chinese fir genome was 0.69%.

### Genome assembly

The assembly of the *C. lanceolata* genome involved three main steps. First, Canu v1.5 (available at https://github.com/marbl/canu)^102^ was used to correct the error of the PacBio clean data. Canu selects longer seed reads (genomeSize=1000000000’ and ‘corOutCoverage=50’), detects the clean read overlaps through the high-sensitive overlapper MHAP (mhap-2.1.2, option ‘corMhapSensitivity=low/normal/high’), and performs an error correction through the falcon_sense method (option ‘correctedErrorRate=0.025’). In the next step, error-corrected reads were trimmed using unsupported bases and hairpin adapters to obtain their longest supported range with the default parameters. In the last step, Canu generated the draft assembly using the longest 80 coverage trimmed reads. In addition, WTDBG2 (https://github.com/ruanjue/wtdbg)^103^ was used to assemble the draft assembly reads. WTDBG2 first generated a draft assembly with the parameter ‘wtdbg -i pbreads.fasta -t 64 -H -k 21 -S 1.02 -e 3 -o wtdbg’. Error-corrected reads from Canu were then used to obtain a better draft assembly performance. The consensus draft assembly results were obtained with the parameter ‘wtdbg-cns -t 64 -i wtdbg.ctg.lay -o wtdbg.ctg.lay.fa -k 15’. Finally, Illumina data were used to perform three rounds of polishing on the consensus draft assembly results through Pilon v1.22 (https://github.com/broadinstitute/pilon)^104^. The first polishing adopted a quiver/arrow algorithm using SMS data with 40 threads. The second polishing adopted the Pilon algorithm using Illumina data with the parameters ‘--mindepth 10 --changes --threads 4 --fix bases’. The completeness of *C. lanceolata* genome assembly was tested against the embryophyta_odb10 lineage (n = 1375) using Benchmarking Universal Single-Copy Orthologues (BUSCO v4.0.1)^105^. To assess the integrity of the assembled genome, Bwa^101^ was used to compare the short sequence obtained through Illumina sequencing with the assembled genome.

### Hi-C library construction and chromosome assembly

Hi-C fragment libraries ranging from 300–700 bp insert size were constructed as described by Rao^106^ *et al*. Based on sequencing using synthesis technology, the Illumina high-throughput sequencing platform was used to sequence Hi-C libraries. The sequencing read length was PE150. After filtering the adapter sequence of raw reads and the low-quality PE reads, we obtained 638.73 Gb clean Hi-C reads, accounting for 62.23× the estimated genome size (**Supplementary Table 26**). The clean Hi-C reads were first truncated at the putative Hi-C junctions. The resulting trimmed reads were aligned to the assembly results with the BWA aligner^104^. Only uniquely aligned read pairs whose mapping quality was more than 20 remained for further analysis. Invalid read pairs, including Da gling-end and self-cycle, re-ligation, and dumped products, were filtered using HiC-Pro v2.8.1^107^. Most (91.98%) of unique mapped read pairs were valid interaction pairs and were used for clustering, or for sorted and orientated scaffolds onto chromosomes by LACHESIS^108^ (**Supplementary Table 27**). The final pseudo-chromosomes were constructed after manual adjustment. To evaluate the result of the Hi-C assembly, we constructed an interaction heat map of the Hi-C assembly chromosome.

### Gene prediction and annotation

Three independent methods were used to predict protein-coding genes: *de novo*, homology-based, and transcriptome-based prediction. Genscan v3.1^109^, Augustu v3.1^110^, Glimmer HMM v3.0.4^111^, Gene ID v1.4^112^, and SNAP v 2006-07-28 (http://homepage.mac.com/iankorf)^113^ with default parameters were applied for the *de novo* gene prediction. Homologous proteins from six known whole-genome sequences (*Arabidopsis*, *Ginkgo*, *G. montanum*, *P. abies*, *Populus trichocarpa*, and *P. taeda*) were aligned to the *C. lanceolata* genome sequence using GeMoMa v1.3.1^114^ with default parameters. Hisat v2.0.4^115^ and Stringtie v1.2.3^116^ default parameters were used to assemble the transcriptome data. TransDecoder v2.0 and GeneMarkS-T v5.1^117^ with default parameters were subsequently applied for gene prediction. PASA v2.0.2^118^ with default parameters was used to predict the Unigene sequences assembled based on transcriptome data and full-length transcripts assembled based on full-length transcriptome data. All acquired results were combined and revised using EVM v1.1.1^119^ and PASA v2.0.2. The completeness of *C. lanceolata* genome annotated was tested using BUSCO v4.0^105^.In addition, the protein-coding gene annotation was performed using BLAST v2.2.31 (1e-5)^120^ against the NR, EuKaryotic Orthologous Groups (KOG), GO, Translated European Molecular Biology Laboratory, and KEGG databases.

### Identification of non-coding RNA, pseudogene, and repetitive sequences

Non-coding RNAs mainly include RNA with a variety of known functions, such as microRNA (miRNA), ribosomal RNA (rRNA), and transfer RNA (tRNA). The Rfam database v12.1^121^ and miRBase database v21 together with Infernal v1.1.1 (http://infernal.janelia.org/)^122^ (1e-5) were used to predict rRNA and miRNA, respectively. The tRNAs were predicted using tRNAs can-SE 1.3.1^123^ with the option ‘-E -H’. Pseudogene homolog sequences were blasted using GenBlastA v1.0.4 (-e 1e-5)^124^. The non-mature termination codes and frameshift mutations were identified using GeneWise v2.4.1 (-both -pseudo)^125^.

Repeat sequences for *C. lanceolata* were predicted as follows. Initially, a *de novo* repeat library was constructed with LTR_FINDER v 1.06 (http://tlife.fudan.edu.cn/ltr_finder/)^126^, Repeat Scount v1.0.5, and PILER-DF v 2.4^127^ with default parameters. The database was classified by PASTEClassifier v1.0^128^ before being combined to build a new repeat database using the RepBase v19.06 database^129^ (http://www.girinst.org/repbase). RepeatMasker v. 4.0.5 (http://www.repeatmasker.org/RepeatModeler/)^130^ was used to align sequences and to screen repeats, including simple repeats, satellites, and low-complexity repeats, using the set parameters ‘-nolow -no_is -norna -engine wublast -qq -frag 20000’. The timing of LTR insertion was estimated using LTR retriever^131^.

### Ortholog detection with OrthoMCL

The amino acid and nucleotide sequences of 19 representative plant species were downloaded from various sources: *A. thaliana*, *Oryza sativa*, *S. moellendorffii*, *Physcomitrella patens* from Phytozome (https://phytozome.jgi.doe.gov/)^132^; *Cinnamomum kanehirae*, *P. taeda, Anthoceros angustus*, *G. montanum*, *W. mirabilis*, *C. panzhihuaensis*, and *T. wallichiana* from NCBI (https://www.ncbi.nlm.nih.gov/genome); *A. alba* and *P. abies* from the Plant Genome Integrative Explorer Resource (http://plantgenie.org/)^133^; Ginkgo from GigaDB^134^; *Amborella* from Ensembl plants; *Nymphaea tetragona* from Genome Warehouse (https://bigd.big.ac.cn/gwh/)^135^; and *Salvinia cucullata* and *Azolla filiculoides* from FernBase (www.fernbase.org.)^136^ Gene families or orthologous groups of these species and *C. lanceolata* were identified using OrthoMCL v1.4 (http://orthomcl.org/orthomcl/)^137^. In addition, we performed KEGG and GO enrichment analyses on the unique gene families in the *C. lanceolata* genome.

### Phylogenetic reconstruction

Orthology analysis revealed the absence of single-copy gene families in these 19 species. Thus, we extracted the genes of the pan-single-copy gene family (i.e., single-copy gene family found in at least 50% of the species) to construct a phylogenetic tree. We initially obtained 27 pan-single-copy gene families. In addition, blastp was used to align multiple-copy gene families in a certain species with single-copy genes in other species. The gene with the best align result was selected as the single-copy gene of the species. Finally, 58 single-copy families were obtained.

MUSCLE v3.8.31 (http://www.drive5.com/muscle/)^138^ was used to align the amino acid sequences of these single-copy orthologs. According to the correspondence of the codons, the amino acid alignment was converted to nucleic acid multiple sequence alignment. After filtering the results of multiple sequence alignments thought Trimal^139^, phylogenetic trees were constructed using RAxML^140^ via concatenation and ASTRAL methods based on nucleic acid sequences and amino acid sequences, respectively. For nucleic acid and amino acid sequences, the parameter was set to -m GTRGAMMA and -m PROTGAMMAJTT, respectively.

In addition, to avoid LBA artifacts, we further selected the first two bases of the codon for nucleotide to constructed concatenated and astral trees and used PhyloBayes and select ‘cat model’ that can effectively avoid LBA artifacts to construct the Bayesian phylogenetic tree.

### Estimation of divergence time

The divergence time of each tree node was inferred using MCMCtree of the PAML4.9 package^141^ (option: correlated molecular clock, JC69 model; the rest is the default). The nucleic acid replacement model was the GTR model, and the molecular clock model the independent rate model. The MCMC process included 100,000 burn-in iterations and 1,000,000 sampling iterations (one sample every 100 iterations). The same parameters were executed twice to obtain more stable results. The phylogeny was calibrated using various fossil records or molecular divergence estimates by placing soft bounds at the split node of *G. montanum* – Ginkgo (230–282 Ma), *A. thaliana* –*G. montanum* (289–330 Ma), *A. angustus* –*G. montanum* (392–422 Ma), *and P. patens* –*G. montanum* (450–514 Ma).

### Gene family expansion and contraction

Based on phylogenetic tree, gene family expansion and loss were inferred using CAFÉ 4.2 (https://github.com/hahnlab/CAFE)^142^. We performed functional enrichment analysis on the genes of the significant expansion and contraction gene family in the *C. lanceolata* genome. Since there are fewer genes with significant contraction and no functions are enriched, the function annotation results of these genes are listed.

### Analysis of genome synteny and WGD

The genes in the collinearity fragments maintained a high degree of conservation during the evolution of the species. After shielding some of the TE-related genes, Blastp v2.2.26 (www.ncbi.nlm.nih.gov/*BLASTP*) was used to perform all-vs-all alignment of gymnosperms plants, including *G. biloba*, *C. panzhihuaensis*, *C. lanceolata*, *T. wallichiana*, *W. mirabilis*, and *G. montanum*. Then, MCScanX^143^ (except for -b=1/2 setting intra-species analysis, the rest are the default parameters) was used to conduct a collinear analysis of between these genomes.

We used substitutions per synonymous site (*K*s) distribution analysis to estimate WGD events in the gymnosperm genomes. Diamond was used to conduct self-alignment of the protein sequences of these species genomes and to extract the mutual optimal alignment in the alignment results. Finally, Codeml in the PAML package was used to calculate the *K*s values between gene pairs in each gene family. For *K*s value calculations between species, we used MCScanX (parameter: -a-e 1e-5-s 5) to find the collinear blocks between two species and then calculate the *K*s values of the orthologous gene pairs in each collinear block using Codeml in the PAML package. In addition, the Whale model-based phylogenetic method^144^ was used to test the occurrence of ancient polyploidy event that *C. lanceolata* may experience.

In addition, the model-based phylogenetic software WHALE was used to test the validity of the above inferred WGD events^144^. The OrthoFinder^145^ (v2.3.3) was used to infer the orthologous gene families with parameters set as default. Gene families which don’t have at least one gene from both clades at the root or have a family size exceeding 2 times the median of the square root of the family size based on a Poisson outlier criterion were filtered out. An amino acid multiply sequence alignment (MSA) was obtained using the PRANK^146^ for each gene family and the resulting MSA was then used as input for the Markov Chain Monte Carlo (MCMC) analysis in mrbayes^147^ (v.3.2.6) to sample from the posterior probability distribution. The Aamodelpr was set as fixed (LG) and the rates was set as gamma-distributed variation approximated using 4 categories. The sample frequency was set as 10 and the number of generations was set as 110000 to get in total 11000 posterior samples. The ALEobserve was then used to construct the conditional clade distribution (CCD) containing marginal clade frequencies with a burnin as 1000 based on the 11000 posterior samples for each gene family. The topology of species tree is set as shown in **Fig. 1** and the divergence time were retrieved from TimeTree^148^.

The duplication-loss (DL)+WGD model under critical and relaxed branch-specific rates were implemented for the inference of the significance and corresponding retention rates of the assumed WGD events in the manner of Bayesian inference (BI). In the critical branch-specific DL+WGD model, the prior η which denotes the parameter of the geometric prior distribution on the number of genes at the root was set to follow a truncated univariate Beta distribution with shape parameters as (3,1) in the interval [0.01, 0.99], the prior r which denotes the mean of the branch rates distribution was set to follow a flat distribution, the prior σ which denotes the deviation of the branch rates distribution was set to follow an exponential distribution with scale 0.1, the λ which denotes the duplication rate of each branch was set to follow a multivariate normal distributions for each branch and the loss rate μ was set to be equal as λ, while in the relaxed branch specific model, the λ and μ are independent with the rates variation parameter τ set to follow an exponential distribution with scale 1. In the model estimating the branch-specific duplication and loss rates, the λ and μ are set to follow a normal distribution with mean as 0 and standard deviation as 5 in log for each branch, independently, with all branch lengths set to 1 and no WGD nodes. The Bayes Factor was calculated using the “bfact.jl” script within the public github repository of WHALE to measure the strength of evidence in favor of the assumed WGD models using the Savage-Dickey density ratio.

### Transcriptomic data and analysis

The material for transcriptome sequencing came from four sources. The first was young tissues of the vegetative organs (leaf, phloem, stem, and root) and reproductive organs (homozygous female gametophytes, male cone, female cone, seed scale, and bract scale) of genome sequencing materials. The second was germinating and astringent seeds at four stages of (105, 115, 125, and 135 d) (**Supplementary Note 1**). The third was drought-tolerant and non-drought-tolerant leaves of *C. lanceolata* seedlings (**Supplementary Note 5**). The fourth was leaves of two-year-old *C. lanceolata* 228 and 226 clone seedlings grow normally in a shade-tolerant environment and the upper, middle, and lower leaves of adult *C. lanceolata*. This adult tree was approximately 30 m in height. The approximate height of self-pruning (HSP) was 12 m. The branches at the HSP were selected as the lowest samples (OD group), while at the top of the plants were selected as the highest samples (OU group). The branches growth near the midpoint of highest and lowest were selected as the middle layer samples (OM group).

For library construction, a total of 1.5 μg RNA was prepared, and libraries were generated using NEBNextR UltraTM Directional RNA Library Prep Kit for IlluminaR (NEB). The index codes were added to attribute sequences to each sample. Clustering of the index-coded samples was performed on an acBot Cluster Generation System using TruSeq PE Cluster Kitv3-cBot-HS (Illumina) according to the manufacturer’s instructions. After cluster generation, the library preparations were sequenced on an Illumina HiSeq platform and paired-end reads were generated. Raw data of fastq format were first processed through in-houseperl scripts. In this step, clean data were obtained by removing reads containing adapter, ploy-N, and low-quality reads from raw data. StringTie (1.3.1) was used to calculate the FPKMs of coding genes in each sample. Gene fragments per kilobase of transcript per million mapped reads (FPKMs) was computed by summing the FPKMs of transcripts in each gene group.

### MADS-box gene family analysis

The HMM profiles of MADS (PF00319) were obtained from Pfam (http://pfam.xfam.org/)^149^. The MADS-box candidate gene protein was separately searched using HMMER 3.2.1 (http://hmmer.org)(with default parameters) and BLASTP (E-value of e^−5^) methods. Subsequently, the domains of all the MADS-box gene candidate sequences were identified using SMART (http://smart.embl-heidelberg.de/)^150^. MADS-box classification was based on sequence similarity searches of identified MADS-box genes from *Arabidopsis* and *Amborella*^49^. All the candidate MADS-box genes were aligned using MAFFT^151^. The phylogenetic tree was constructed using FastTree v2.1.10^152^ and edited in Figtree v1.4.4 (http://tree.bio.ed.ac.uk/software/figtree/).

### Evolutionary analysis of CAM genes

CAM genes were identified by searching the InterProScan^153^ results of all predicted *C. lanceolata* proteins. *Arabidopsis* CAM gene was downloaded from TAIR^154^. We identified orthologs of the *C. lanceolata* CAM genes in the genomes of other *Arabidopsis* using a reciprocal best hit strategy implemented with NCBI BLAST (https://blast.ncbi.nlm.nih.gov/Blast.cgi) and custom scripts. Then, we manually deleted the short reading frame and suspicious genes caused by the relaxed annotation of InterProScan. The coding sequences of each gene family were aligned using MUSCLE implemented in MEGA5^155^. A gene tree was constructed with MEGA5 using the maximum likelihood for each gene family.

### Identification of the shade avoidance-related gene

We downloaded all the shade avoidance-related genes of *Arabidopsis* from TAIR (https://www.arabidopsis.org) as the query sequences. TBLASTN (NCBI Blast v. 2.2.23)^156^ was used to align the query sequences against each genome sequence (E-value cutoff < 1e−10). Subsequently, the domains of shade avoidance-related sequences were identified using SMART (http://smart.embl-heidelberg.de/), and putative shade avoidance-related genes were predicted using the NCBI BLAST (https://blast.ncbi.nlm.nih.gov/Blast.cgi). For gene families that need to build a gene tree, the gene tree construction methods were similar to those described in the “Evolutionary analysis of CAM genes” section.

The identification method of the *C. lanceolata* AUX/IAA gene family was similar to that of the “MADS-box gene family analysis” section. The HMM profiles of AUX/IAA (PF02309) were obtained from Pfam (http://pfam.xfam.org/)^147^. AUX/IAA classification was based on sequence similarity searches of identified *Arabidopsis* AUX/IAA genes^157^.

### *Cunninghamia lanceolata* aluminum stress experiment

Astringent seeds were ground into powder and used to make three different concentrations of extracts. AlCl3·6H2O was used as the source of aluminum ions. Three concentration gradients were set. The healthy seeds of the same mother plant were collected, germinated, and cultured for 14 days. Seedlings with similar root length, seedling height, and leaf growth potential were selected for the aluminum stress experiment. Different concentrations of astringent water extracts and aluminum were used as solvents for cultivating seedlings. A total of three sets of experiments were designed, and the reagents were added with nutrients according to the Hoagland nutritional formula. Three replicates were set for each treatment group, and 30 seedlings were cultivated for each replicate, and all the seedlings were cultivated in the plant growth chamber. After 30 days of cultivation, we measured the tip aluminum ion concentration, relative root elongation, tip malondialdehyde, and superoxide dismutase of the seedling roots. See **Supplementary Note 2** for details.

## Data Availability

PacBio whole-genome sequencing data, Illumina data, and genome assembly sequences have been deposited to the NCBI Sequence Read Archive (SRA) as Bioproject PRJNA668674.

## Acknowledgements

We acknowledge the support received through the Innovative Research and Constructive Platform Funds of Fujian Agriculture and Forestry University (118-612014032 and KLE18010A), awarded to S-Z. L.; and Forestry Peak Discipline Construction Project of Fujian Agriculture and Forestry University (72202200205), awarded to Z.-J. L.

## Author contributions

S.-Z.L & Z.-J.L. managed the project; Y.V.d.P., R.M., H.-K.Z., S.-Z.L., Y.C, W.-H.S., C.W., and Z.L planned and coordinated the project; S.-Z.L., Z.-J.L., Y.C, W.-H.S., C.W., and Z.L. wrote the manuscript; W.-H.S., Y.C, C.W., Y.-C.X., L.Y., Y.Z., M.-M.L., W.-C.T. and Y.-C.X. collected and sequenced the plant material; H.-K.Z., Q.-G. Z., W.-H.S., C.W., L.X., D.-K.L., D.-Q.C., and L.N. assembled and annotated the genome; S.-Z.L., S.-Z.L., Z.L., H.-C.C., J.-Y.W., Z.-W.W., Z.-W.L., performed gene family clustering and comparative phylogenomics; Y.C., D.-Y.Z., X.Y., D.-K.L., G.-Z.C., J.H., M.-Z.H., X.Z., W.-Y.Z., F.-L.W., Y. L., Q. Z, executed transcriptome sequencing and analysis; W.-H.S., W.-C.T., Y.-C.X., S.-S.X., A.-Q.L., Y.-Q.L., S.-N.R., B.L. conducted the evolution of reproductive organs analysis; C.W., X.-Q., M., F.-P., Z., Y., L.Y., Y.Z., M.-M.L., J.-J.Z., Z.-M.H., P.-F.W., K.-M.L., conducted the formation of astringent seeds analysis; S.-Z.L., C.-M.J., S.-R.L., S.-B.L., Y.-Q.Y., Z.-H.M., G.-C.D., G.-Q.C., S.X., J.Z., conducted the shade tolerance of seeds analysis. All authors read and approved the manuscript.

## Additional information

Methods and Supplementary information is available in the online version of the paper.

## Reference

1. Bowe, L. M. et al. Phylogeny of seed plants based on all three genomic compartments: extant gymnosperms are monophyletic and Gnetales’ closest relatives are conifers. Proc. Natl. Acad. Sci. USA. 97, 4092–4097 (2000).

2. Chaw, S. M., et al. Seed plant phylogeny inferred from all three plant genomes: monophyly of extant gymnosperms and origin of gnetales from conifers. Proc. Natl. Acad. Sci. USA. 97, 4086–4091 (2000).

3. Xi, Z. et al. Phylogenomics and coalescent analyses resolve extant seed plant relationships. PLoS ONE 8, e80870 (2013).

4. Rothwell, G. W. & Scheckler, S. E. In Origin and Evolution of Gymnosperms (ed. Beck, C. B.) 85–134, New York, Columbia University Press (1988).

5. Savard, L. et al. Chloroplast and nuclear gene sequences indicate late Pennsylvanian time for the last common ancestor of extant seed plants. Proc. Natl. Acad. Sci. USA. 91, 5163–5167 (1994).

6. Stewart, W. N. & Rothwell, G. W. Paleobotany and The Evolution of Plants. Cambridge University Press (1993).

7. Lu, Y. et al. Phylogeny and divergence times of gymnosperms inferred from single-copy nuclear genes. PLoS ONE 9, e107679 (2014).

8. Wang, X. Q. & Ran, J. H. Evolution and biogeography of gymnosperms. Mol. Phy. Evol. 75, 24–40 (2014).

9. Mathews, S. Phylogenetic relationships among seed plants: Persistent questions and the limits of molecular data. Am. J. Bot. 96, 228–236 (2009).

10. Wickett, N. J. et al. Phylotranscriptomic analysis of the origin and early diversification of land plants. Proc. Natl. Acad. Sci. USA. 111, e4859–4868 (2014).

11. Ruhfel, B. R. et al. From algae to angiosperms-inferring the phylogeny of green plants (Viridiplantae) from 360 plastid genomes. BMC Evol. Biol. 14, 23 (2014).

12. Wan, T. et al. A genome for gnetophytes and early evolution of seed plants. Nat. Plants 4, 82–89 (2018).

13. Li, Z. et al. Single-copy genes as molecular markers for phylogenomic studies in seed plants. Genome Biol. Evol. 9, 1130–1147 (2017).

14. Ahuja, M. R. & Neale, D. B. Evolution of genome size in gonifers. Silvae. Genet. 54, 126–137 (2005).

15. Morse, A. M. et al. Evolution of genome size and complexity in Pinus. PLoS ONE 4, e4332 (2009).

16. Kovach, A. et al. The *Pinus taeda* genome is characterized by diverse and highly diverged repetitive sequences. BMC Genom. 11, 420 (2010).

17. Mackay, J. et al. Towards decoding the conifer giga-genome. Plant Mol. Biol. 80, 555–569 (2012).

18. De La Torre, A. R. et al. Contrasting rates of molecular evolution and patterns of selection among gymnosperms and flowering plants. Mol. Biol. Evol. 34, 1363–1377 (2017).

19. Nystedt, B. et al. The *Norway spruce* genome sequence and conifer genome evolution. Nature 497, 579–584 (2013).

20. Birol, I. et al. Assembling the 20 Gb white spruce (*Picea glauca*) genome from whole-genome shotgun sequencing data. Bioinformatics 29, 1492–1497 (2013).

21. Wegrzyn, J. L. et al. Insights into the loblolly pine genome: characterization of BAC and fosmid sequences. PLoS ONE 8, e72439 (2013).

22. Wegrzyn, J. L. et al. Unique features of the loblolly pine (*Pinus taeda* L.) megagenome revealed through sequence annotation. Genetics 196, 891–909 (2014).

23. Zimin, A. et al. Sequencing and assembly of the 22-gb loblolly pine genome. Genetics 196, 875–890 (2014).

24. Niu, S. et al. The Chinese pine genome and methylome unveil key features of conifer evolution. Cell 185, 204–217 (2022).

25. Sun, C. et al. The *Larix kaempferi* genome reveals new insights into wood properties. J. Integr. Plant Biol. doi: 10.1111/jipb.13265 (2022).

26. Guan, R. et al. Draft genome of the living fossil *Ginkgo biloba*. GigaScience 5, 49 (2016).

27. Zhao, Y. P. et al. Resequencing 545 ginkgo genomes across the world reveals the evolutionary history of the living fossil. Nat. Commun. 10, 4201 (2019).

28. Xiong, X. et al. The *Taxus* genome provides insights into paclitaxel biosynthesis. Nat. Plants 7, 1026–1036 (2021).

29. Liu, Y. et al. The *Cycas* genome and the early evolution of seed plants. Nat. Plants 8, 389–401(2022).

30. Wan, T. et al. The *Welwitschia* genome reveals a unique biology underpinning extreme longevity in deserts. Nat. Commun. 12, 4247 (2021).

31. Farjon, A. A handbook of the world’s conifers. BRILL, Leiden, Boston RBG 1, 1–526 (2010).

32. Forest, F. et al. Gymnosperms on the EDGE. Sci. Rep. 8, 6053 (2018).

33. Yu, X. T. Culture of Chinese Fir. Fujian Science & Technology Press, 16–19 (1997).

34. Lin, Y. D. & Zhang, C. X. The brief history of cultivating *Cunninghamia lanceolata* in China. J. Dialect. Nat. 29, 79–82 (2007).

35. Zhang, X. et al. Estimated biomass carbon in thinned *Cunninghamia lanceolata* plantations at different stand-ages. J. Forestry Res. 32, 1–13 (2021).

36. Xu, J. & Shi, J. S. C banding and fluorescent banding pattern of the chromosome of *Cuninghamia lanceolate*. Mol. Plant Breeding 5, 515–520 (2007).

37. Vinogradov, A. E. Intron-genome size relationship on a large evolutionary scale. J. Mol. Evol. 49, 376–384 (1999).

38. Neale, D. B. et al. Decoding the massive genome of loblolly pine using haploid DNA and novel assembly strategies. Genome Biol. 15, R59 (2014).

39. Stull, G.W. et al. Gene duplications and phylogenomic conflict underlie major pulses of phenotypic evolution in gymnosperms. Nat. Plants 7, 1015–1025 (2021).

40. Bergsten J. A review of long-branch attraction. Cladistics. 21:163–193 (2005).

41. Ran, J. H. et al. Phylogenomics resolves the deep phylogeny of seed plants and indicates partial convergent or homoplastic evolution between Gnetales and angiosperms. Proc. Biol Sci. 285, 20181012 (2018)

42. Leebens-Mack, J. H. et al. One thousand plant transcriptomes and the phylogenomics of green plants. Nature 574, 679–685 (2019).

43. Van de Peer, Y. et al. Polyploidy: an evolutionary and ecological force in stressful times. Plant Cell 33, 11–26 (2021).

44. Jin, W. T. et al. Phylogenomic and ecological analyses reveal the spatiotemporal evolution of global pines. Proc. Natl. Acad. Sci. USA. 118, e2022302118 (2021)

45. Sensalari, C. et al. ksrates: positioning whole-genome duplications relative to speciation events in KS distributions. Bioinformatics 18, btab602 (2021)

46. Zwaenepoel, A. & Van de Peer, Y. Inference of ancient Whole-Genome Duplications and the evolution of gene duplication and loss rates. Mol. Biol. Evol. 36,1384–1404 (2019).

47. Jiao, Y. et al. Ancestral polyploidy in seed plants and angiosperms. Nature, 473, 97–100 (2011).

48. Van de Peer, Y, et al. The evolutionary significance of polyploidy. Nat. Rev. Genet. 18, 411–424 (2017).

49. Amborella Genome Project. The *Amborella* genome and the evolution of flowering plants. Science 342,1241089 (2013).

50. Qin, L. et al. Insights into angiosperm evolution, floral development and chemical biosynthesis from the *Aristolochia fimbriata* genome. Nat. Plants 7, 1239–1253 (2021).

51. Crane, P. R. et al. The origin and early diversification of angiosperms. Nature 374, 27–33 (1995).

52. Melzer, R. et al. The naked and the dead: the ABCs of gymnosperm reproduction and the origin of the angiosperm flower. Semin. Cell Dev. Biol. 21, 118–128 (2010).

53. Chen, F. et al. Evolutionary analysis of MIKC(c)-Type MADS-Box genes in gymnosperms and angiosperms. Front. Plant Sci. 8, 895 (2017).

54. Zhang, L. et al. The water lily genome and the early evolution of flowering plants. Nature 577, 79–84 (2020).

55. Nam, J. et al. Type I MADS-box genes have experienced faster birth-and-death evolution than type II MADS-box genes in angiosperms. Proc. Natl. Acad. Sci. USA. 101, 1910–1915 (2004).

56. Mouradov, A. et al. Family of MADS-Box genes expressed early in male and female reproductive structures of monterey pine. Plant Physiol. 117, 55–62 (1998).

57. Dreni, L. & Zhang, D. Flower development: the evolutionary history and functions of the AGL6 subfamily MADS-box genes. J. Exp. Bot. 67,1625–38 (2016).

58. Theissen, G. et al. MADS-domain transcription factors and the floral quartet model of flower development: linking plant development and evolution. Development 143, 3259–3271 (2016).

59. Levin, J. Z. & Meyerowitz, E. M. *UFO*: an *Arabidopsis* gene involved in both floral meristem and floral organ development. Plant Cell 7, 529–548 (1995).

60. Denay, G. et al. A flower is born: an update on *Arabidopsis* floral meristem formation. Curr. Opin. Plant Biol. 35, 15–22 (2017).

61. Chae, E. et al. An *Arabidopsis* F-box protein acts as a transcriptional co-factor to regulate floral development. Development 135, 1235–1245 (2008).

62. Lenhard, M. et al. Termination of stem cell maintenance in *Arabidopsis* floral meristems by interactions between *WUSCHEL* and *AGAMOUS*. Cell 105, 805–814 (2001).

63. Hedman, H. et al. Analysis of the WUSCHEL-RELATED HOMEOBOX gene family in the conifer *Picea abies* reveals extensive conservation as well as dynamic patterns. BMC Plant Biol. 13, 89 (2013).

64. Van der Graaff, E. et al. The *WUS* homeobox-containing (*WOX*) protein family. Genome Biol. 10, 248 (2009).

65. Rebocho, A. B. et al. Role of EVERGREEN in the development of the cymose petunia inflorescence. Dev. Cell 15, 437–447 (2008).

66. Breuninger, H. et al. Differential expression of WOX genes mediates apical-basal axis formation in the *Arabidopsis* embryo. Dev. Cell 14, 867–876 (2008).

67. Haecker, A. et al. Expression dynamics of WOX genes mark cell fate decisions during early embryonic patterning in *Arabidopsis thaliana*. Development 131, 657–668 (2004).

68. Wu, X. et al. Combinations of *WOX* activities regulate tissue proliferation during *Arabidopsis* embryonic development. Dev. Bio. 309, 306–316 (2007).

69. Palovaara, J. et al. Comparative expression pattern analysis of WUSCHEL-related homeobox 2 (*WOX2*) and *WOX8*/*9* in developing seeds and somatic embryos of the gymnosperm *Picea abies*. New Phytol. 188, 122–135 (2010).

70. Yu, X. Y. & Li, P. Studies of the abortive seed and its constituents in *Cunninghamia lanceolata* (Lamb.) Hook. Acta. Bot. Boreal. Occident. Sin. 9, 252–256 (1989).

71. Blanco, J. A. et al. Impacts of enhanced nitrogen deposition and soil acidification on biomass production and nitrogen leaching in Chinese fir plantations. Can. J. Forest. Res. 42, 1296–1298 (2012).

72. Ye, Y. Q. et al. Effects of aluminum stress on growth, photosynthetic characteristics and chloroplast ultrastructure in leaves of *Cunninghamia lanceolata* seedlings. Journal of Northeast Forsetry University 48, 7–11,16 (2020).

73. Bandowe, B. A. et al. Sources and fate of polycyclic aromatic compounds (PAHs, oxygenated PAHs and azaarenes) in forest soil profiles opposite of an aluminium plant. Sci. Total Environ. 630, 83–95 (2018).

74. Hubova, P. et al. Behaviour of aluminium in forest soils with different lithology and herb vegetation cover. J. Inorg. Biochem. 181, 139–144 (2018).

75. Bradova, M. et al. The variations of aluminium species in mountainous forest soils and its implications to soil acidification. Environ. Sci. Pollut. Res. 22, 16676–16687 (2015).

76. Lin, X. Q. et al. Optimization of ultrasonic-assisted extraction process for polyphenols from the seeds of Chinese fir. J. Forest Environ. 37, 47–53 (2017).

77. Meinke, D. W. Genome-wide identification of *EMBRYO*-*DEFECTIVE* (*EMB*) genes required for growth and development in *Arabidopsis*. New Phytol. 226, 306–325 (2020).

78. Dutta, A. et al. Hypersensitive response-like lesions 1 codes for *AtPPT1* and regulates accumulation of ROS and defense against bacterial pathogen *Pseudomonas syringae* in *Arabidopsis thaliana*. Antioxid. Redox. Sign. 22, 785–796 (2015).

79. Sharma, P. et al. Reactive oxygen species, oxidative damage, and antioxidative defense mechanism in plants under stressful conditions. J. Bot. 2012, 1–26 (2012).

80. Inostroza-Blancheteau, C. et al. Molecular and physiological strategies to increase aluminum resistance in plants. Mol. Biol. Rep. 39, 2069–2079 (2012).

81. Yan, L. F. et al. Responses of Non-structural Carbohydrates in above ground tissues/organs and root to shading and light restoration in *Cunninghamia lanceolata* saplings. Acta. Bot. Boreal. Occident. Sin. 40, 0311–0318 (2020).

82. Smith, H. Physiological and ecological function within the phytochrome Family. Annu. Rev. Plant Physi. Plant Mol. Biol. 46, 289–315 (1995).

83. Cagnola, J. I. et al. Stem transcriptome reveals mechanisms to reduce the energetic cost of shade-avoidance responses in tomato. Plant Physiol. 160,1110–19 (2012)

84. Mathews, S. Phytochrome-mediated development in land plants: red light sensing evolves to meet the challenges of changing light environments. Mol. Ecol. 15, 3483–3503 (2006).

85. Casal, J. J. Photoreceptor signaling networks in plant responses to shade. Annu. Rev. Plant Biol. 64, 403–427 (2013).

86. Leivar, P. et al. Dynamic antagonism between phytochromes and PIF family basic helix-loop-helix factors induces selective reciprocal responses to light and shade in a rapidly responsive transcriptional network in *Arabidopsis*. Plant Cell 24, 1398–1419 (2012).

97. Sellaro, R. et al. Diurnal dependence of growth responses to shade in *Arabidopsis*: role of hormone, clock, and light signaling. Mol. Plant 5, 619–628 (2012).

88. Li, L. et al. Linking photoreceptor excitation to changes in plant architecture. Genes Dev. 26, 785–790 (2012).

89. Lorrain, S. et al. Phytochrome-mediated inhibition of shade avoidance involves degradation of growth-promoting bHLH transcription factors. Plant J. 53, 312–323 (2008).

90. Wu, L. & Yang, H. Q. CRYPTOCHROME 1 is implicated in promoting R protein-mediated plant resistance to *Pseudomonas syringae* in *Arabidopsis*. Mol. Plant 3, 539–548 (2010).

91. Christie, J. M. Phototropin blue-light receptors. Annu. Rev. Plant Biol. 58, 21–45 (2007).

92. Sakai, T. et al. *Arabidopsis* nph1 and npl1: blue light receptors that mediate both phototropism and chloroplast relocation. Proc. Natl. Acad. Sci. USA. 98, 6969–6974 (2001).

93. Davis, P. A. et al. Changes in leaf optical properties associated with light-dependent chloroplast movements. Plant Cell Environ. 34, 2047–2059 (2011).

94. Brown, B. A. et al. A UV-B-specific signaling component orchestrates plant UV protection. Proc. Natl. Acad. Sci. USA. 102, 18225–18230 (2005).

95. Kliebenstein, D. J. et al. *Arabidopsis UVR8* regulates ultraviolet-B signal transduction and tolerance and contains sequence similarity to human regulator of chromatin condensation 1. Plant Physiol. 130, 234–243 (2002).

96. Demkura, P. V. et al. Jasmonate-dependent and -independent pathways mediate specific effects of solar ultraviolet B radiation on leaf phenolics and antiherbivore defense. Plant Physiol. 152, 1084–1095 (2010).

97. Lau, O. S. & Deng, X. W. The photomorphogenic repressors *COP1* and *DET1*: 20 years later. Trends Plant Sci. 17, 584–593 (2012).

98. Heijde, M. & Ulm, R. UV-B photoreceptor-mediated signalling in plants. Trends Plant Sci. 17, 230–237 (2012).

99. Kang, C. Y. et al. Cryptochromes, phytochromes, and *COP1* regulate light-controlled stomatal development in *Arabidopsis*. Plant Cell 21, 2624–2641 (2009).

100. Vurture, G. W. et al. GenomeScope: fast reference-free genome profiling from short reads. Bioinformatics 33, 2202–2204 (2017).

101. Li, H. & Durbin, R. Fast and accurate short read alignment with Burrows-Wheeler transform. Bioinformatics 25, 1754–1760 (2009).

102. Koren, S. et al. Canu: scalable and accurate long-read assembly via adaptive k-mer weighting and repeat separation. Genome Res. 27, 722–736 (2017).

103. Ruan, J. & Li, H. Fast and accurate long-read assembly with Wtdbg2. Nat. Methods 17, 155–158 (2020).

104. Walker, B. J. et al. Pilon: an integrated tool for comprehensive microbial variant detection and genome assembly improvement. PLoS ONE 9, e112963 (2014)

105. Simao, F.A. et al. BUSCO: assessing genome assembly and annotation completeness with single-copy orthologs. Bioinformatics 31, 3210–3212 (2015).

106. Rao, S. S. et al. A 3D map of the human genome at kilobase resolution reveals principles of chromatin looping. Cell 159, 1665–1680 (2014).

107. Servant, N. et al. HiC-Pro: an optimized and flexible pipeline for Hi-C data processing. Genome Biol. 16, 259 (2015).

108. Burton, J. N. et al. Chromosome-scale scaffolding of de novo genome assemblies based on chromatin interactions. Nat. Biotechnol. 31, 1119–1125 (2013).

109. Burge, C. & Karlin, S. Prediction of complete gene structures in human genomic DNA. J. Mol. Biol. 268, 78–94 (1997).

110. Stanke, M. & Waack, S. Gene prediction with a hidden Markov model and a new intron submodel. Bioinformatics 19, ii215–225 (2003).

111. Majoros, W. H. et al. TigrScan and GlimmerHMM: two open source ab initio eukaryotic gene-finders. Bioinformatics 20, 2878–2879 (2004).

112. Alioto, T. et al. Using geneid to identify genes. Curr. Protoc. Bioinformatics 64, e56 (2018).

113. Johnson, A. D. et al. SNAP: a web-based tool for identification and annotation of proxy SNPs using HapMap. Bioinformatics 24, 2938–2939 (2008).

114. Keilwagen, J. et al. Using intron position conservation for homology-based gene prediction. Nucleic Acids Res. 44, e89 (2016).

115. Kim, D. et al. HISAT: a fast spliced aligner with low memory requirements. Nat. Methods 12, 357–360 (2015).

116. Pertea, M. et al. Transcript-level expression analysis of RNA-seq experiments with HISAT, StringTie and Ball-gown. Nat. Protoc. 11,1650–1667 (2016)

117. Tang, S. et al. Identification of protein coding regions in RNA transcripts. Nucleic Acids Res. 43, e78 (2015).

118. Haas, B. J. et al. Improving the *Arabidopsis* genome annotation using maximal transcript alignment assemblies. Nucleic Acids Res. 31, 5654–5666 (2003).

119. Haas, B. J. et al. Automated eukaryotic gene structure annotation using EVidenceModeler and the program to assemble spliced alignments. Genome Biol. 9, R7 (2008).

120. Altschul, S. F. et al. Basic local alignment search tool. J. Mol. Biol. 215, 403–410 (1990).

121. Griffiths-Jones, S. et al. Rfam: annotating non-coding RNAs in complete genomes. Nucleic Acids Res. 33(2005).

122. Nawrocki, E. P. et al. Infernal 1.0: inference of RNA alignments. Bioinformatics 25, 1335–1337 (2009).

123. Lowe, T. M. & Eddy, S. R. tRNAscan-SE: a program for improved detection of transfer RNA genes in genomic sequence. Nucleic Acids Res. 25, 955–964 (1997).

124. She, R. et al. GenBlastA: enabling BLAST to identify homologous gene sequences. Genome Res. 19, 143–149 (2009).

125. Birney, E. et al. GeneWise and Genomewise. Genome Res. 14, 988–995 (2004).

126. Zhao, X. & Hao, W. LTR_FINDER: an efficient tool for the prediction of full-length LTR retrotransposons. Nucleic Acids Res. 35, W265–268 (2007).

127. Edgar, R. C. & Myers, E. W. PILER: identification and classification of genomic repeats. Bioinformatics 21 Suppl 1, i152–158 (2005).

128. Hoede, C. et al. PASTEC: an automatic transposable element classification tool. PLoS ONE 9, e91929 (2014).

129. Jurka, J. et al. Repbase Update, a database of eukaryotic repetitive elements. Cytogenet. Genome Res. 110, 462–467 (2005).

130. Tarailo-Graovac, M. & Chen, N. Using RepeatMasker to identify repetitive elements in genomic sequences. Curr. Protoc. Bioinformatics Chapter 4, Unit 4 10 (2009).

131. Ou, S. and Jiang, N. LTR_retriever: a highly accurate and sensitive program for identification of long terminal repeat retrotransposons. Plant Physiol. 176,1410–1422 (2018)

132. Goodstein, D. M. et al. Phytozome: a comparative platform for green plant genomics. Nucleic Acids Res. 40, D1178–1186 (2012).

133. Sundell, D. et al. The plant genome integrative explorer resource: plantgenie.org. New Phytol. 208, 1149–1156 (2015).

134. Sneddon, T. P. et al. GigaDB: announcing the GigaScience database. Gigascience 1, 11 (2012).

135. Chen, M. et al. Genome Warehouse: a public repository housing genome-scale data. Genom. Proteom. Bioinf. 19, 584–589 (2021).

136. Li, F. W. et al. Fern genomes elucidate land plant evolution and cyanobacterial symbioses. Nat. Plants 4, 460–472 (2018).

137. Li, L. et al. OrthoMCL: identification of ortholog groups for eukaryotic genomes. Genome Res. 13, 2178–2189 (2003).

138. Edgar, R. C. MUSCLE: multiple sequence alignment with high accuracy and high throughput. Nucleic Acids Res. 32, 1792–1797 (2004).

139. Capella-Gutierrez, S. et al. Trimal: a tool for automated alignment trimming in large-scale phylogenetic analyses. Bioinformatics 25, 1972–1973 (2009)

140. Stamatakis, A. Using RAxML to infer phylogenies. Curr. Protoc. Bioinformatics 51, 6.14.11–16.14 (2015).

141. Yang, Z. PAML 4: phylogenetic analysis by maximum likelihood. Mol. Biol. Evol. 24, 1586–1591 (2007).

142. De Bie, T. et al. CAFE: a computational tool for the study of gene family evolution. Bioinformatics 22, 1269–1271 (2006).

143. Wang, Y. et al. MCScanX: a toolkit for detection and evolutionary analysis of gene synteny and collinearity. Nucleic Acids Res. 40, e49 (2012).

144. Zwaenepoel, A. & Van de Peer, Y. Inference of ancient Whole-Genome Duplications and the evolution of gene duplication and loss rates. Mol. Biol. Evol. 36, 1384–1404 (2019).

145. Emms, D. M. & S. Kelly. OrthoFinder: phylogenetic orthology inference for comparative genomics. Genome Biol. 20, 238 (2019).

146. Löytynoja, A. & Goldman, N. An algorithm for progressive multiple alignment of sequences with insertions. Proc. Natl. Acad. Sci. USA 102, 10557 (2005).

147. Huelsenbeck, J. P. & Ronquist, F. MRBAYES: Bayesian inference of phylogenetic trees. Bioinformatics 17, 754–755 (2001).

148. Kumar, S. et al. TimeTree: A Resource for timelines, timetrees, and bivergence times. Mol. Biol. Evol. 34, 1812–1819 (2017).

149. Finn, R. D. et al. The Pfam protein families database. Nucleic Acids Res. 38, D211–222 (2010).

150. Letunic, I. et al. Recent improvements to the SMART domain-based sequence annotation resource. Nucleic Acids Res. 30, 242–244 (2002).

151. Katoh, K. & Standley, D. M. MAFFT multiple sequence alignment software version 7: improvements in performance and usability. Mol. Biol. Evol. 30, 772–780 (2013).

152. Price, M. N. et al. FastTree 2–approximately maximum-likelihood trees for large alignments. PLoS ONE 5, e9490 (2010).

153. Jones, P. et al. InterProScan 5: genome-scale protein function classification. Bioinformatics 30, 1236–1240 (2014).

154. Swarbreck, D. et al. The *Arabidopsis* Information Resource (TAIR): gene structure and function annotation. Nucleic Acids Res. 36, D1009–1014 (2008).

155. Tamura, K. et al. MEGA5: molecular evolutionary genetics analysis using maximum likelihood, evolutionary distance, and maximum parsimony methods. Mol. Biol. Evol. 28, 2731–2739 (2011).

156. Kent, W. J. BLAT--the BLAST-like alignment tool. Genome Res. 12, 656–664 (2002).

157. Liscum, E. & Reed, J. W. Genetics of Aux/IAA and ARF action in plant growth and development. Plant Mol. Biol. 49, 387–400 (2002).

