## Supplemental Information for "Chinese fir genome and the evolution of gymnosperms"

**Content**

Supplementary Reference	54

**Supplementary Notes**

**Supplementary Note 1. Observation and determination of the structure of astringent and germinating seeds in different development stages.**

Samples were collected from the *Cunninghamia lanceolata* third-generation seed orchard of Youxi National Forest Farm, Fujian Province, China. The geographical coordinates of the orchard are 25°50 ' – 26°26' N, 117°48 ' – 118°39' E. In 2017, ten healthy individuals with similar height, diameter, and annual astringent seed incidence rate were selected from the same area of the orchard for pollination randomly. The date of the artificial supplementary pollination (March 1) was recorded as 0 d. After 60 d, three cones from each individual were collected every 5 d until the seeds were harvested. We then observed the structures of the astringent seeds and germinating seeds at 95 d, 105 d, 115 d, 125 d, 135 d, 145 d, 155d, 165d, 175d, and 185d using an optical microscope. During observation, the seed coats were quickly and carefully removed and the embryos were examined under 20× using an optical microscope. Based on its color, luster, and hardness, the embryos in different growing stages were judged to be astringent seeds or germinating seeds.

**Supplementary Note 2. Aluminum stress experiment on *C. lanceolata*.**

**2.1 Astringent seed water extract preparation**

Mature seeds of *C. lanceolata* were prepared for the water extract. An incision was gently made in the middle of the seed coat with an anatomical needle. The color of the embryo was observed to confirm whether it was an astringent seed. After removing the seed coat and cleaning, the collected astringent seeds were freeze-dried to a constant weight in a vacuum freeze-dryer. They were then ground to powder for further experiments.

A mass of 100 g of astringent seed powder was weighed and dissolved in 10 L ultrapure water. Magnetic stirrers were used constantly during the dissolving process to help the powder fully dissolve. After 48 h, the solution was slowly filtered through filter paper. The wet powder on the paper was collected and re-filtered several times to minimize the solution loss.

**2.2 Aluminum stress experiment design**

The final water extract with a concentration of approximately 10 g L^-1^ was recorded as W0. The 10× diluted extraction was recorded as W10. Ultrapure water without adding this extraction was recorded as WN. They were used as high level, low level, and blank control treatments in subsequent experiments.

AlCl_3_·6H_2_O was used as the source of aluminum ions in the stress-response test. Three concentration gradients of 0.0 mmol·L^-1^ (Al0), 0.5 mmol·L^-1^ (Al1), and 1.0 mmol·L^-1^ (Al2) were set to stress the test seedlings. Combined with different concentrations of astringent seed water extract, the experimental treatments are shown in **Supplementary Table 28**.

**2.3 Seedling stress and culture**

To ensure that the genetic background of *C. lanceolata* seedlings used for the experiment was as consistent as possible, the seeds collected from the same mother individual in the third-generation seed orchard of Youxi National Forest Farm were selected as seedling materials. The seeds were germinated in a plant growth chamber and were cultivated for another 14 d. Seedlings with similar root length, seedling height, and leaf growth potential were selected for the stress test.

The stress treatment was performed using the hydroponics method. The astringent seed water extract at different concentrations was treated as the solvent. Different masses of AlCl_3_·6H_2_O were added to the solvent as shown in Table 1 to fit the aluminum ion concentration requirement. The nutrient elements were then added according to the Hoagland nutrient formula. Next, 1.0 mmol·L^-1^ NaOH/HCl solution was used to adjust the pH to 4.5. Three duplications were set for each treatment; thus, there were a total of 27 hydroponics tanks. Thirty seedlings were cultured in each tank. The tanks were placed in a plant growth chamber, and the parameters were set as follows: day-night cycle and temperature: light for 14 h at 25 °C, darkness for 10 h at 22 °C, light intensity of 110 μmol·m^-2^·s^-1^, and relative humidity of 75%. The stress test lasted for 30 days.

**2.4 Measurement of evaluation parameters**

After the stress test, the corresponding parameters of the roots and leaves of the seedlings were measured to evaluate the level of aluminum stress and the interaction between the water extract and aluminum stress. In the roots, the tip aluminum ion concentration, relative root elongation, tip malondialdehyde (MDA) content, and superoxide dismutase (SOD) activity were measured to evaluate the stress effect. The concentration of aluminum ions in the root tip was determined using the hematoxylin and eosin staining method. The MDA content was determined using the thiobarbituric acid colorimetric method. SOD activity was determined by nitrogen blue tetrazolium photochemical reduction determination. The relative root elongation was calculated by comparing root photographs before and after stress. To evaluate the effect of stress on the assimilative capacity of the seedlings, the photosynthetic parameters of leaves, including net photosynthetic rate (*Pn*), stomatal conductance (*Cond*), intercellular CO_2_ concentration (*Ci*), and transpiration rate (*Tr*), were measured using an LI-6400/XT portable photosynthetic apparatus (LI-COR, USA).

**Supplementary Note 3. Metabolome detection at different stages of astringent seeds and germinating seeds.**

**3.1 Sample collection, processing, and storage**

The samples used for the transcriptomic analysis were also used for the metabolomics assayed. The sampling collection, processing, and storage are described in the ‘Transcriptomic data and analysis’ in the ‘Online methods’ section. Please refer to this section for details of the sampling methods.

**3.2 Sample preparation and extraction**

All the freeze-dried samples were crushed using a mixer mill (MM 400, Retsch) with zirconia beads for 1.5 min at 30 Hz. One hundred milligrams of powder of each sample was weighed and extracted overnight at 4°C with 1.0 mL 70% aqueous methanol. Following centrifugation at 10000 × *g* for 10 min, the extracts were absorbed (CNWBOND Carbon-GCB SPE Cartridge, 250 mg, 3 mL; ANPEL, Shanghai, China, www.anpel.com.cn/cnw) and filtered (SCAA-104, 0.22 μm pore size; ANPEL, Shanghai, China, http://www.anpel.com.cn/) before LC-MS analysis.

**3.3 HPLC and ESI-Q TRAP-MS/MS conditions**

The sample extracts were analyzed using an LC-ESI-MS/MS system (HPLC, Shim-pack UFLC SHIMADZU CBM30A system, www.shimadzu.com.cn/; MS, Applied Biosystems 4500 Q TRAP, www.appliedbiosystems.com.cn/). The analytical conditions were as follows, HPLC: column, Waters ACQUITY UPLC HSS T3 C18 (1.8 µm, 2.1 mm × 100 mm); solvent system, water (0.04% acetic acid): acetonitrile (0.04% acetic acid); gradient program, 95:5 V/V at 0 min, 5:95 V/V at 11.0 min, 5:95 V/V at 12.0 min, 95:5 V/V at 12.1 min, 95:5 V/V at 15.0 min; flow rate, 0.40 mL/min; temperature, 40 °C; injection volume: 5 μL. The effluent was alternatively connected to an ESI-triple quadrupole-linear ion trap (QTRAP)-MS.

LIT and triple quadrupole (QQQ) scans were acquired on a triple quadrupole-linear ion trap mass spectrometer (Q TRAP), API 4500 Q TRAP LC/MS/MS System, equipped with an ESI Turbo Ion-Spray interface, operating in positive ion mode and controlled by the Analyst 1.6 software (AB Sciex). The ESI source operation parameters were as follows: ion source, turbo spray; source temperature, 550°C; ion-spray voltage (IS), 5500 V; ion source gas I (GSI), gas II(GSII), and curtain gas (CUR) were set at 55, 60, and 25.0 psi, respectively; the collision gas (CAD) was high. Instrument tuning and mass calibration were performed with 10 and 100 μmol/L polypropylene glycol solutions in QQQ and LIT modes, respectively. QQQ scans were acquired as multiple reaction monitoring (MRM) experiments with a collision gas (nitrogen) set to 5 psi. DP and CE for individual MRM transitions were performed with further DP and CE optimization.

**3.4 Qualitative and quantitative analyses of metabolites**

Based on the self-built database and the public databases (HMDB, http://www.hmdb.ca/; METLIN, https://metlin.scripps.edu/; and KEGG, http://www.kegg.jp/kegg/compound/), the qualitative analysis was carried out according to the secondary spectral information. The isotope signal, repeated signal containing K^+^, Na^+^, and NH4^+^ ions, and repeated signal of fragment ions with other larger molecular weight substances were removed.

Quantification was accomplished using the MRM model of tripe four-stage rod mass spectrometry. After obtaining the metabolite spectrum analysis data of different samples, the peak integral area was performed for all the mass spectrum peaks, and integral correction was performed for the mess spectrum peaks of the same metabolite in different samples.

**3.5 Identification and verification of different expressive metabolites**

Analyst v1.6.1 software was used to process the raw mass spectrometry data. MultiaQuant software was used to integrate and correct the chromatographic peaks. The peak area of each chromatographic peak represented the relative content of the corresponding compounds. Finally, all the chromatographic peak integral area data were derived. Principal component analysis, orthogonal partial least squares discriminant analysis (OPLS-DA), and Pearson correlation coefficient calculations were performed using the RStudio software (www.r-project.org/).

Metabolites of different groups were preliminarily screened by the variable importance in project (VIP) value obtained from the OPLS-DA model. The fold change value was combined to further screen out the different expressive metabolites (DEMs). The metabolites with FC of ≥ 2.0 or FC of ≤ 0.5 were considered to differ more than twice or less than 0.5 between the AS and GS groups, while a VIP value of ≥ 1.0 was indicative of a significant effect on the classification of samples in each group in the OPLS-DA model. Metabolites that fitted both of the conditions were selected as DEMs.

To verify the accuracy of the relative quantificational results generated from the LC-MS signals, 16 metabolites were selected randomly for further HPLC quantification. The HPLC conditions were the same as those described in Section 2.3. Standards of the selected metabolites were obtained from the Shinemro chemical platform (http://www.shinemro.com).

**Supplementary Note 4. Determination of aluminum in Chinese fir leaves.**

We analyzed the aluminum content in Chinese fir leaves at different stages of development. All materials from ten individuals randomly selected in third-generation seed orchard of Youxi National Forest Farm, Fujian Province, China. A total of five groups of materials were collected with 10 replicates in each group. The five materials are as follows: the young (YL) and old leaves from living branches, survive for one year (WL1) and five years (WL2) of leaves in withered branches, and fallen two months of leaves (WL3) with withered branches.

After 1 h deactivation of enzymes at 105 °C, all the samples were dried at 70 °C until a constant weight was obtained. The dry samples were ground and crushed, and then were filtered through a 0.5 mm sieve. Sample in each group were accurately weighted at 0.2 g respectively, and were digested with nitric acid and hydrogen peroxide solution using microwave digestion system (Multiwave ECO). The digested solution was diluted with pure water to a final volume of 50 mL after it cooled to room temperature. After that, the determination of Al ions concentrations in each group samples were presented using inductively coupled plasma method by a PE Optima 8000 system (PerkinElmer, USA).

### Supplementary Figures

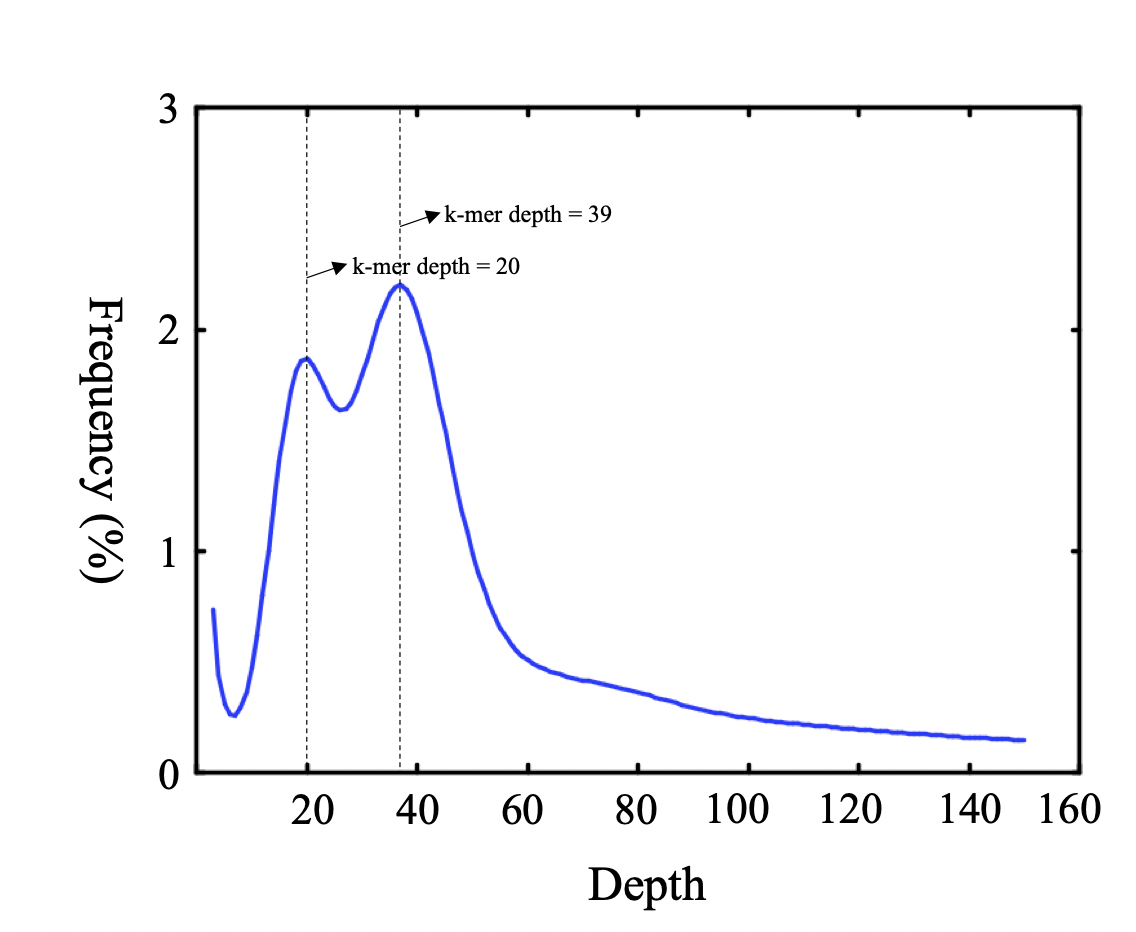

#### Supplementary Figure 1. Genome size and heterozygosity of *C. lanceolata* estimation using *K*-mer distribution.

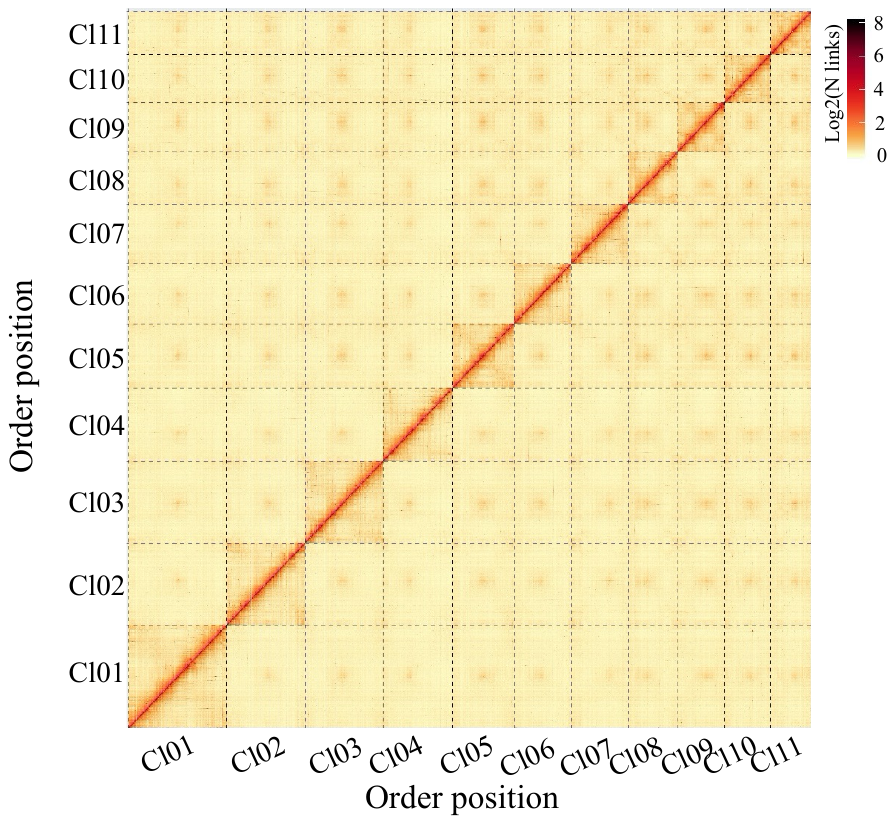

#### Supplementary Figure 2. Hi-C interaction heatmap for *C. lanceolata* genome showing interactions among eleven chromosomes*.* Darker red pixels denote higher contact probabilities. Most interactions were observed within the chromosomes.

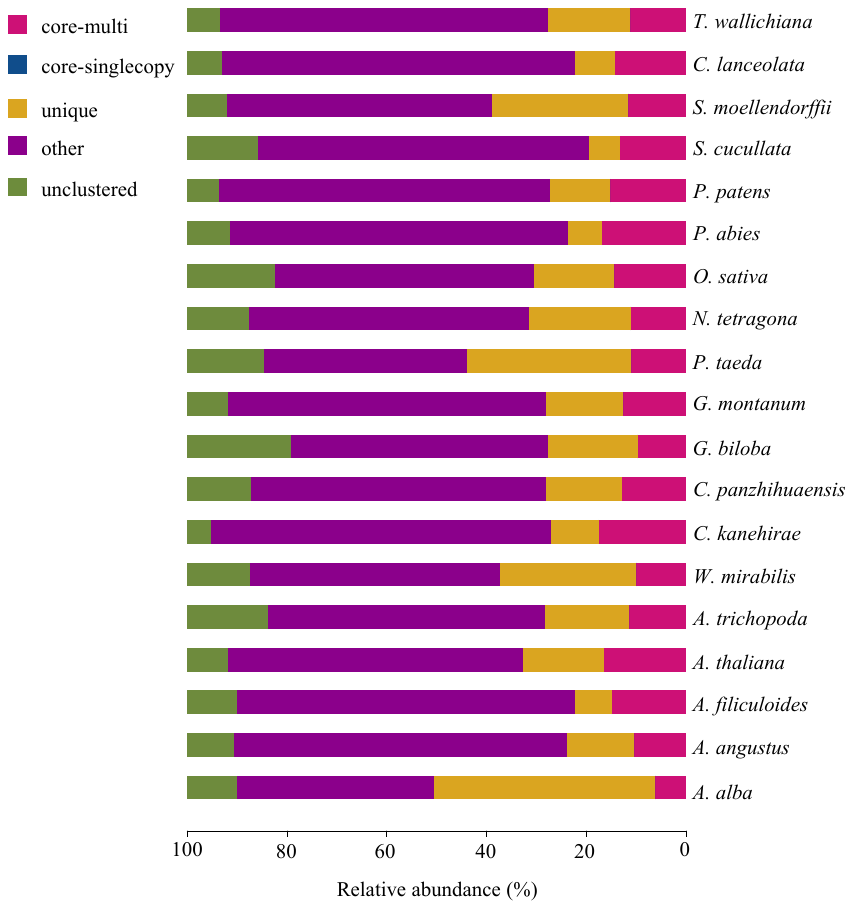

#### Supplementary Figure 3. Orthologous genes in *C. lanceolata* and other species.

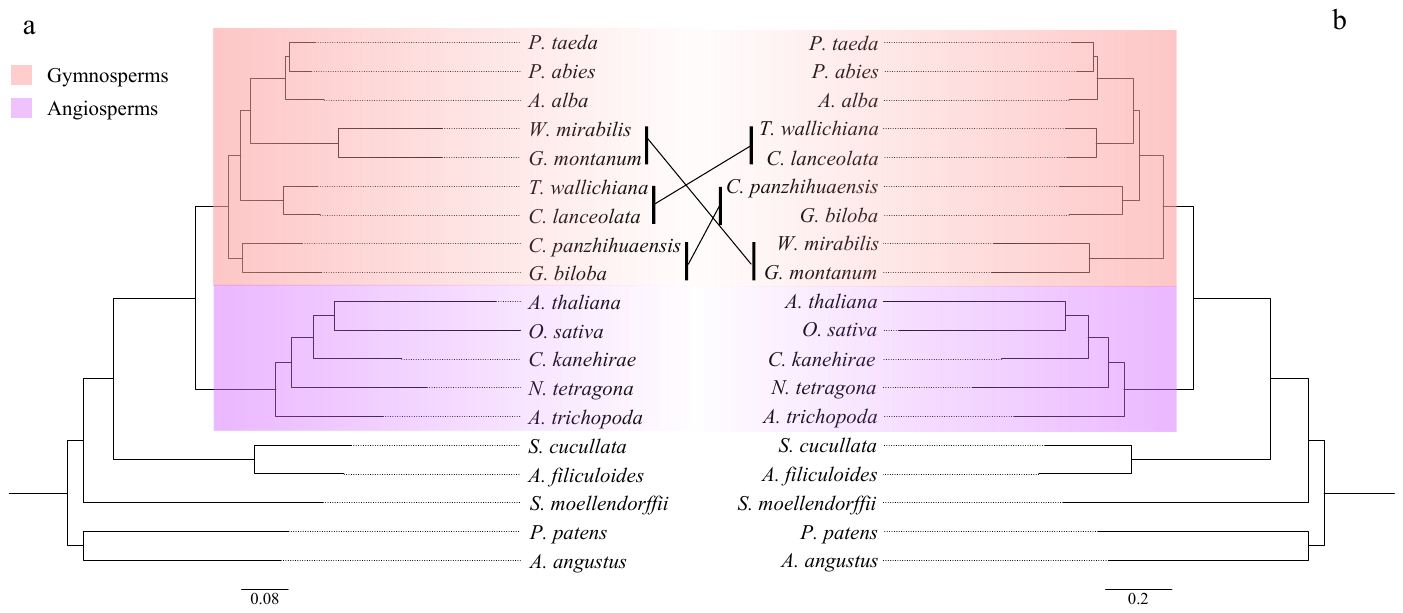

#### Supplementary Figure 4. Phylogenetic tree constructed by different methods based on single copy-genes. a. The concatenated tree based on amino acid, concatenated tree constructed by the first and two codons, and the Bayesian tree. b. The concatenated tree based on nucleotides, ASTRAL tree, and ASTRAL tree based on constructed by the first and two codons. The topological structure of concatenated tree based on amino acid, concatenated tree based on constructed by the first and two codons, and the Bayesian tree are the same, while the topological structure of concatenated tree based on nucleotides, ASTRAL tree, and ASTRAL tree based on constructed by the first and two codons are same.

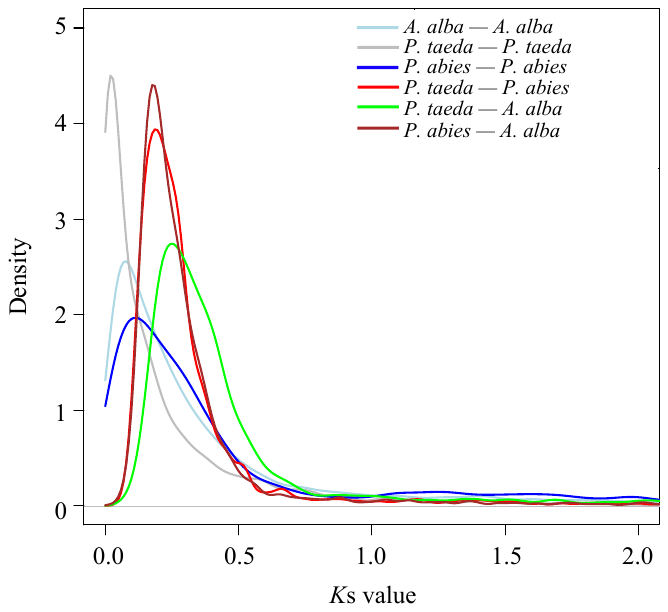

#### Supplementary Figure 5. *K*s distribution in *C. lanceolata* and Pinaceae species genomes.

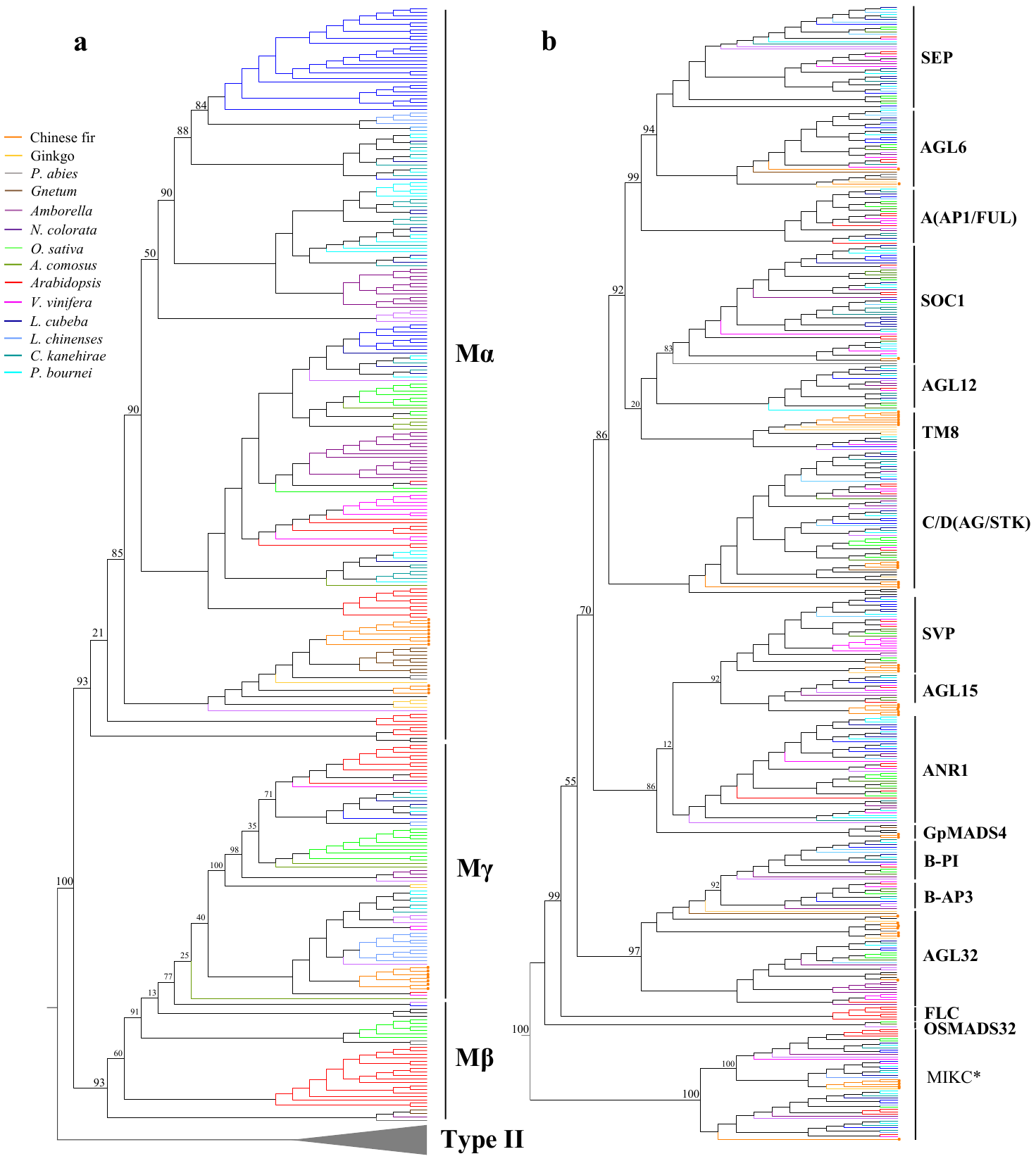

#### Supplementary Figure 6. Phylogenetic tree of MADS-box genes from gymnosperms and angiosperms. a. Phylogenetic tree of MADS-box Type I genes. b. Phylogenetic tree of MADS-box Type II genes.

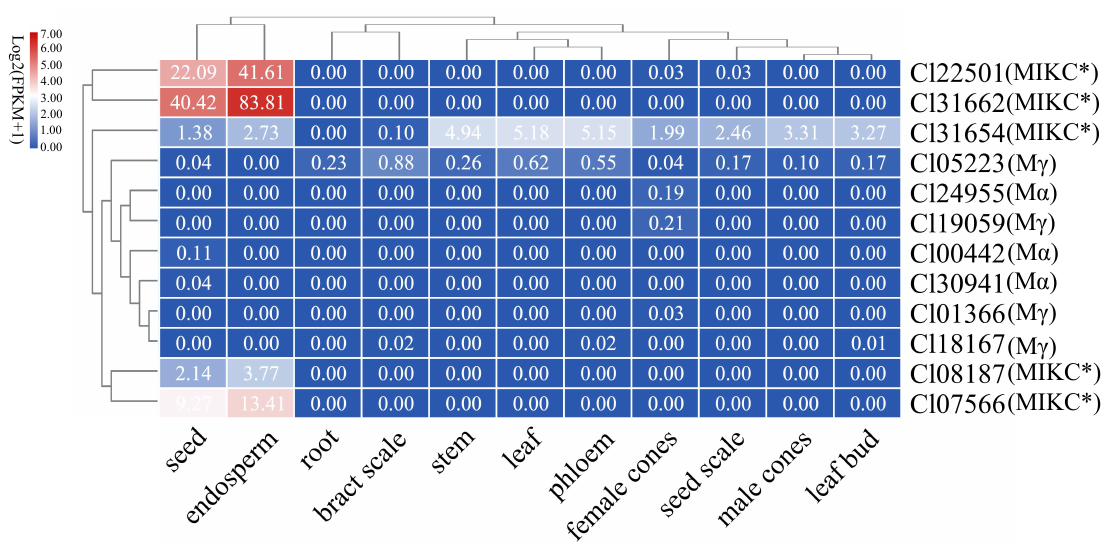

#### Supplementary Figure 7. The expression of MADS type I genes in different tissues of the *C. lanceolata*.

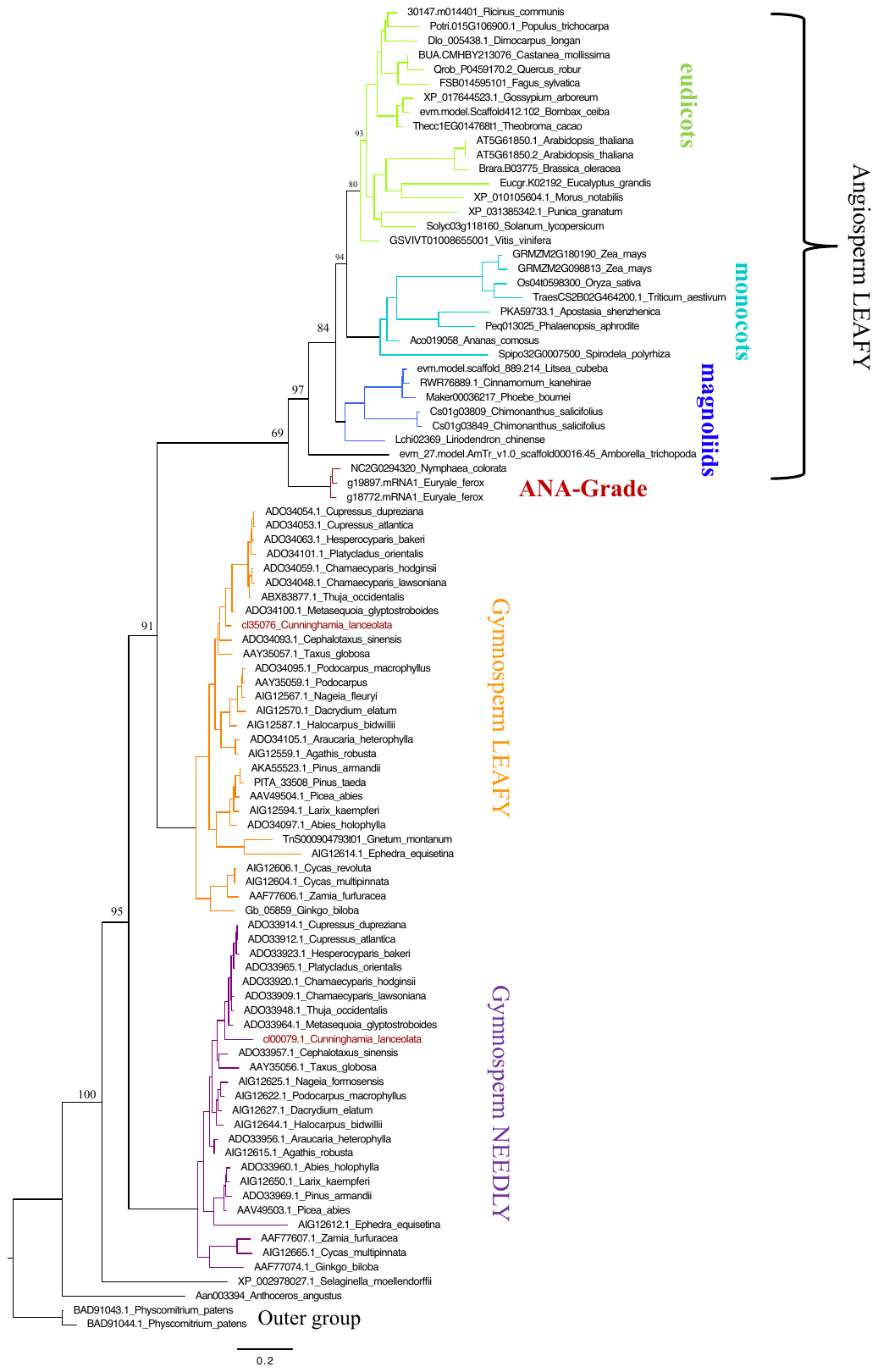

#### Supplementary Figure 8. Phylogenetic analysis of LFY/FLO and NLY orthologs genes from gymnosperms, angiosperms, *Selaginella*, *Anthoceros* and *Physcomitrium*. The underscore of the gene ID is followed by the Latin name of the species.

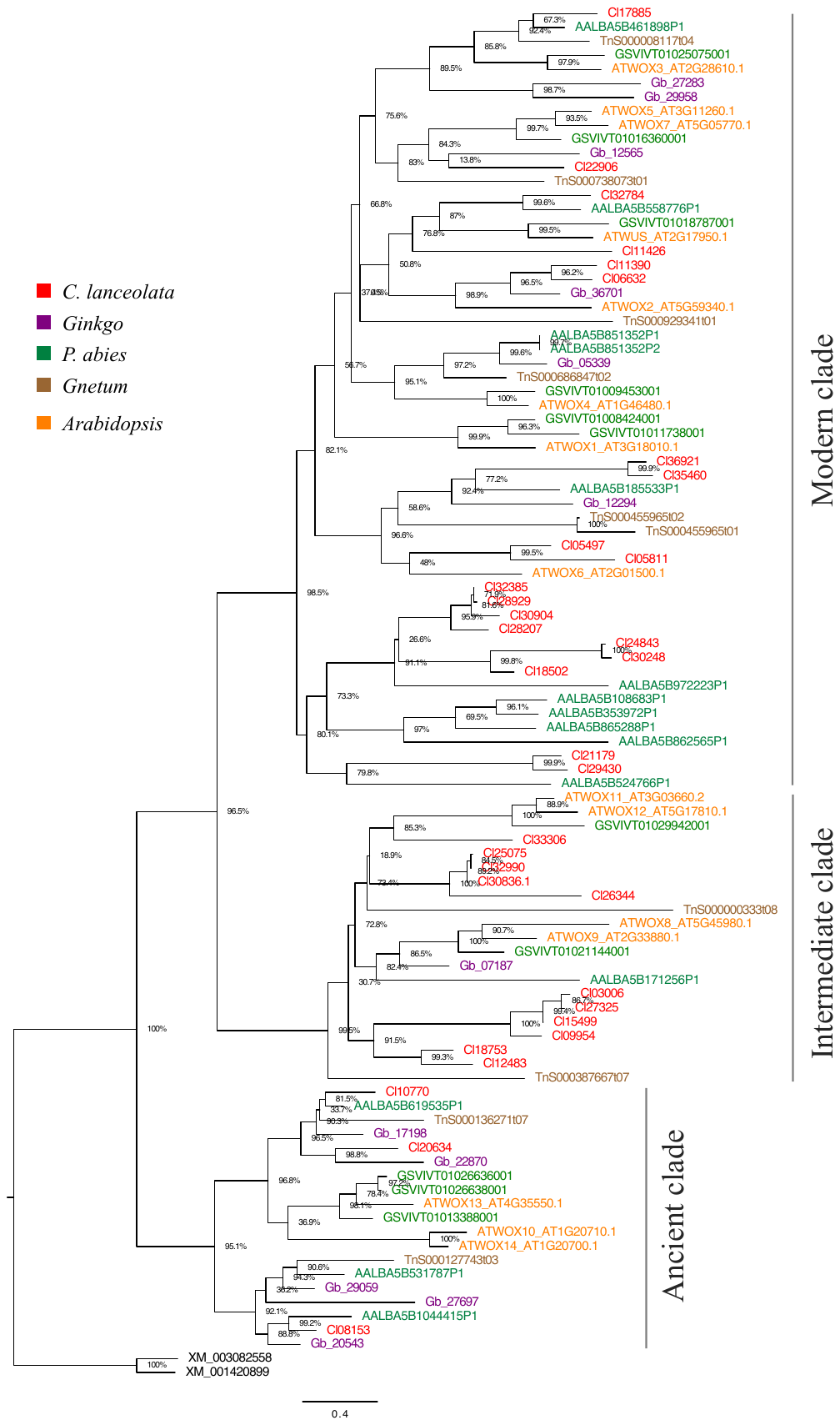

#### Supplementary Figure 9. Phylogenetic relationships of WOX gene families from *C. lanceolata*, Ginkgo, *P. abies*, *Gnetum*, *Arabidopsis*. Two genes, *XM_001420899* (*Ostreococcus lucimarinus*) and *XM_003082558* (*O. tauri*), from algae act as outer group. A total of 36 *WOX* genes were identified in *C. lanceolata,* much higher than those found in Ginkgo and *P. abies*. All land plant *WOX* genes have been classified into the ancient, intermediate, and modern clades.

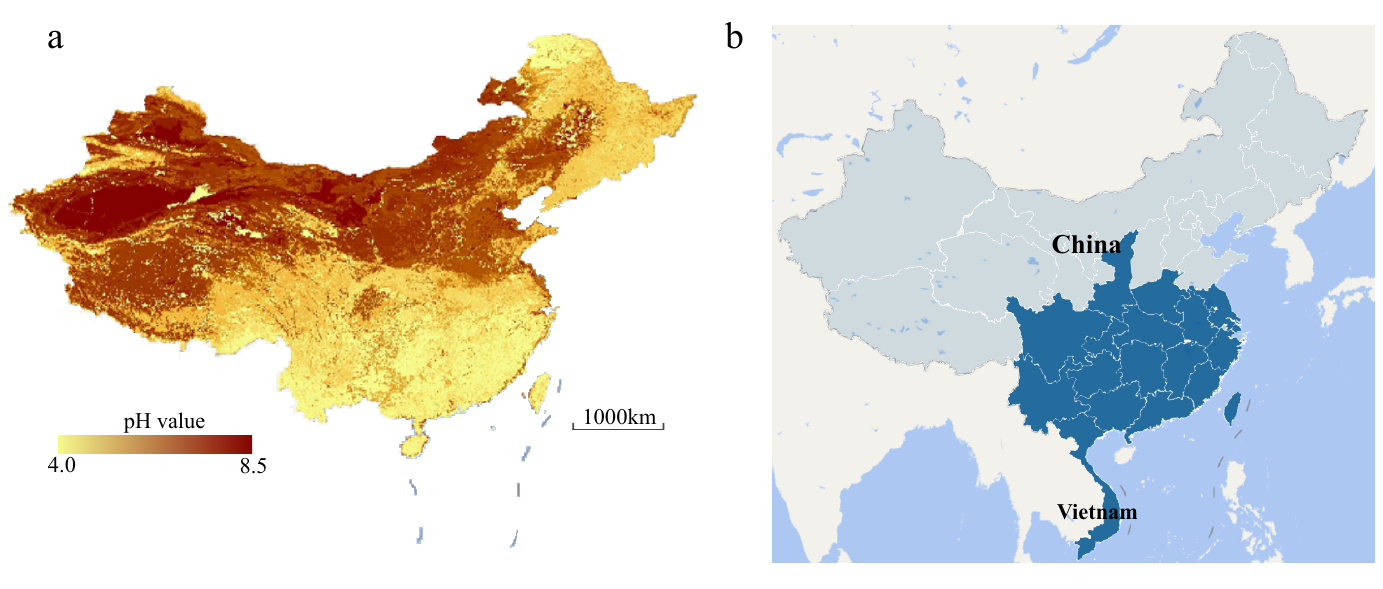

#### Supplementary Figure 10. The distribution of *C. lanceolata* coincides with the distribution of acid red soil. a. China’s acid soil distribution map. Data sourced from National Soil Information Service Platform of China (<http://www.soilinfo.cn>) b. *C. lanceolata* distribution map.

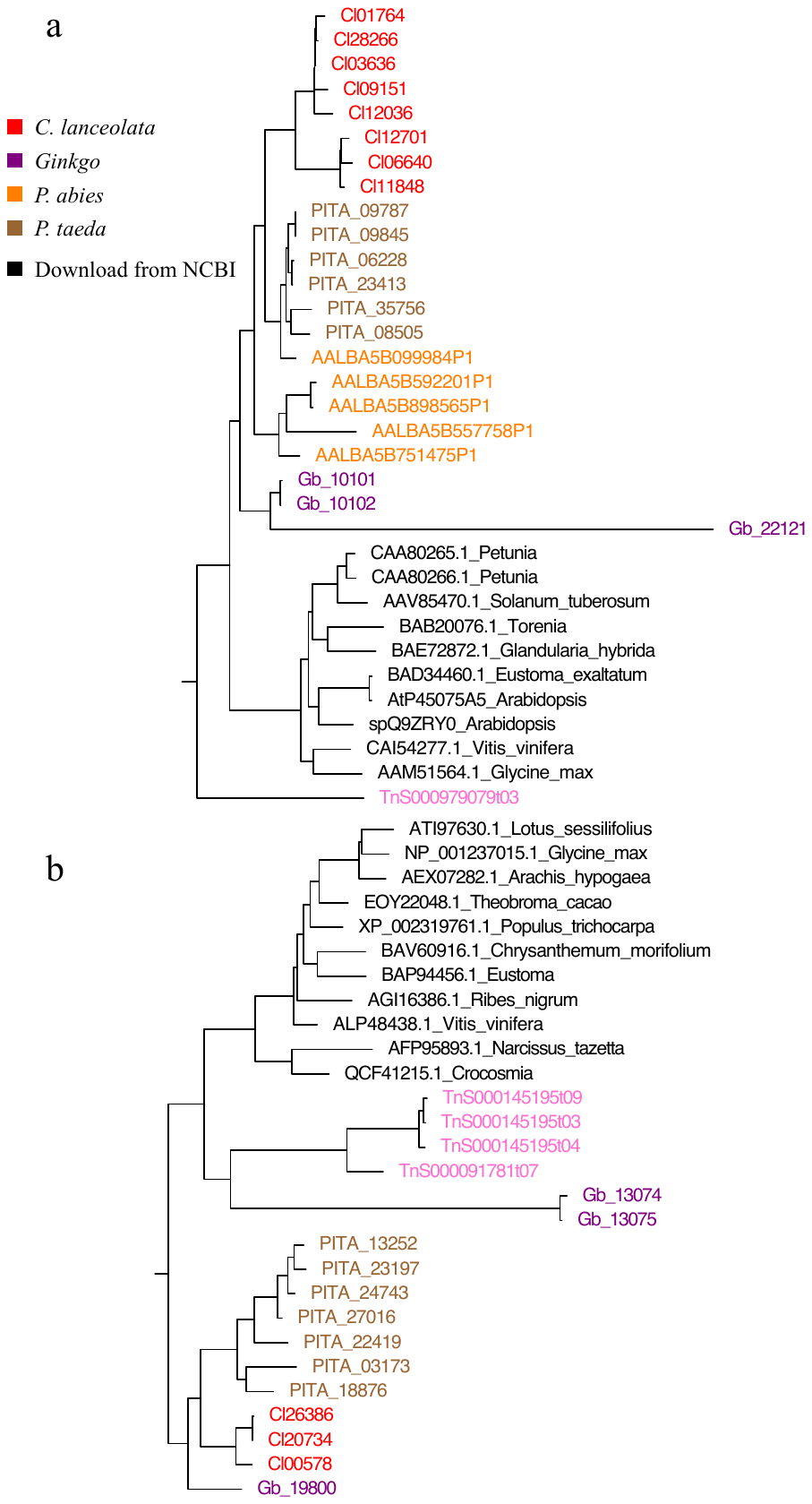

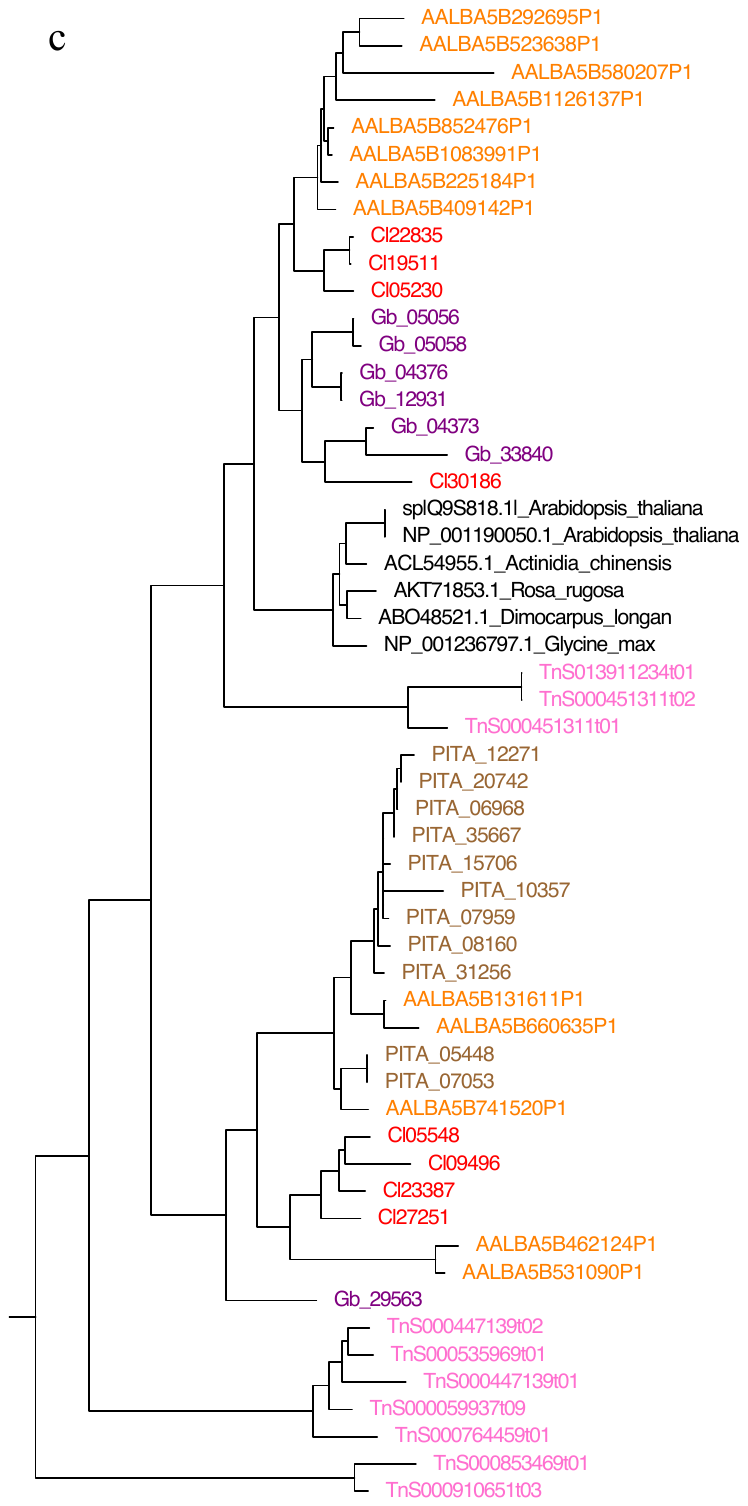

#### Supplementary Figure 11. Phylogenetic tree of flavonoid 3',5'-hydroxylase (F3'5'H), flavonoid 3'-hydroxylase (F3'H), and flavanone-3-hydroxylase (F3H) from seed plants. a. Flavonoid 3',5'-hydroxylase (F3'5'H). b. Flavonoid 3'-hydroxylase (F3'H). c. Flavanone-3-hydroxylase (F3H).

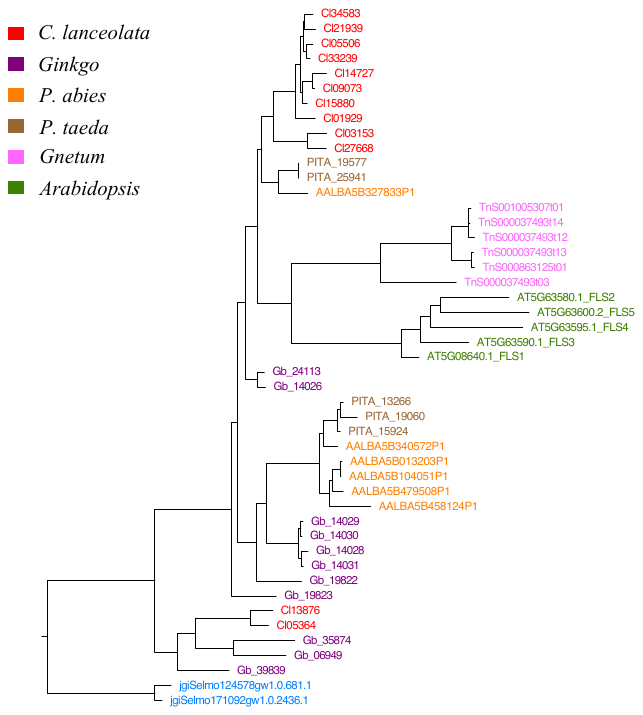

#### Supplementary Figure 12. Phylogenetic relationships of flavonol synthase (FLS) gene. Two genes, *jgiSelmo124578gw1.0.681* and *jgiSelmo171092w1.0.2436.1*, from *Selaginella tamariscina* as outer group.

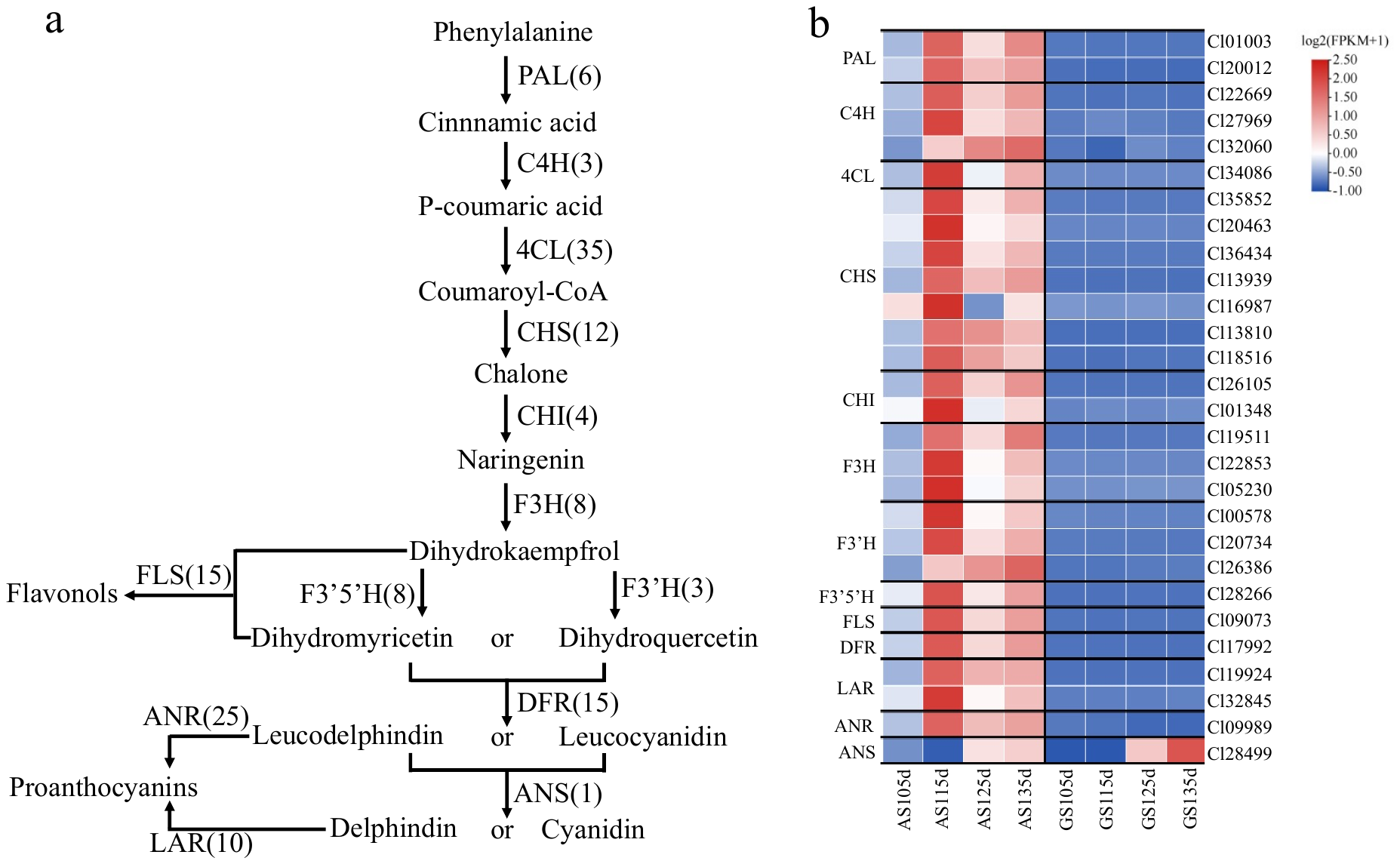

#### Supplementary Figure 13. Expression of flavonoid synthesis related genes in astringent seeds (AS) and germinating seeds (GS) during different developmental stage. a. Synthesis pathways of flavonoids. b. The expression patterns of genes related to flavonoid synthesis in astringent seeds and germinated seeds at different stages. (see the full name of genes in Supplementary Table 23)

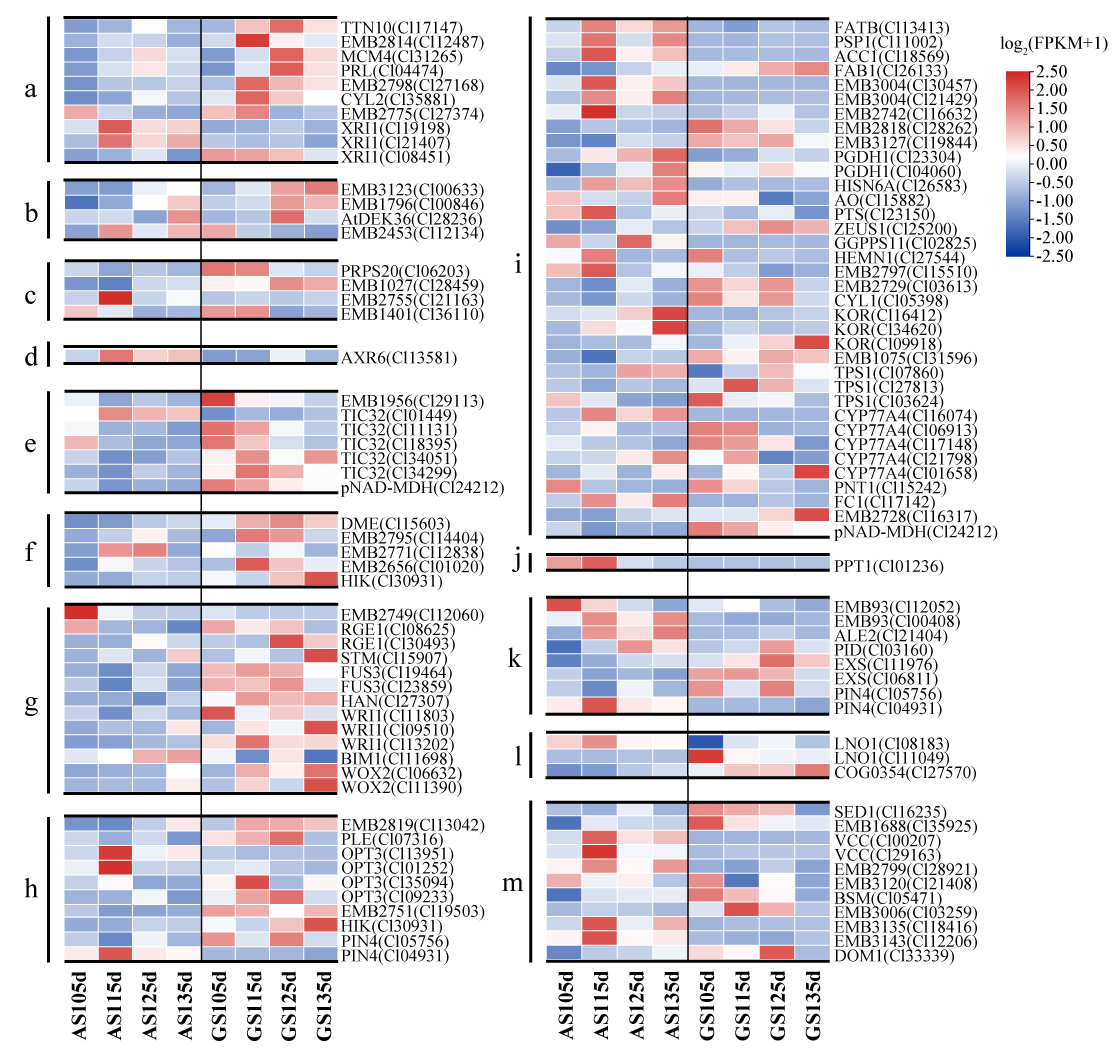

#### Supplementary Figure 14. Expression pattern of 109 differential embryo-defective genes (EMBs) in astringent seeds (AS) and germinating seeds (GS) during four developmental stages. The letters on the left of the heat map represent the functional classification of the gene, the right side is the gene name, and the *C. lanceolata* gene ID in parentheses. a. DNA synthesis/repair. b. DNA synthesis/modification. c. Protein synthesis. d. Protein degradation. e. Protein modification/transport. f. Chromosome dynamics. g. Transcriptional regulation. h. Cell structure. i. metabolism. j. Energy electron. k. Signaling and regulatory pathways. l. other (miscellaneous). M. Uncertain/unknown.

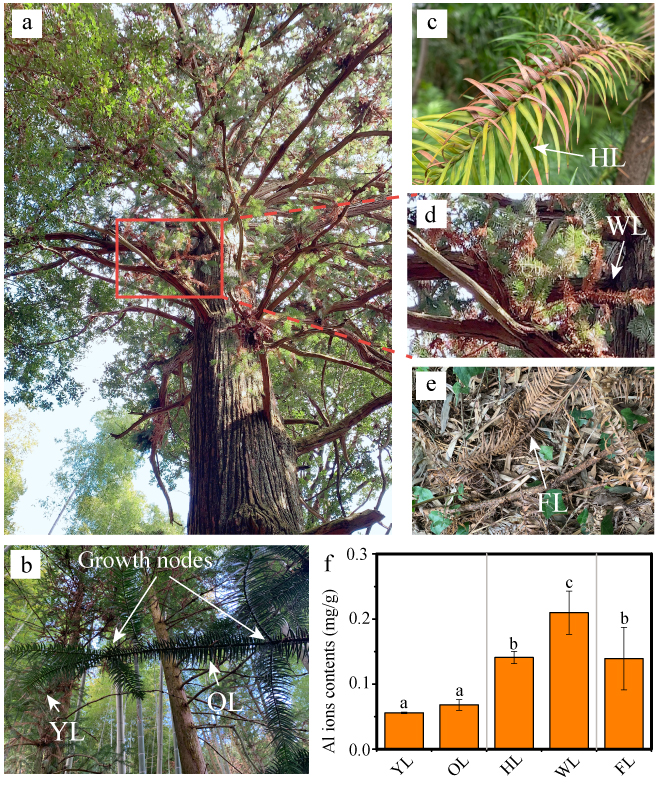

#### Supplementary Figure 15. *C. lanceolata* leaves in different growth periods. a. A mature individual of *C. lanceolata*. Living and withered leaves could be observed in this individual. b-e. The details of the leaves at different stages. b. The young leaves (YL) and the old leaves (OL). c. The half-withered leaves (HL). d. Persistent withered leaves (WL). e. The fallen withered leaves (FL). Withered leaves fall off with the branches only when self-pruning occurs. f. Aluminum ion content in different growth stages of leaves. There is no significant difference at p = 0.05 level between groups labeled with same letter.

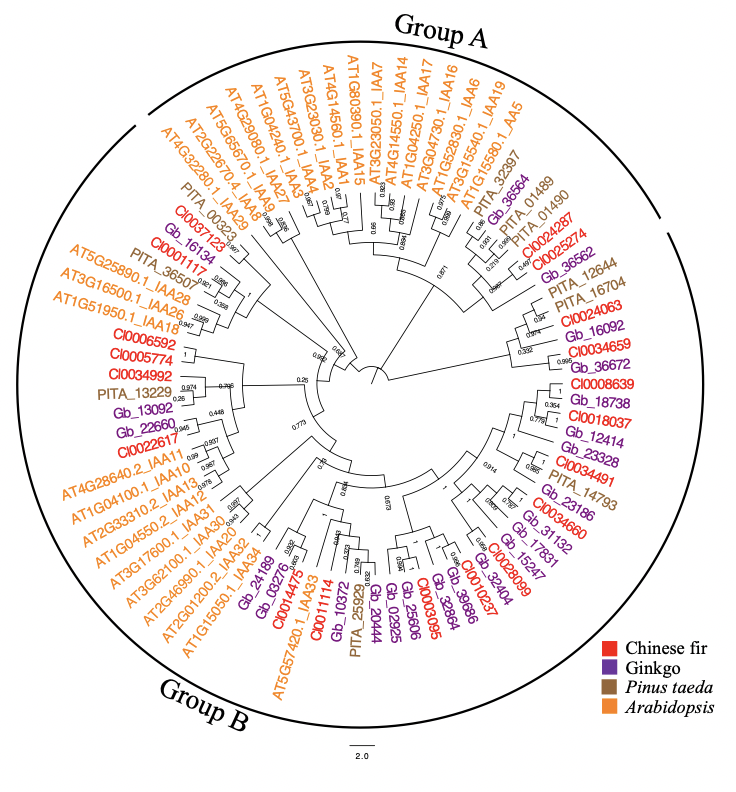

#### Supplementary Figure 16. Phylogenetic relationships of AUX/IAA gene families from *C. lanceolata*, Ginkgo, *P. taeda*, and *Arabidopsis*. Auxin regulates cell division and elongation to drive plant growth and development ^1^. AUX/IAA is the main gene family that perception auxin in plants ^2^. Low R: FR of shade indirectly promotes auxin synthesis genes ^3^. We have identified 19, 21 and nine Aux/IAA genes in *C. lanceolata*, Ginkgo and *P. taeda* genomes, respectively. Phylogenetic tree indicated that Aux/IAA proteins can be classified into two major groups, A and B similar to Arabidopsis ^4^. The Aux/IAA gene of *C. lanceolata*, Ginkgo and *P. taeda* were mainly amplified in Group B.

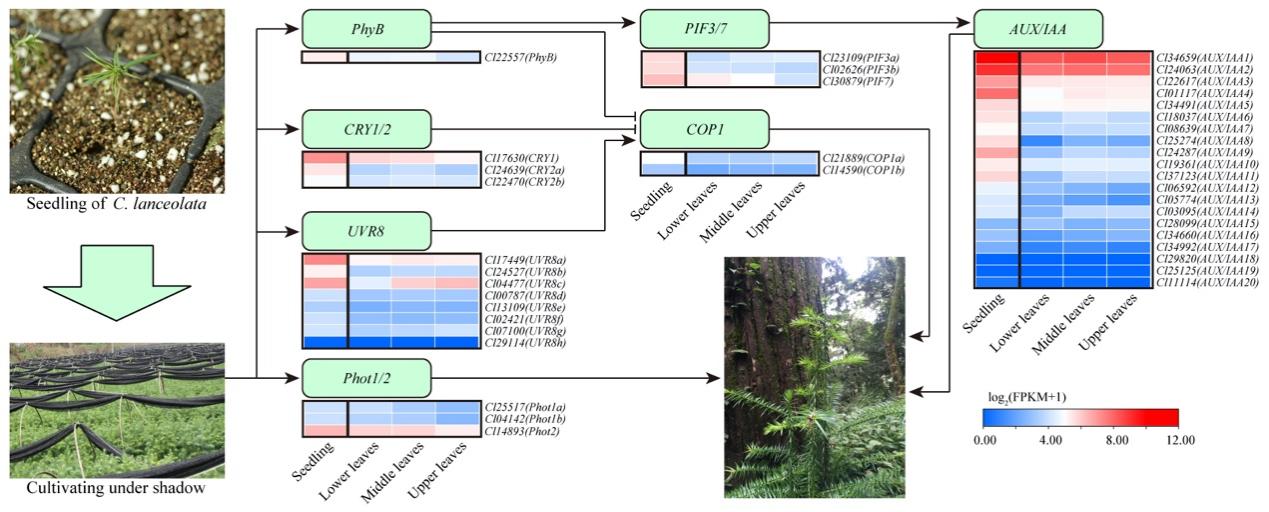

#### Supplementary Figure 17. Analysis of shade tolerance of *C. lanceolata* seedlings. a. Expression profile of shade-tolerance-resistance-related genes in the leaves of seedlings and adult trees. Seeding refers to the leaves of seedlings cultivated for two years. The lower, middle, and upper leaves came from an adult *C. lanceolata* with a height of 30 m. (See the full name of genes in Supplementary Table 25)

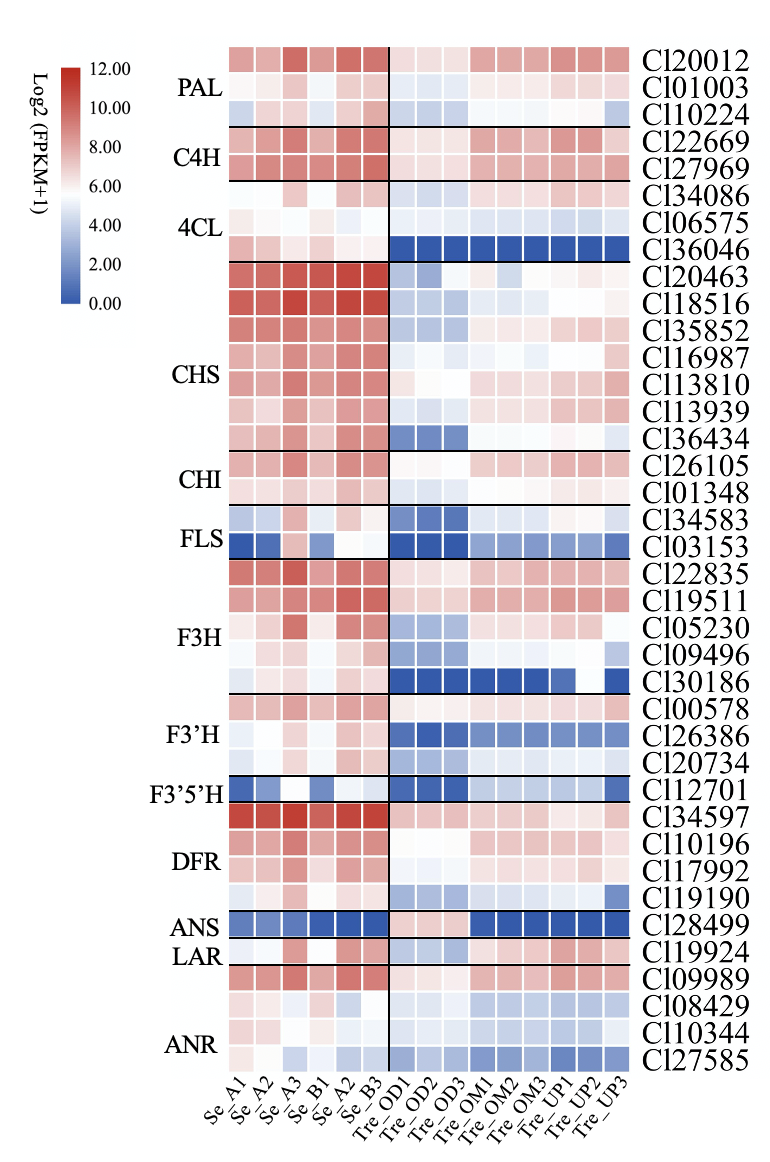

#### Supplementary Figure 18. Expression of flavonoid synthesis related genes in *C. lanceolata* leaves of seedlings and adult tree. Se_A and Se_B are the leaves of 228 and 226 clonal seedlings cultivated for two years, respectively. Tre_OD, Tre_OM, and Tre_OP are the lower, middle, and upper leaves of the *C. lanceolata* with a height of 30m. (See the full name of genes in Supplementary Table 23)

### Supplementary Tables

**Supplementary Table 1.** **The statistics of the Illumina sequencing data volume, sequencing depth and quality inspection of each library of *C. lanceolata* genome.**

| **Library** | **Data (Gb)** | **Depth (X)** | **Q20 (%)** | **Q30 (%)** |
| --- | --- | --- | --- | --- |
| 270_1 bp | 67.40 | 6.47 | 94.87 | 85.19 |
| 270_2 bp | 61.53 | 5.91 | 94.23 | 86.42 |
| 270_3 bp | 65.17 | 6.25 | 94.67 | 87.25 |
| 270_4 bp | 58.48 | 5.61 | 94.94 | 87.86 |
| 270_5 bp | 63.25 | 6.07 | 94.34 | 86.70 |
| 270_6 bp | 66.94 | 6.42 | 94.42 | 86.88 |
| 270_7 bp | 70.19 | 6.74 | 94.50 | 87.06 |
| 270_8 bp | 63.46 | 6.09 | 94.31 | 86.47 |
| Total | 516.42 | 49.56 | -- | -- |

Library: the sequencing library of the survey graph; Data (Gb): the amount of sequencing data of the corresponding sequencing library; Depth (X): the depth of sequencing; Q20 (%): the percentage of bases with a sequencing quality value of 20 or more; Q30 (%): The proportion of bases whose sequencing quality value is above 30.

**Supplementary Table 2. The statistics of the quality on the PacBio sequencing data of *C. lanceolata* genome.**

| **Reads type** | **Reads num** | **Total bases (bp)** | **Reads N50 (bp)** | **Mean length (bp)** | **Longest read (bp)** |
| --- | --- | --- | --- | --- | --- |
| Subreads | 89,507,038 | 1,113,159,857,423 | 20,648 | 12,437 | 129,985 |
| ZMWreads | 66,589,949 | 959,970,038,241 | 23,042 | 14,416 | 129,985 |

Reads Type: Subreads is the data set produced by Subreads, and ZMWreads is the longest subreads data set in the ZMW hole.

#### Supplementary Table 3. Assembly statistics of the *C. lanceolata* genome.

| **Type** | **Number/length/percentage** |
| --- | --- |
| Contig number | 28,364 |
| Contig length (bp) | 11,242,038,337 |
| Contig N50 (bp) | 2,155,103 |
| Contig N90 (bp) | 232,277 |
| Contig max (bp) | 22,388,200 |
| GC content (%) | 36.95 |

#### Supplementary Table 4. Comparison of the BUSCO assessment of the assembled and annotated gymnosperms genomes.

| **Species** | **Assembled genome BUSCO** | **Annotated genome BUSCO** |
| --- | --- | --- |
| *G. biloba*^5^ | 44.80% | 60.04% |
| *P. abies*^7^ | 35.25% | 25.40% |
| *P. taeda*^9^ | 22.30% | 23.23% |
| *G. montanum^6^* | 74.85% | 82.84% |
| *C. lanceolata* | 50.00% | 85.07% |
| *Taxus chinensis*^10^ | / | 65.18% |

#### Supplementary Table 5. Illumina sequence alignment statistics of the genome of *C. lanceolata*.

| **Type** | **Number/percentage** |
| --- | --- |
| Total reads | 1,882,235,817 |
| Mapped reads | 1,864,713,766 |
| Mapped (%) | 99.07 |
| Properly mapped reads | 1,737,960,488 |
| Properly mapped (%) | 92.83 |

Total reads: reads statistics of filtered data; Mapped Reads: Reads statistics of matched genomes; Properly mapped reads: Reads statistics of mapped genomes and paired.

#### Supplementary Table 6. The length of chromosome by Hi-C assemble of the *C. lanceolata* genome.

| **Chromosome** | **Cluster Num** | **Cluster Len (bp)** | **Order Num** | **Order Len (bp)** |
| --- | --- | --- | --- | --- |
| Cl01 | 3,692 | 1,550,680,665 | 3,251 | 1,488,319,881 |
| Cl02 | 2,701 | 1,263,832,510 | 2,267 | 1,184,309,377 |
| Cl03 | 2,640 | 1,234,995,222 | 2,238 | 1,182,746,149 |
| Cl04 | 2,760 | 1,178,736,379 | 2,063 | 1,047,693,580 |
| Cl05 | 1,913 | 960,425,685 | 1,611 | 927,725,105 |
| Cl06 | 1,901 | 931,596,408 | 1,552 | 862,932,688 |
| Cl07 | 1,804 | 897,221,591 | 1,527 | 863,305,216 |
| Cl08 | 1,365 | 778,813,723 | 1,139 | 747,012,144 |
| Cl09 | 1,232 | 749,408,730 | 988 | 712,708,477 |
| Cl10 | 2,070 | 710,732,703 | 1,914 | 694,515,374 |
| Cl11 | 1,682 | 637,087,857 | 1,491 | 616,297,228 |
| Total (Ratio %) | 23,760(83.03) | 10,893,531,473 (96.9) | 20,041 (84.35) | 10,327,565,219 (94.8) |

Cluster Len (bp): The length of the sequence located on the chromosome. Order Len (bp): In the sequence located on the chromosome, the length of the sequence can be determined in order and direction.

#### Supplementary Table 7. The statistic result of Hi-C assembled of the genome of *C. lanceolata*.

| **Type** | **Contig Scaffold** | |
| --- | --- | --- |
|  | **Size (bp)** | **Size (bp)** |
| N90 | 227,972 | 616,446,228 |
| N50 | 2,097,655 | 927,886,105 |
| Longest | 21,974,839 | 1,488,644,881 |
| Total length | 11,242,038,337 | 11,244.041,337 |

#### Supplementary Table 8. The prediction of gene numbers of the *C. lanceolata* genome.

| **Method** | **Software** | **Species** | **Gene number** |
| --- | --- | --- | --- |
| Ab initio | Genscan |  | 68,124 |
|  | GlimmerHMM |  | 190,423 |
|  | GeneID |  | 179,448 |
|  | SNAP |  | 118,942 |
| Homology-based | GeMoMa | *Arabidopsis thaliana* | 25,405 |
|  |  | *Ginkgo biloba* | 38,598 |
|  |  | *Gnetum montanum* | 29,888 |
|  |  | *Picea abies* | 32,849 |
|  |  | *Populus trichocarpa* | 30,476 |
|  |  | *Pinus taeda* | 53,107 |
| RNAseq | TransDecoder |  | 115,300 |
|  | GeneMarkS-T |  | 58,760 |
|  | PASA |  | 55,025 |
| Integration | EVM |  | 37,225 |

#### Supplementary Table 9. The statistics results of function annotation in genome of *C. lanceolata*.

| **Annotation database** | **Annotated number** | **Percentage (%)** |
| --- | --- | --- |
| GO Annotation | 18,857 | 50.66 |
| KEGG Annotation | 12,874 | 34.58 |
| KOG Annotation | 20,771 | 55.8 |
| Pfam Annotation | 29,051 | 78.04 |
| Swissprot Annotation | 26,239 | 70.49 |
| TrEMBL Annotation | 34,202 | 91.88 |
| Nr Annotation | 34,392 | 92.39 |
| All Annotation | 34,559 | 92.84 |

#### Supplementary Table 10. Statistics on the annotation of non-coding RNA of the *C. lanceolata* genome.

| **RNA classification** | **Number** | **Family** |
| --- | --- | --- |
| miRNA | 50 | 13 |
| rRNA | 2,930 | 4 |
| tRNA | 3,955 | 24 |
| snRNA | 284 | 8 |
| snoRNA | 208 | 2 |

#### Supplementary Table 11. Statistics of *C. lanceolata* gene structure information.

| **Gene type** | **Number/length** |
| --- | --- |
| Gene number | 37,225 |
| Gene length (bp) | 1,112,864,908 |
| Average gene length (bp) | 29,895.63 |
| Exon number | 164,956 |
| Exon length (bp) | 43,643,130 |
| Average exon length (bp) | 264.57 |
| Intron number | 164,955 |
| Intron length (bp) | 1,069,221,778 |
| Average intron length (bp) | 6,481 |

Supplementary Table 12. Statistic of different types of repetitive sequences in gymnosperms that annotated genome.

| **Species** | ***C. lanceolata*** | ***Ginkgo***^5^ | ***G. montanum***^6^ | ***P. abies***^7^ |
| --- | --- | --- | --- | --- |
| protein coding genes | 37,225 | 41,840 | 27,491 | 58,587 |
| repetitive sequences | 92.31% | 76.58% | 85.9% | 70% |
| LTR | 69.92% | 60.65% | 25.42% | 58% |
| DNA | 8.26% | 3.34% | 0.83% | 1% |
| LINE | 3.26% | 4.34% | 16.69% | 1% |
| SINE | 0.01% | 0.00% | 0.53% | 0.00% |

#### Supplementary Table 13. Statistic of different types of repeat sequence in *C. lanceolata* genome.

| **Type** | **Number** | **Length** | **Rate (%)** |
| --- | --- | --- | --- |
| ClassI | 13,217,000 | 9,755,331,388 | 86.78 |
| ClassI/DIRS | 954,220 | 928,392,143 | 8.26 |
| ClassI/LINE | 595,272 | 365,970,956 | 3.26 |
| ClassI/LTR | 485,367 | 411,052,389 | 3.66 |
| ClassI/LTR/Copia | 3,107,217 | 2,684,662,505 | 23.88 |
| ClassI/LTR/Gypsy | 4,233,916 | 4,764,617,838 | 42.38 |
| ClassI/LTR\|DIRS | 140 | 65,857 | 0 |
| ClassI/PLE\|LARD | 3,797,243 | 2,243,877,805 | 19.96 |
| ClassI/SINE | 3,706 | 805,063 | 0.01 |
| ClassI/SINE\|TRIM | 832 | 875,820 | 0.01 |
| ClassI/TRIM | 33,556 | 21,340,727 | 0.19 |
| ClassI/Unknown | 5,531 | 4,488,270 | 0.04 |
| ClassII | 514,745 | 382,500,579 | 3.4 |
| ClassII/Crypton | 145 | 183,686 | 0 |
| ClassII/Helitron | 70,297 | 34,790,020 | 0.31 |
| ClassII/MITE | 4,567 | 2,757,596 | 0.02 |
| ClassII/Maverick | 19,158 | 11,853,833 | 0.11 |
| ClassII/TIR | 368,383 | 249,279,504 | 2.22 |
| ClassII/Unknown | 52,195 | 84,861,181 | 0.75 |
| PotentialHostGene | 183,062 | 118,181,144 | 1.05 |
| SSR | 26,634 | 30,066,332 | 0.27 |
| Unknown | 1,401,192 | 700,661,880 | 6.23 |
| Total | 13,941,441 | 10,377,888,279 | 92.31 |

#### Supplementary Table 14. Statistic result of clustered gene families of 19 species.

| **Species** | **Genes** | **Unclustered genes** | **Clustered genes** | **Familys** | **Unique families** | **Unique families genes** | **Common families** | **Common families genes** | **Average genes per family** |
| --- | --- | --- | --- | --- | --- | --- | --- | --- | --- |
| *A. alba* | 50,466 | 4,996 | 45,470 | 11,628 | 2,556 | 22,384 | 876 | 3,119 | 3.91 |
| *A. angustus* | 14,629 | 1,358 | 13,271 | 8,461 | 428 | 1,953 | 876 | 1,529 | 1.568 |
| *A. filiculoides* | 20,203 | 2,016 | 18,187 | 9,615 | 473 | 1,478 | 876 | 3,015 | 1.892 |
| *A. thaliana* | 27,416 | 2,244 | 25,172 | 10,084 | 983 | 4,422 | 876 | 4,523 | 2.496 |
| *A. trichopoda* | 26,846 | 4,326 | 22,520 | 10,929 | 986 | 4,546 | 876 | 3,075 | 2.061 |
| *W. mirabilis* | 39,019 | 4,877 | 34,142 | 10,458 | 1,519 | 10,647 | 876 | 3,921 | 3.265 |
| *C. kanehirae* | 26,531 | 1,273 | 25,258 | 10,012 | 571 | 2,551 | 876 | 4,660 | 2.523 |
| *C. panzhihuaensis* | 32,353 | 4,114 | 28,239 | 11,851 | 1,083 | 4,940 | 876 | 4,182 | 2.383 |
| *G. biloba* | 41,309 | 8,606 | 32,703 | 12,991 | 1,646 | 7,400 | 876 | 4,008 | 2.517 |
| *G. montanum* | 27,491 | 2,223 | 25,268 | 10,999 | 984 | 4,245 | 876 | 3,464 | 2.297 |
| *P. taeda* | 36,732 | 5,668 | 31,064 | 6,380 | 1,995 | 12,092 | 876 | 4,051 | 4.869 |
| *N. tetragona* | 31,589 | 3,930 | 27,659 | 10,834 | 1,100 | 6,484 | 876 | 3,500 | 2.553 |
| *O. sativa* | 27,694 | 4,856 | 22,838 | 9,883 | 1,188 | 4,445 | 876 | 3,986 | 2.311 |
| *P. abies* | 26,437 | 2,273 | 24,164 | 9,880 | 545 | 1,788 | 876 | 4,466 | 2.446 |
| *P. patens* | 20,328 | 1,273 | 19,055 | 9,155 | 866 | 2,455 | 876 | 3,097 | 2.081 |
| *S. cucullata* | 19,779 | 2,821 | 16,958 | 9,634 | 387 | 1,205 | 876 | 2,641 | 1.76 |
| *S. moellendorffii* | 22,285 | 1,770 | 20,515 | 9,721 | 1,550 | 6,100 | 876 | 2,594 | 2.11 |
| *C. lanceolata* | 37,225 | 2,605 | 34,620 | 13,266 | 797 | 2,982 | 876 | 5,337 | 2.61 |
| *T. wallichiana* | 44,035 | 2,882 | 41,153 | 14,364 | 1,761 | 7,231 | 876 | 4,959 | 2.865 |

#### Supplementary Table15. KEGG enrichment of significant expansion and construction gene families of *C. lanceolata* genome. (see separate file)

#### Supplementary Table 16. GO enrichment of significant expansion gene family in *C. lanceolata* genome. (see separate file)

**Supplementary Table17. The gene ID in Cluser of GO significant enrichment of significant expansion gene families of plants genomes.** (see separate file)

**Supplementary Table 18. Gene information of significantly expanded genes enriched to OG0000016 in *C. lanceolata* genome.** (see separate file)

**Supplementary Table19. Gene information of significantly expanded genes enriched to OG0000039 in *C. lanceolata* genome.** (see separate file)

**Supplementary Table 20. Hypothetical WGDs, posterior mean of duplicate retention rate (*q*), and the Bayes Factor (*K*) to compare the likelihood of *q* = 0 (H_0_) to the likelihood of *q* > 0 (H_1_) using the Savage-Dickey density ratio.**

| Hypotheses | Relaxed branch-specific model | |  | Critical branch-specific model | |
| --- | --- | --- | --- | --- | --- |
|  | $\bar{q}$ | *K* |  | $\bar{q}$ | *K* |
| WGD1 | 0.11233 | 0.25962* |  | 0.23817 | 0.06824** |
| WGD2 | 0.37868 | 0.04426** |  | 0.17672 | 0.09463** |
| WGD3 | 0.08559 | 0.72805 |  | 0.00028 | 2899.97850 |
| WGD4 | 0.03037 | 3.58108 |  | 0.00037 | 3564.65813 |
| WGD5 | 0.59532 | 0.02821*** |  | 0.00332 | 246.95914 |

*K* < 1/102 or *K* < 0.01, decisive evidence against H_0_****; *K* < 1/101.5 or *K* < 0.0316, very strong evidence against H_0_***; *K* < 1/10 or K < 0.1, substantial evidence against H_0_**; *K* < 1/100.5 or *K* < 0. 3162 substantial evidences against H_0_*; K < 1, H_1_ supported, not worth more than a bare mention; *K* > 1, H_0_ supported.

#### Supplementary Table 21. List of MADS-box genes identified in *C. lanceolata* genome.

| **Gene ID** | **Accession number** | **Location** | **Chr** | **ORF (bp)** | **Size (aa)** | **Group** |
| --- | --- | --- | --- | --- | --- | --- |
| Cl10262 | MT103468 | 519587430-519650702 | 1 | 678 | 225 | AG |
| Cl36126 | MT103469 | 526300958-526365655 | 1 | 678 | 225 | AG |
| Cl26543 | MT103470 | 521807977-521978150 | 1 | 756 | 251 | AG |
| Cl26063 | MT103471 | 70041826-70094329 | 11 | 693 | 230 | AG |
| Cl13850 | MT103472 | 70233297-70328225 | 11 | 471 | 156 | AG |
| Cl35439 | MT103473 | 89314620-89335958 | 2 | 666 | 221 | AG |
| Cl35065 | MT103474 | 94642941-94733811 | 2 | 738 | 245 | AGL6 |
| Cl29520 | MT103475 | 366538841-366706949 | 11 | 471 | 156 | AGL6 |
| Cl35039 | MT103477 | 1392696700-1393060149 | 1 | 927 | 308 | TM8 |
| Cl22970 | MT103478 | 1393706469-1394059060 | 1 | 636 | 211 | TM8 |
| Cl12097 | MT103479 | 1392261362-1392431943 | 1 | 636 | 211 | TM8 |
| Cl11956 | MT103480 | 201000156-201216026 | 1 | 636 | 211 | TM8 |
| Cl30647 | MT103481 | 1395412787-1395891794 | 1 | 636 | 211 | TM8 |
| Cl17451 | MT103482 | 426919818-427049392 | 10 | 465 | 154 | TM8 |
| Cl36264 | MT103483 | 98633241-98845766 | 2 | 696 | 231 | TM3 |
| Cl34784 | MT103484 | 596818838-596974165 | 6 | 708 | 235 | SVP |
| Cl35571 | MT103485 | 745221257-745358270 | 3 | 651 | 216 | SVP |
| Cl01446 | MT103486 | 746814647-747037347 | 3 | 720 | 239 | SVP |
| Cl22549 | MT103492 | 505217517-505359167 | 11 | 681 | 226 | GpMADS4 |
| Cl29629 | MT103493 | 502196965-502224894 | 11 | 612 | 203 | GpMADS4 |
| Cl27678 | MT103494 | 291146007-291466416 | 8 | 813 | 270 | GGM13 |
| Cl28889 | MT103495 | 758894530-758896706 | 2 | 846 | 281 | DEF/GLO |
| Cl22633 | MT103496 | 750408487-750411952 | 2 | 588 | 195 | DEF/GLO |
| Cl08169 | MT103497 | 750587944-750592183 | 2 | 621 | 206 | DEF/GLO |
| Cl27804 | MT103498 | 759848871-759850338 | 2 | 525 | 174 | DEF/GLO |
| Cl17023 | MT103499 | 759890589-759892578 | 2 | 726 | 241 | DEF/GLO |
| Cl14691 | MT103500 | 746553907-746558043 | 2 | 642 | 213 | DEF/GLO |
| Cl31654 | MT103501 | 568065106-568070272 | 2 | 1062 | 353 | MIKC* |
| Cl08187 | MT103502 | 685865937-685938418 | 9 | 1152 | 383 | MIKC* |
| Cl07566 | MT103503 | 684871634-684879797 | 9 | 1131 | 376 | MIKC* |
| Cl22501 | MT103504 | 567007448-567046037 | 11 | 1131 | 376 | MIKC* |
| Cl31662 | MT103505 | 687228890-687234262 | 9 | 1146 | 381 | MIKC* |
| Cl15999 | MT103506 | 621864422-621864784 | 4 | 363 | 120 | Mα |
| Cl24955 | MT103507 | 716567113-716567590 | 4 | 318 | 105 | Mα |
| Cl21596 | MT103508 | 4079400-4079645 | 6 | 246 | 81 | Mα |
| Cl00442 | MT103509 | 143546-144199 | / | 654 | 217 | Mα |
| Cl08495 | MT103510 | 718529476-718530006 | 4 | 531 | 176 | Mα |
| Cl21016 | MT103511 | 695835786-695836619 | 1 | 834 | 277 | Mα |
| Cl09032 | MT103512 | 575513819-575514562 | 3 | 744 | 247 | Mα |
| Cl29136 | MT103513 | 574354493-574355266 | 3 | 774 | 257 | Mα |
| Cl07839 | MT103514 | 835739771-835740823 | 3 | 1053 | 350 | Mα |
| Cl07629 | MT103515 | 6402122-6403189 | 8 | 1068 | 355 | Mα |
| Cl30941 | MT103516 | 492704690-492705757 | 10 | 1068 | 355 | Mα |
| Cl01366 | MT103517 | 561794249-561795499 | 3 | 1251 | 416 | Mγ |
| Cl19059 | MT103518 | 979636832-979638091 | 3 | 1260 | 419 | Mγ |
| Cl05223 | MT103519 | 494123266-494124511 | 5 | 1182 | 393 | Mγ |
| Cl18167 | MT103520 | 979988133-979989272 | 3 | 1140 | 379 | Mγ |
| Cl14558 | MT103521 | 531037074-531037772 | 7 | 699 | 232 | Mγ |
| Cl34141 | MT103522 | 770338139-770339431 | 7 | 1293 | 430 | Mγ |
| Cl00625 | MT103523 | 980578572-980579186 | 3 | 615 | 204 | Mγ |

#### Supplementary Table 22. Metabolomic sequencing of *C. lanceolata* astringent seeds at different stages. (see separate file)

#### Supplementary Table 23. List of putative flavonoid synthesis related genes in *C. lanceolata* genome.

| **Abbreviation** | **Gene name** | **Gene ID** |
| --- | --- | --- |
| *PAL* | Phenylalanone ammonia-lyase | Cl16939; Cl23281; Cl20012; Cl15629; Cl01003; Cl10224 |
| *C4H* | Cinnamate 4-hydroxylase | Cl32060; Cl22669; Cl27969 |
| *4CL* | Coumarate-4-CoA ligase | Cl32500; Cl18950; Cl23923; Cl29221; Cl20157; Cl11300; Cl08738; Cl36046; Cl18336; Cl23029; Cl33619; Cl25933; Cl21436; Cl09422; Cl29190; Cl26264; Cl16358; Cl02585; Cl23628; Cl07412; Cl15006; Cl27201; Cl16061; Cl01710; Cl33345; Cl22156; Cl13819; Cl22391; Cl33941; Cl36440; Cl15332; Cl34086; Cl04819; Cl06575; Cl24383 |
| *CHS* | Chalcone synthase | Cl20463; Cl16987; Cl13810; Cl18516; Cl13939; Cl35852; Cl36558; Cl25812; Cl36434; Cl23043; Cl12414; Cl02164 |
| *CHI* | chalcone-flavanone isomerase | Cl23637; Cl26105; Cl35530; Cl01348 |
| *F3H* | flavanone-3-hydroxylase | Cl05230; Cl22835; Cl19511; Cl30186; Cl27251; Cl05548; Cl23387; Cl09496 |
| *F3’H* | flavonoid 3’-hydroxylase | Cl26386; Cl20734; Cl00578 |
| *F3’5’H* | flavonoid 3’,5’-hydroxylase | Cl11848; Cl12701; Cl03636; Cl28266; Cl06640; Cl09151; Cl01764; Cl12036 |
| *FLS* | Flavonol synthase | Cl13876; Cl03153; Cl27668; Cl21034; Cl01929; Cl15880; Cl05506; Cl33239; Cl21939; Cl14727; Cl09073; Cl34583 |
| *DFR* | dihydroflavonols 4-reductase | Cl01163; Cl14835; Cl32921; Cl34319; Cl17992; Cl09745; Cl19190; Cl10044; Cl25990; Cl21164; Cl20763; Cl34597; Cl16924; Cl30627; Cl10196 |
| *LAR* | leucoanthocyanidins reductase | Cl19924; Cl17849; Cl02972; Cl22572; Cl04043; Cl32845; Cl08749; Cl03800; Cl01405; Cl26343 |
| *ANS* | anthocyanidin synthase | Cl28499 |
| *ANR* | anthocyanidin reductase | Cl02683; Cl08429; Cl27585; Cl01531; Cl00030; Cl14400; Cl29306; Cl29276; Cl27034; Cl27658; Cl25902; Cl03085; Cl09863; Cl21775; Cl06423; Cl10344; Cl18261; Cl10859; Cl23026; Cl30270; Cl14318; Cl20814; Cl15108; Cl26250; Cl09989 |

#### Supplementary Table 24. Identified embryo-defective genes (EMBs) in *C. lanceolata* genome. (see separate file)

#### Supplementary Table 25. List of putative *C. lanceolata* shade-related genes.

| **Gene Name** | ***C. lanceolata* Gene ID** | |
| --- | --- | --- |
| Phototropin 1 | | Cl14893 |
| Phototropin 2 | | Cl25517; Cl04142 |
| UV resistance locus 8 (UVR8) | | Cl24527; Cl04477; Cl29114; Cl00787; Cl13109; Cl02421; Cl07100; Cl17449 |
| Cryptochrome (Cry1) | | Cl17630 |
| Cryptochrome (Cry2) | | Cl24639; Cl22470 |
| Phytochrome B (PhyB) | | Cl22557; Cl32891; Cl04073; Cl07369 |
| Constitutive photomor-phogensis 1(COP1) | | Cl21889; Cl14590 |
| PIF3 | | Cl02626; Cl23109 |
| PIF7 | | Cl30879 |
| AUX/IAA | | Cl24287; Cl34491; Cl01117; Cl08639; Cl34992; Cl29820; Cl28099; Cl25274; Cl03095; Cl05774; Cl06592; Cl24063; Cl34660; Cl22617; Cl18037; Cl25125; Cl37123; Cl34659; Cl11114 |

#### Supplementary Table 26. The sequencing quality of the raw data from the nine Hi-C sequencing libraries.

| **Library** | **ReadSum** | **BaseSum** | **GC (%)** | **N (%)** | **Q20 (%)** | **Q30 (%)** |
| --- | --- | --- | --- | --- | --- | --- |
| L01 | 231,458,781 | 69,320,910,642 | 38.33 | 0 | 97.01 | 93.09 |
| L02 | 216,574,783 | 64,867,446,176 | 38.26 | 0.03 | 94.86 | 88.84 |
| L03 | 245,663,762 | 73,581,674,890 | 38.2 | 0.03 | 95.09 | 89.24 |
| L04 | 214,400,226 | 64,223,373,938 | 38.23 | 0.01 | 95.81 | 90.6 |
| L05 | 196,182,529 | 58,763,528,824 | 38.33 | 0.01 | 95.71 | 90.43 |
| L06 | 248,263,233 | 74,357,707,184 | 38.34 | 0 | 97.05 | 93.21 |
| L07 | 274,734,867 | 82,259,170,272 | 38.17 | 0 | 97 | 95.29 |
| L08 | 251,334,974 | 75,261,718,710 | 38.17 | 0 | 96.16 | 94.07 |
| L09 | 254,106,397 | 76,093,916,328 | 38.15 | 0 | 96.76 | 94.94 |
| Total | 2,132,719,552 | 638,729,446,964 | / | / | / | / |

#### Supplementary Table 27. The statistics of the mapped efficiency of the pairs on the nine Hi-C sequencing library data.

| **Library** | **Type** | **Number** | **Ratio (%)** |
| --- | --- | --- | --- |
| L01 | Unique Paired Alignments | 74,149,083 | 100 |
|  | Valid Interaction Pairs | 68,649,601 | 92.58 |
|  | Dangling End Pairs | 3,640,918 | 4.91 |
|  | Re-ligation Pairs | 405,084 | 0.55 |
|  | Self-cycle Pairs | 99,755 | 0.13 |
|  | Dumped Pairs | 1,353,725 | 1.83 |
| L02 | Unique Paired Alignments | 67,062,534 | 100 |
|  | Valid Interaction Pairs | 60,859,580 | 90.75 |
|  | Dangling End Pairs | 4,395,499 | 6.55 |
|  | Re-ligation Pairs | 415,483 | 0.62 |
|  | Self-cycle Pairs | 110,980 | 0.17 |
|  | Dumped Pairs | 1,280,992 | 1.91 |
| L03 | Unique Paired Alignments | 75,942,872 | 100 |
|  | Valid Interaction Pairs | 68,950,679 | 90.79 |
|  | Dangling End Pairs | 4,966,975 | 6.54 |
|  | Re-ligation Pairs | 468,638 | 0.62 |
|  | Self-cycle Pairs | 124,476 | 0.16 |
|  | Dumped Pairs | 1,432,104 | 1.89 |
| L04 | Unique Paired Alignments | 68,132,268 | 100 |
|  | Valid Interaction Pairs | 62,320,281 | 91.47 |
|  | Dangling End Pairs | 3,960,157 | 5.81 |
|  | Re-ligation Pairs | 403 | 0.59 |
|  | Self-cycle Pairs | 116,034 | 0.17 |
|  | Dumped Pairs | 1,333,095 | 1.96 |
| L05 | Unique Paired Alignments | 60,413,157 | 100 |
|  | Valid Interaction Pairs | 54,939,692 | 90.94 |
|  | Dangling End Pairs | 3,799,713 | 6.29 |
|  | Re-ligation Pairs | 371,197 | 0.61 |
|  | Self-cycle Pairs | 92,730 | 0.15 |
|  | Dumped Pairs | 1,209,825 | 2.00 |
| L06 | Unique Paired Alignments | 80,471,301 | 100 |
|  | Valid Interaction Pairs | 74,585,391 | 92.69 |
|  | Dangling End Pairs | 3,831,391 | 4.76 |
|  | Re-ligation Pairs | 480,089 | 0.6 |
|  | Self-cycle Pairs | 87,930 | 0.11 |
|  | Dumped Pairs | 1,486,500 | 1.85 |
| L07 | Unique Paired Alignments | 89,802,324 | 100 |
|  | Valid Interaction Pairs | 82,996,299 | 92.42 |
|  | Dangling End Pairs | 4,306,629 | 4.8 |
|  | Re-ligation Pairs | 475,349 | 0.53 |
|  | Self-cycle Pairs | 146,436 | 0.16 |
|  | Dumped Pairs | 1,877,611 | 2.09 |
| L08 | Unique Paired Alignments | 82,128,266 | 100 |
|  | Valid Interaction Pairs | 76,225,099 | 92.81 |
|  | Dangling End Pairs | 3,705,017 | 4.51 |
|  | Re-ligation Pairs | 481,714 | 0.59 |
|  | Self-cycle Pairs | 86,016 | 0.1 |
|  | Dumped Pairs | 1,630,420 | 1.99 |
| L09 | Unique Paired Alignments | 84,783,347 | 100 |
|  | Valid Interaction Pairs | 79,165,448 | 93.37 |
|  | Dangling End Pairs | 3,374,676 | 3.98 |
|  | Re-ligation Pairs | 474,558 | 0.56 |
|  | Self-cycle Pairs | 76,547 | 0.09 |
|  | Dumped Pairs | 1,692,118 | 2.0 |

Unique Paired Alignments: The only read pairs aligned to the assembled genome; Valid Interaction Pairs: Valid Read Pairs; Dangling End Pairs: Data of end suspension type in invalid read pairs; Re-ligation Pairs: Dara of adjacent connection type in invalid read pairs; Self-circle Ligation Pairs: The invalid data belongs to read pairs of self-connecting type; Dumped Pairs: The invalid data belongs to other undefined read pairs.

**Supplementary Table 28. Interaction combination of aluminum and astringent seed water extract treatment.**

| Factors/treatments | WN | W10 | W0 |
| --- | --- | --- | --- |
| Al0 | Al0WN | Al0W10 | Al0W0 |
| Al1 | Al1WN | Al1W10 | Al1W0 |
| Al2 | Al2WN | Al2W10 | Al2W0 |

### Supplementary Reference

1. Woodward, A.W. & Bartel, B. Auxin: regulation, action, and interaction. *Ann. Bot.* **95**, 707–735 (2005).

2. Singh, V. K. *et al*. Genome-wide survey and comprehensive expression profiling of Aux/IAA gene family in chickpea and soybean. *Front. Plant Sci.* **6**, 918 (2015).

3. Casal, J. J. Photoreceptor signaling networks in plant responses to shade. *Annu. Rev. Plant Biol.* **64**, 403–427 (2013)

4. Remington, D. L. *et al*. Contrasting modes of diversification in the Aux/IAA and ARF gene families. *Plant Physiol.* **135**, 1738–1752 (2004).

5. Guan, R. *et al.* Draft genome of the living fossil *Ginkgo biloba*. *GigaScience* **5**, 49 (2016)

6. Wan, T. *et al*. A genome for gnetophytes and early evolution of seed plants. *Nat. Plants* **4**, 82–89 (2018).

7. Nystedt, B. *et al*. The *Norway spruce* genome sequence and conifer genome evolution. *Nature* **497**, 579–584 (2013).

8. Amborella Genome Project. The *Amborella* genome and the evolution of flowering plants. *Science* **342**,1241089 (2013).

9. Birol, I. *et al.* Assembling the 20 Gb white spruce (*Picea glauca*) genome from whole-genome shotgun sequencing data. *Bioinformatics* **29**, 1492–1497 (2013).

10. Xiong, X. *et al.* The *Taxus* genome provides insights into paclitaxel biosynthesis. *Nat. Plants* **7**, 1026–1036 (2021).
